# A novel family of secreted insect proteins linked to plant gall development

**DOI:** 10.1101/2020.10.28.359562

**Authors:** Aishwarya Korgaonkar, Clair Han, Andrew L. Lemire, Igor Siwanowicz, Djawed Bennouna, Rachel Kopec, Peter Andolfatto, Shuji Shigenobu, David L. Stern

## Abstract

In an elaborate form of inter-species exploitation, many insects hijack plant development to induce novel plant organs called galls that provide the insect with a source of nutrition and a temporary home. Galls result from dramatic reprogramming of plant cell biology driven by insect molecules, but the roles of specific insect molecules in gall development have not yet been determined. Here we study the aphid *Hormaphis cornu*, which makes distinctive “cone” galls on leaves of witch hazel *Hamamelis virginiana*. We found that derived genetic variants in the aphid gene *determinant of gall color* (*dgc*) are associated with strong downregulation of *dgc* transcription in aphid salivary glands, upregulation in galls of seven genes involved in anthocyanin synthesis, and deposition of two red anthocyanins in galls. We hypothesize that aphids inject DGC protein into galls, and that this results in differential expression of a small number of plant genes. *Dgc* is a member of a large, diverse family of novel predicted secreted proteins characterized by a pair of widely spaced cysteine-tyrosine-cysteine (CYC) residues, which we named BICYCLE proteins. *Bicycle* genes are most strongly expressed in the salivary glands specifically of galling aphid generations, suggesting that they may regulate many aspects of gall development. *Bicycle* genes have experienced unusually frequent diversifying selection, consistent with their potential role controlling gall development in a molecular arms race between aphids and their host plants.

**One Sentence Summary:** Aphid *bicycle* genes, which encode diverse secreted proteins, contribute to plant gall development.

## Main Text

Organisms often exploit individuals of other species, for example through predation or parasitism. Parasites sometimes utilize molecular weapons against hosts, which themselves respond with molecular defenses, and the genes that encode or synthesize these molecular weapons may evolve rapidly in a continuous ‘arms race’ (Obbard et al., 2006; Papkou et al., 2019; Paterson et al., 2010). Some of the most elaborate molecular defenses—such as adaptive immune systems, restriction modification systems, and CRISPR—have resulted from such host-parasite conflicts. In many less well-studied systems, parasites not only extract nutrients from their hosts but they also alter host behavior, physiology, or development to the parasite’s advantage (Heil, 2016). Insect galls represent one of the most extreme forms of such inter-species manipulation.

Insect-induced galls are intricately patterned homes that provide insects with protection from environmental vicissitude and from some predators and parasites (Bailey et al., 2009; Mani, 1964; Stone and Schönrogge, 2003). Galls are also resource sinks, drawing nutrients from distant plant organs, and providing insects with abundant food (Larson and Whitham, 1991). Insect galls are atypical plant growths that do not result simply from unpatterned cellular over-proliferation, as observed for microbial galls like the crown gall induced by *Agrobacterium tumefaciens*. Instead, each galling insect species appears to induce a distinctive gall, even when related insect species attack the same plant, implying that each species provides unique instructions to re-program latent plant developmental networks (Cook and Gullan, 2008; Crespi and Worobey, 1998; Dodson, 1991; Hearn et al., 2019; Leatherdale, 1955; Martin, 1938; Martinson et al., 2020; Parr, 1940; Plumb, 1953; Stern, 1995; Stone and Cook, 1998).

At least some gall-inducing insects produce phytohormones (Dorchin et al., 2009; McCalla et al., 1962; Suzuki et al., 2014; Tanaka et al., 2013; Tooker and Helms, 2014; Yamaguchi et al., 2012), although it is not yet clear whether insects introduce these hormones into plants to support gall development. However, injection of phytohormones alone probably cannot generate the large diversity of species-specific insect galls. In addition, galling insects induce plant transcriptional changes independently of phytohormone activity (Bailey et al., 2015; Hearn et al., 2019; Nabity et al., 2013; Papkou et al., 2019; Shih et al., 2018; Takeda et al., 2019). Thus, given the complex cellular changes required for gall development, insects probably introduce many molecules into plant tissue to induce galls.

In addition to the potential role of phytohormones in promoting gall growth, candidate gall effectors have been identified in several gall-forming insects (Cambier et al., 2019; Eitle et al., 2019; Zhao et al., 2015). However, none of these candidate effectors have yet been shown to contribute to gall development or physiology. In addition, while many herbivorous insects introduce effector molecules into plants to influence plant physiology (Elzinga and Jander, 2013; Hogenhout and Bos, 2011; Kaloshian and Walling, 2016; Stuart, 2015), there is no evidence that any previously described effectors contribute to gall development. Since there are currently no galling insect model systems that would facilitate a genetic approach to this problem, we turned to natural variation to identify insect genes that contribute to gall development.

We studied the aphid, *Hormaphis cornu*, which induces galls on the leaves of witch hazel, *Hamamelis virginiana*, in the Eastern United States (Fig. 1A-F). In early spring, each *H. cornu* gall foundress (fundatrix) probes an expanding leaf with her microscopic mouthparts (stylets) (Fig. 1A, B, Video S1) and pierces individual mesophyll cells with her stylets (Fig. 1G and H) (Lewis and Walton, 1947, 1958). We found that plant cells near injection sites, revealed by the persistent stylet sheaths, over-proliferate through periclinal cell divisions (Figure 1H). This pattern of cytokinesis is not otherwise observed in leaves at this stage of development (Figure 1I) and contributes to the thickening and expansion of leaf tissue that generates the gall (Fig. 1D-G). The increased proliferation of cells near the tips of stylet sheaths suggests that secreted effector molecules produced in the salivary glands are deposited into the plant via the stylets.

**Fig. 1.**
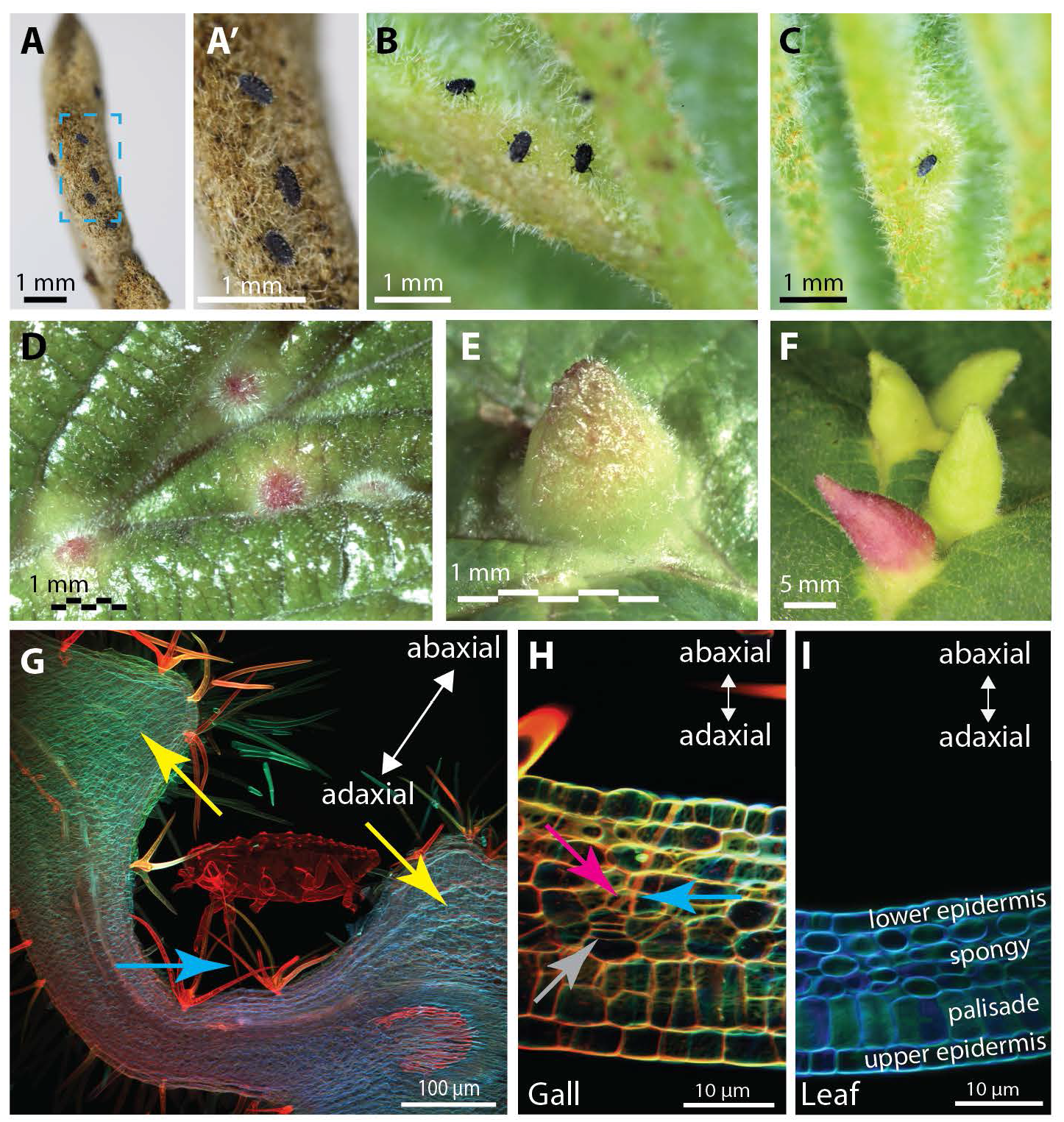
*Hormaphis cornu* aphids drive patterned over-proliferation of plant cells to produce galls of on leaves *Hamamelis virginiana*. (A-C) Photographs of the abaxial surfaces of *H. virginiana* leaves being attacked by first-instar fundatrices of *H. cornu*. Nymphs gather on the unopened leaf buds (A) and soon after bud break the fundatrices gather near leaf veins (B) and inject material that begins to transform the leaf into a gall (C). Magnified region in blue rectangle of panel (A) shows three fundatrices waiting on unopened bud (A’). (D-F) Photographs of the adaxial leaves of *H. virginiana*, showing galls at early (D), middle (E), and late (F) growth stages. (G-I) Confocal micrographs of sections through a *H. cornu* gall (G,H) and un-galled *H. virginiana* leaf (I) stained with Congo red and calcofluor white. Extensive hypertropy is observed in the mesophyll (yellow arrows) at a considerable distance from the location of the aphid’s stylets (blue arrows) (G). The tips of aphid stylets (pink arrow) can be observed within cells of *H. virginiana* and plant tissue shows evidence of hyperproliferation and periclinal divisions (grey arrows) close to the tips of stylets and the termini of stylet sheaths (H). Periclinal divisions are observed in both spongy and palisade mesophyll cells during gall development, but never in ungalled leaf tissue (I). Plant tissue is presented in the aphid’s frame of reference, with abaxial leaf surface up.

After several days, the basal side of the gall encloses the fundatrix and the gall continues to grow apically and laterally, providing the fundatrix and her offspring with protection and abundant food. After several weeks, the basal side of the gall opens to allow aphids to remove excreta (honeydew) and molted nymphal skins from the gall and, eventually, to allow winged migrants to depart. Continued gall growth requires the constant presence of the fundatrix and gall tissue dies in her absence (Lewis and Walton, 1958; Rehill and Schultz, 2001), suggesting that the fundatrix must continuously inject salivary-gland produced effectors to overcome plant defenses.

### A natural gall color polymorphism is linked to regulatory variation in a novel aphid gene, *determinant of gall color*

We found that populations of *H. cornu* include approximately 4% red galls and 96% green galls (Fig. 1F). We inferred that this gall color polymorphism results from differences among aphids, rather than from differences associated with leaves or the location of galls on leaves, because red and green galls are located randomly on leaves and often adjacent to each other on a single leaf (Fig. 1F). We sequenced and annotated the genome of *H. cornu* (Supplemental Material) and performed a genome-wide association study (GWAS) on fundatrices isolated from 43 green galls and 47 red galls by resequencing their genomes to approximately 3X coverage. There is no evidence for genome-wide differentiation of samples from red and green galls, suggesting that individuals making red and green galls were sampled from a single interbreeding population (Fig. S1A-D). We identified SNPs near 40.5 Mbp on Chromosome 1 that were strongly associated with gall color (Fig. 2A). We re-sequenced approximately 800 kbp flanking these SNPs to approximately 60X coverage and identified 11 single-nucleotide polymorphisms (SNPs) within the introns and upstream of gene *Horco_16073* that were strongly associated with gall color (Fig. 2B-D). There is no evidence that large scale chromosomal aberrations are associated with gall color (Supplementary Text).

**Figure 2.**
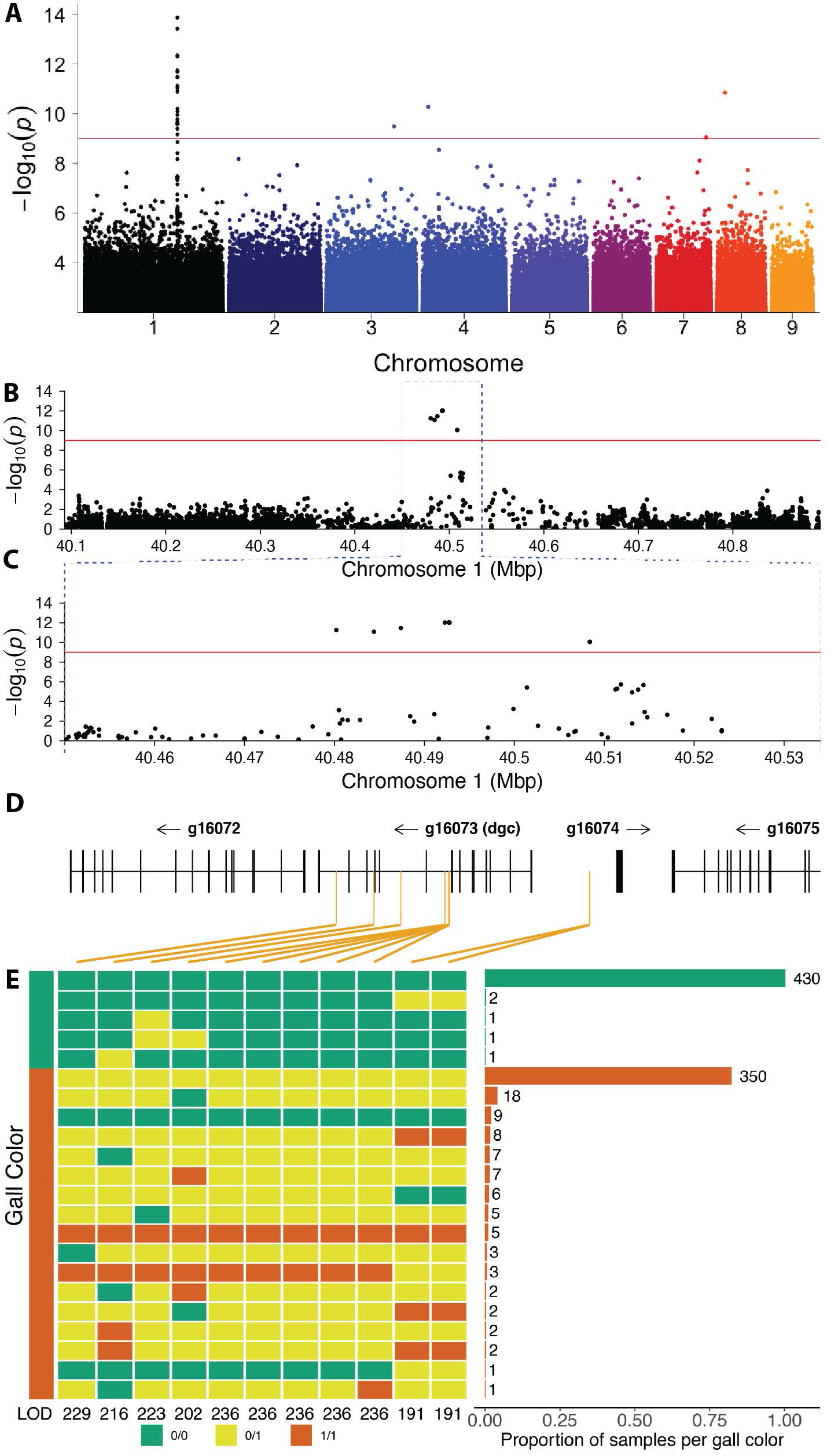
A genome-wide association study (GWAS) identifies variation within a novel aphid gene associated with gall color. (A) A GWAS of fundatrices isolated from 43 green and 47 red galls identified a small region on chromosome 1 of the *H. cornu* genome that is strongly associated with gall color. Red line indicates FDR = 0.05. Colors of points on chromosomes are arbitrary. (B-D) Resequencing 800 kbp of Chromosome 1 centered on the most significant SNPs from the original GWAS to approximately 60X coverage identified 11 spatially clustered SNPs significantly associated with gall color located within the introns and upstream of *Horco_g16073*, which was named *dgc* (D). (Some SNPs are closely adjacent and cannot be differentiated at this scale.) Significant SNPs are indicated with orange vertical lines in (D). (E) Genotypes of all 11 SNPs associated with gall color from an independent sample of aphids from 435 green and 431 red galls. Color of gall for aphid samples shown on left and genotype at each SNP is shown adjacent in green (0/0, homozygous ancestral state), yellow (0/1, heterozygous), or red (1/1, homozygous derived state). LOD scores for association with gall color shown for each SNP at bottom of genotype plot. Histogram of frequencies of each multilocus genotype ordered by frequency and collected within gall color is shown on the right. All 11 SNPs are strongly associated with gall color (P < 10^-192^), and a cluster of 5 SNPs in a 61bp region are most strongly associated with gall color. Individuals homozygous for ancestral alleles at all or most loci and making red galls likely carry variants at other loci that influence gall color (Supplementary Text).

Since GWAS can sometimes produce spurious associations, we performed an independent replication study and found that all 11 SNPs were highly significantly associated with gall color in fundatrices isolated from 435 green and 431 red galls (LOD = 191 – 236; Fig. 2E). All fundatrices from green galls were homozygous for the ancestral allele at 9 or more of these SNPs (Fig. 2E). In contrast, 98% of fundatrices from red galls were heterozygous or homozygous for derived alleles at 9 or more SNPs (Fig. 2E). This pattern suggests that alleles contributing to red gall color are genetically dominant to alleles that generate green galls. Two percent of fundatrices that induce red galls were homozygous for ancestral alleles at these SNPs and likely carry genetic variants elsewhere in the genome that confer red color to galls (Supplementary Text).

Based on these genetic associations and further evidence presented below, we assigned the name *determinant of gall color* (*dgc*) to *Horco_16073*. *Dgc* encodes a predicted protein of 23 kDa with an N-terminal secretion signal sequence (Fig. S4). The putatively secreted portion of the protein shares no detectable sequence homology with any previously reported proteins.

Most SNPs associated with green or red galls were found in one of two predominant haplotypes (Fig. 2E) and exhibited strong linkage disequilibrium (LD) (Fig. S5). LD can result from suppressed recombination. However, these 11 SNPs are in linkage equilibrium with many other intervening and flanking SNPs (Fig. S5, S6). Also, multiple observed genotypes are consistent with recombination between these 11 SNPs (Fig. 2E) and we found no evidence for chromosomal aberrations that could suppress recombination (Fig. S1E-I; Supplementary Text). Thus, LD among the 11 *dgc* SNPs associated with gall color cannot be explained by suppressed recombination. It is more likely that the non-random association of the 11 *dgc*^Red^ SNPs has been maintained by natural selection, suggesting that the combined action of all 11 SNPs may have a stronger effect on gene function than any single SNP alone.

### Regulatory variants at *dgc* dominantly silence *dgc* expression

Since all 11 *dgc* polymorphisms associated with gall color occur outside of *dgc* exons (Fig. 2D), we tested whether these polymorphisms influence expression of *dgc* or of any other genes in the genome. We first determined that *dgc* is expressed highly and specifically in fundatrix salivary glands and lowly or not at all in other tissues or other life cycle stages (Fig. S7A). We then performed RNAseq on salivary glands from fundatrices with *dgc*^Green^ /*dgc*^Green^ or *dgc*^Red^/*dgc*^Green^ genotypes. *Dgc* stands out as the most strongly differentially expressed gene between these genotypes (Figure 3A and S7B). Since *dgc*^Red^ alleles appeared to be dominant to the *dgc*^Green^ alleles for gall color, we expected that *dgc* transcripts would be upregulated in animals with *dgc*^Red^ alleles. In contrast, *dgc* transcripts were almost absent in fundatrices carrying *dgc*^Red^ alleles (Fig. 3B). That is, red galls are associated with strongly reduced *dgc* expression in fundatrix salivary glands.

**Figure 3.**
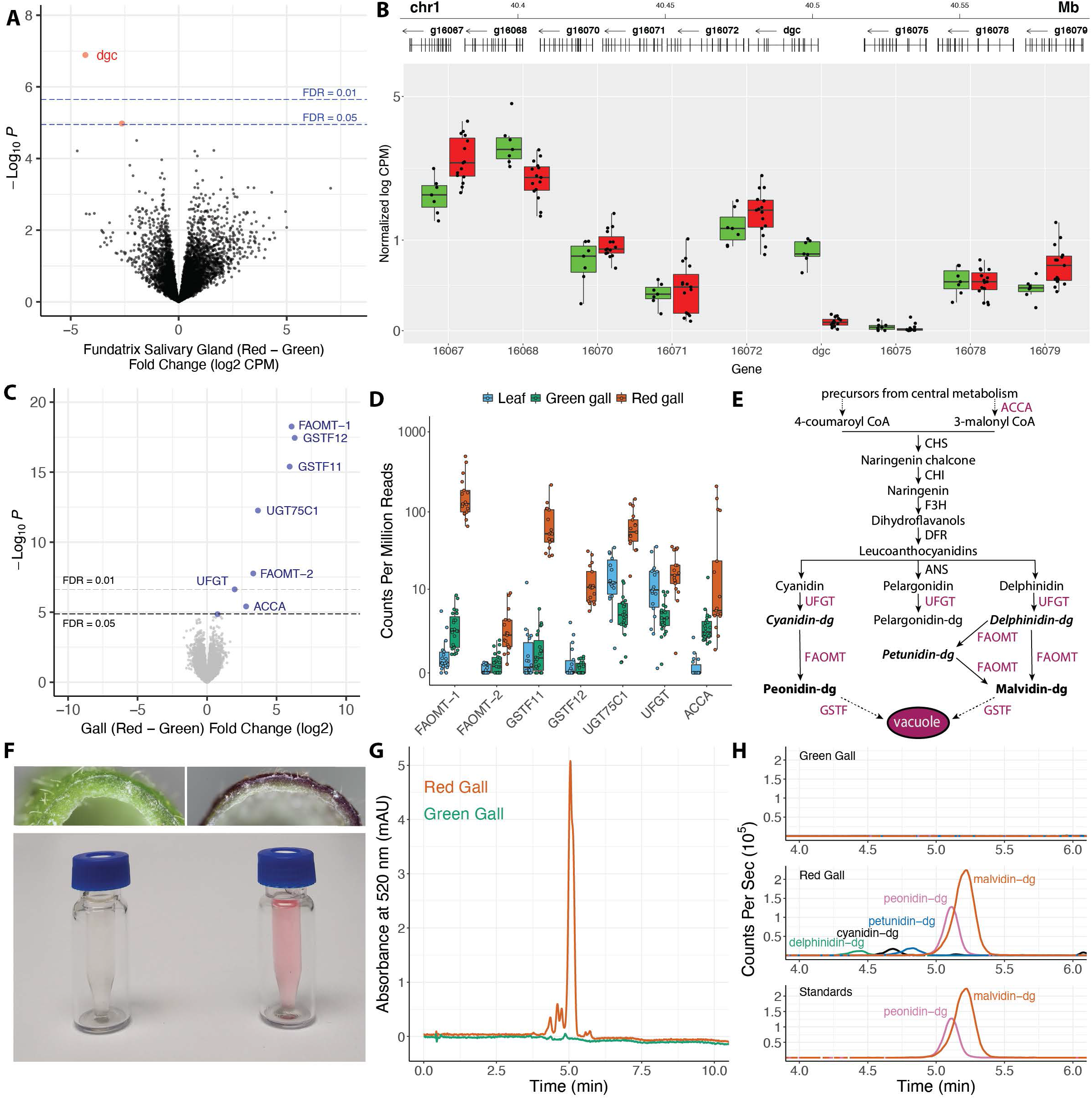
*Dgc* is the most strongly differentially expressed gene in salivary glands of fundatrices collected from red versus green galls. (A) Genome-wide differential expression analysis of *H. cornu* fundatrix salivary gland transcripts from individuals heterozygous for *dgc*^Red^/*dgc*^Green^ (Red; N = 15) versus homozygous for *dgc*^Green^ (Green; N = 7) illustrated as a volcano plot. D*gc* is strongly downregulated in *dgc*^Red^/*dgc*^Green^ fundatrices. (B) Salivary gland expression of genes in the paralog cluster that includes *dgc*^Red^/*dgc*^Green^. Gene models of the paralogous genes are shown at the top and expression levels normalized by mean expression across all paralogs is shown below. Only *dgc* is strongly downregulated in this gene cluster between individuals carrying *dgc*^Red^/*dgc*^Green^ (Red) versus *dgc*^Green^/*dgc*^Green^ (Green) genotypes. (C) Genome-wide differential expression analysis of *H. virginiana* transcripts isolated from galls made by fundatrices heterozygous for *dgc*^Red^/*dgc*^Green^ (Red; N = 17) versus homozygous for *dgc*^Green^ (Green; N = 23) illustrated as a volcano plot. Only eight genes are differentially expressed at FDR < 0.05, and all are overexpressed in red galls. The seven most strongly differentially expressed genes that encode anthocyanin biosynthetic enzymes (FAOMT-1 = Hamvi_23591; FAOMT-2 = Hamvi_7147; GSTF11 = Hamvi_134919; GSTF12 = Hamvi_109682; UGT75C1 = Hamvi_14194; UFGT = Hamvi_22774; ACCA = Hamvi_97071). (D) Expression levels, in counts per million reads, of the seven anthocyanin biosynthetic genes overexpressed in red galls, in green (green) and red (red) galls and ungalled leaves (blue). Each data point is from a separate genome-wide RNA-seq sample. (E) Simplified diagram of the anthocyanin biosynthetic pathway. Enzyme classes upregulated in red galls are shown in purple font. The two terminal anthocyanins that generate the red color in galls, peonidin-3,5-diglucoside and malvidin-3,5-diglucoside, are shown in bold font, and three precursor molecules found in red galls are shown in bold italic font. Anthocyanin names are abbreviated (dg = 3,5-diglucoside). (F) Photos of cross-sections of green (top left) and red (top right) and the pigments extracted from green and red galls (below). (G) UHPLC-DAD chromatograms at 520 nm of extract from red (red line) and green (green line) galls. (H) Overlaid UHPLC-MS chromatograms of green (top) and red (middle) gall extracts and authentic standards (bottom). Each pigment is indicated with a different color: green = delphinidin-3,5-diglucoside (*m/z* = 627.1551); black = cyanidin-3,5-diglucoside (*m/z* = 611.1602); blue = petunidin-3,5-diglucoside (*m/z* = 641.1709); purple = peonidin-3,5-diglucoside (*m/z* = 625.1768); and red = malvidin-3,5-diglucoside (*m/z* = 655.1870). Anthocyanin names are abbreviated (dg = 3,5-diglucoside). Peonidin-3,5-diglucoside and malvidin-3,5-diglucoside together account for 87% of pigment detected in red galls.

*Dgc* expression is reduced approximately 20-fold in fundatrix salivary glands with *dgc*^Red^/*dgc*^Green^ (27 ± 22.6 CPM, mean ± SD) versus *dgc*^Green^/*dgc*^Green^ genotypes (536 ± 352.3 CPM, mean ± SD). This result suggested that *dgc*^Red^ alleles downregulate both the *dgc*^Red^ and *dgc*^Green^ alleles in heterozygotes. To confirm whether the *dgc*^Red^ allele downregulates the *dgc*^Green^ allele in *trans*, we identified exonic SNPs that were specific to each allele and could be identified in the RNAseq data. We found that both *dgc*^Red^ and *dgc*^Green^ alleles were strongly downregulated in heterozygotes, confirming the *trans* activity of the *dgc*^Red^ allele (Fig. S7D). We observed no systematic transcriptional changes in neighboring genes (Figure 3B), most of which are *dgc* paralogs, indicating that *dgc*^Red^ alleles exhibit a perhaps unique example of locus-specific repressive transvection (Duncan, 2002).

### High levels of *dgc* transcription are associated with downregulation specifically of plant anthocyanin genes and two anthocyanins

Since red galls are associated with strong differential expression of only *dgc*, we wondered how the plant responds to changes in this single putative effector. To examine this question, we sequenced and annotated the genome of the host plant *Hamamelis virginiana* and then performed whole-genome differential expression on plant mRNA isolated from galls induced by aphids with *dgc*^Red^/*dgc*^Green^ versus *dgc*^Green^/*dgc*^Green^ genotypes (Supplementary Text). Only eight plant genes were differentially expressed between green and red galls and all eight genes were downregulated in green galls (Fig. 3C-D). That is, high levels of *dgc* are associated with downregulation of only eight plant genes in galls.

Red pigmented galls could result from production of carotenoids (Smits and Peterson, 1942), anthocyanins (Blunden and Challen, 1965; Bomfim et al., 2019), or betacyanins. However, in red galls induced by *H.cornu*, the seven most strongly downregulated plant genes are all homologous to genes annotated as enzymes of the anthocyanin biosynthetic pathway (Fig. 3E). One gene encodes an enzyme (ACCA) that irreversibly converts acetyl-CoA to malonyl CoA, a biosynthetic precursor of multiple anthocyanins. Two genes encode anthocyanidin 3-0-glucosyltransferases (UFGT and UGT75C1), which glycosylate unstable anthocyanidins to allow their accumulation (Springob et al., 2003). Two genes encode flavonoid 3’-5’ methyltransferases (FAOMT-1, FAOMT-2), which methylate anthocyanin derivatives (Hugueney et al., 2009). Finally, two genes encode phi class glutathione S-transferases (GSTF11, GSTF12), which conjugate glutathione to anthocyanins, facilitating anthocyanin transport and stable accumulation in vacuoles (Marrs et al., 1995).

Six of the enzymes upregulated in red galls are required for final steps of anthocyanin production and deposition (Fig. 3E) and their upregulation in red galls may account for the accumulation of pigments in red galls. To test this hypothesis, we extracted and analyzed pigments from galls (Fig. 3F) and identified high levels of two pigments only in red galls (Fig. 3G), the anthocyanins malvidin-3,5-diglucoside and peonidin-3,5-diglucoside (Fig. 3G-H, S8). Thus, the pigments present in red galls are products of enzymes in the anthocyanin biosynthetic pathway, such as those encoded by genes that are upregulated in red galls. The two abundant anthocyanins are produced from distinct intermediate precursor molecules (Fig. 3E), three of which were also detected in red galls (Fig. 3H, S8), and synthesis of these two anthocyanins likely requires activity of different methyltransferases and glucosyltransferases. The three pairs of glucosyltransferases, methyltransferases, and glutathione transferases upregulated in red galls may provide the specific activities required for production of these two anthocyanins.

Taken together, these observations suggest that *dgc* represses transcription of seven anthocyanin biosynthetic enzymes. It is not clear how *dgc* induces specific transcriptional changes in seven plant genes; it may act by altering activity of an upstream regulator of these plant genes.

### Aphids induce widespread transcriptomic changes in galls

Gall color represents only one aspect of the gall phenotype, apparently mediated by changes in expression of seven plant genes, and the full complement of cell biological events during gall development presumably requires changes in many more plant genes. To estimate how many plant genes are differentially expressed during development of the *H. cornu* gall on *H. virginiana*, we performed differential expression analysis of plant genes in galls versus the surrounding leaf tissue. Approximately 31% of plant genes were upregulated and 34% were downregulated at FDR = 0.05 in galls versus leaf tissue (Figure S9E; 27% up and 29% down in gall at FDR = 0.01). Results of gene ontology analysis of up and down-regulated genes is consistent with the extensive growth of gall tissue and down regulation of chloroplasts seen in aphid galls (Fig. S9F; Supplementary Text), a pattern observed in other galling systems (Hearn et al., 2019).

Thus, approximately 15,000 plant genes are differentially expressed in galls, representing a system-wide re-programming of plant cell biology. If other aphid effector molecules act in ways similar to *dgc*, which is associated with differential expression of only eight plant genes, then gall development may require injection of hundreds or thousands of effector molecules.

### *Dgc* is a member of a large class of novel *bicycle* genes expressed specifically in the salivary glands of gall-inducing aphids

To identify additional proteins that aphids may inject into plants to contribute to gall development, we exploited the fact that only some individuals in the complex life cycle of *H. cornu* induce galls (Figure 4A). Only the fundatrix generation induces galls and only her immediate offspring live alongside her in the developing gall. In contrast, individuals of generations that live on river birch (*Betula nigra*) through the summer and the sexual generation that feed on *H. virginiana* leaves in the autumn do not induce any leaf malformations. Thus, probably only the salivary glands of the generations that induce galls (the fundatrix (G1) and possibly also her immediate offspring (G2)) produce gall-effector molecules. We identified 3,048 genes upregulated in fundatrix salivary glands versus the fundatrix body (Fig. S10A) and 3,427 genes upregulated in salivary glands of fundatrices, which induce galls, versus sexuals, which do not induce galls although they feed on the same host plant (Fig. S10B). Intersection of these gene sets identified 1,482 genes specifically enriched in the salivary glands of fundatrices (Figure 4B).

**Figure 4.**
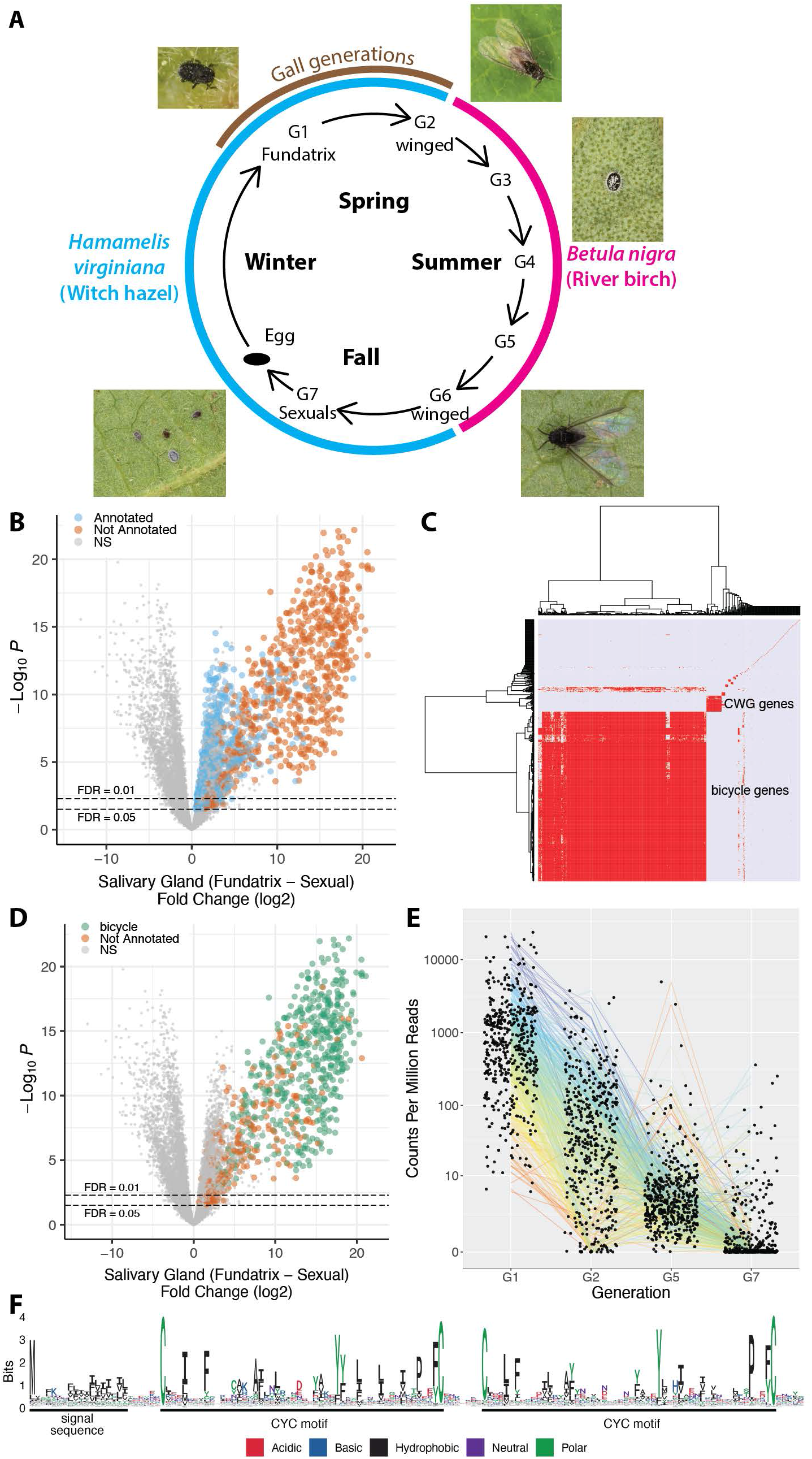
*bicycle* genes are salivary gland enriched transcripts of gall-associated *H. cornu* generations. (A) Diagram of life cycle of *H. cornu*. *H. cornu* migrates annually between *H. virginiana* (blue line) and *Betula nigra* (pink line) and the gall is produced only in the spring on *H. virginiana* (brown line). Each nymph of the first generation (G1, the fundatrix) hatches from an over-wintering egg and initiates development of a single gall. Her offspring (G2) feed and grow within the gall and develop with wings, which allows them to fly to *B. nigra* in late spring. For three subsequent generations (G3-G5) the aphids develop as small, coccidiform morphs on *B. nigra*. In the fall, aphids develop with wings (G6), fly to *H. virginiana* plants, and deposit male and female sexuals (G7), the only generation possessing meiotic cells. The sexuals feed and complete development on the senescing leaves of *H. virginiana*. As adults they mate and the females deposit eggs that overwinter and give rise to fundatrices the following spring. (B) Differential expression of fundatrix versus sexual salivary glands with only genes significantly upregulated in fundatrix salivary glands marked with colors, shown as a volcano plot. Genes with and without homologs in public databases are labeled as “Annotated” (blue) and “Not Annotated” (brown), respectively. (C) Hierarchical clustering of unannotated salivary-gland specific genes reveals one large cluster of *bicycle* genes, one small cluster of *CWG* genes, and many largely unique, unclustered genes. (D) *Bicycle* (green) and remaining unannotated (brown) genes labelled on a differential expression volcano plot illustrates that, on average, *bicycle* genes are the most strongly differentially expressed genes expressed specifically in fundatrix salivary glands. (E) *Bicycle* genes are expressed at highest levels in the salivary glands of the fundatrix and mostly decline in expression during subsequent generations. (Sample sizes: G1 (N = 20); G2 (N = 4); G5 (N = 6); and G7 (N = 6). (F) Amino-acid logo for predicted BICYCLE proteins. Sequence alignment used for logo was filtered to highlight conserved positions. Most genes encode proteins with an N-terminal signal sequence and a pair of conserved cysteine-tyrosine-cysteine motifs (CYC).

Half of these genes (744) were homologous to previously identified genes, many of which had functional annotations. Gene Ontology analysis of the “annotated” genes suggests that they contribute mostly to the demands for high levels of protein secretion in fundatrix salivary glands (Fig. S10C). Most do not encode proteins with secretion signals (671; Fig. S10D) and are thus unlikely to be injected into plants. We searched for homologs of genes that have been proposed as candidate gall-effector genes in other insects but found no evidence that these classes of genes contribute to aphid gall development (Supplementary Text). We therefore focused on the remaining 738 unannotated genes, which included 459 genes encoding proteins with predicted secretion signals (Fig. S10D).

Hierarchical clustering of the unannotated genes by sequence similarity identified one large (476 genes) and one small (43 genes) cluster of related genes, and 222 genes sharing few or no homologs (Fig. 4C). Genes in both the large and small clusters encode proteins with N-terminal secretion signals, as expected for effector proteins that might be injected into plants. The small cluster encodes a divergent set of proteins containing several conserved cysteines (C) and a well conserved tryptophan (W) and glycine (G), and we named these CWG genes (Fig. S11A).

Proteins encoded by the large cluster display conservation mainly of a pair of widely spaced cysteine-tyrosine-cysteine (CYC) motifs and spacing between the C, Y, and C residues of each motif is not well conserved (Fig. 4F). This pair of CYC motifs led us to name these *bicycle* (bi-CYC-like) genes. The *bicycle* genes were the most strongly upregulated class of genes in fundatrix salivary glands (Fig. 4D, Fig. S11B-F) and were expressed specifically in the salivary glands of the two generations associated with galls (G1 and G2) (Fig. 4E). Many *bicycle* genes are found in paralog clusters throughout the *H. cornu* genome (Fig. S10E and F) and each *bicycle* gene contains approximately 5-25 microexons (Fig. S10F)—a large excess relative to the genomic background (Fig. S10G)—interrupted by long introns (Fig. S10F).

We found that *dgc* shares many features with *bicycle* genes—it is strongly expressed specifically in fundatrix salivary glands, and it exhibits many microexons and a pair of CYC motifs (Fig. S4)—and that it is evolutionarily related to other *bicycle* genes. Thus, *dgc* is a member of a diverse family of genes encoding secreted proteins expressed specifically in the salivary glands of gall forming generations. *Bicycle* genes are therefore good candidates to encode many of the molecules required to generate the extensive transcriptional changes observed in galls.

### *bicycle* genes experienced intense diversifying selection, consistent with a potential arms race between aphids and plants

*Bicycle* genes are extremely diverse at the amino acid sequence level, as has been observed for other candidate insect gall effector genes (Chen et al., 2010). Each *bicycle* protein has accumulated approximately two substitutions per amino acid site since divergence from paralogs (Fig. S10H). To explore whether this diversity resulted from natural selection rather than genetic drift, we compared rates of non-synonymous (*d*_N_) versus synonymous (*d*_S_) substitutions between the sister species *H. cornu* and *H. hamamelidis* (which also induces galls) in *bicycle* versus non-*bicycle* genes, because *d*_N_/*d*_S_ values greater than one provide evidence for positive selection (Yang and Bielawski, 2000). To calculate polymorphism of orthologous genes in each species, we mapped sequencing reads from individuals of each species to the *H. cornu* genome. For divergence estimates, we estimated the *H. hamamelidis* genome by mapping *H. hamamelidis* sequencing reads to the *H. cornu* genome (Supplemental Methods) and compared this genome with the original *H. cornu* genome. A large excess of *bicycle* genes displayed *d*_N_/*d*_S_ significantly greater than 1 relative to the genomic background (Fig. S12, Table S1; P < 2.2e-16), revealing recurrent adaptive amino acid substitution at many *bicycle* genes since these species diverged.

We then quantified the frequency of adaptive amino acid substitutions at *bicycle* genes and other categories of genes overexpressed in fundatrix salivary glands by calculating *α*, the proportion of non-synonymous substitutions fixed by positive selection (Fay et al., 2001; Smith and Eyre-Walker, 2002). We estimate that for all *bicycle* genes *α* = 0.33 (95%CI 0.24 - 0.41) and that for the subset of *bicycle* genes displaying *d*_N_/*d*_S_ significantly greater than 1, *α* = 0.62 (95%CI 0.45-0.73, Table S2). Other categories of genes overexpressed in fundatrix salivary glands display values of *α* in the range of 0.27-0.38, however, *bicycle* genes display a considerably higher ratio of non-synonymous to synonymous substitutions than other categories of genes (Table S2). Most strikingly, as a fraction of protein length, *bicycle* genes display a considerably greater fraction of adaptive amino acid substitutions than other categories of genes (Fig. S12C). Since speciation between *H. cornu* and *H. hamamelidis*, positive selection has resulted, on average, in fixation of approximately 2-3 substitutions in each *bicycle* gene, and approximately 10 substitutions in the most rapidly evolving *bicycle* genes. This represents a considerable fraction of the average length of these proteins (∼200 residues) and reveals intense selection on *bicycle* genes, presumably for novel functions that require multiple amino acid substitutions.

The previous tests cannot detect selection on non-coding regions and do not discriminate between selection acting in the deep past versus more recently. To search for recent selection in *bicycle* gene regions, we examined patterns of polymorphism and divergence within and between *H. cornu* and *H. hamamelidis*. Polymorphism was strongly reduced relative to divergence in *bicycle* gene regions compared with the genomic background (Fig. S13, S14) and patterns of reduced polymorphism were strikingly similar in both species (Fig. S13, S14). This pattern is suggestive of recent selective sweeps in *bicycle* gene regions in both species, which we tested by performing genome-wide scans for positive selection (Pavlidis et al., 2013). Genome-wide signals of selective sweeps were enriched near *bicycle* genes and multiple signals fell within *bicycle* gene clusters in both species (Fig. S14 and S15). Thus, in addition to long-term adaptive protein evolution of bicycle proteins, it appears that strong positive selection has acted recently and presumably frequently near many *bicycle* genes throughout the genome.

In summary, we find evidence for widespread, strong, recent, and frequent positive selection on *bicycle* genes. Since *bicycle* genes are likely secreted from salivary glands specifically in gall-forming aphids, these observations are consistent with the hypothesis that *bicycle* genes encode proteins that are intimately involved in reciprocal molecular evolution between the aphid and their host plant.

## DISCUSSION

We presented multiple lines of evidence that suggest that *bicycle* gene products provide instructive molecules required for aphid gall development. The strongest evidence is that eleven derived regulatory polymorphisms at *dgc* are associated with red galls, with almost complete silencing of *dgc* in aphid salivary glands (Fig. 3A-B, S7B), and with upregulation of seven anthocyanin biosynthetic genes and two red-purple anthocyanins in galls (Fig. 3C-H, S8). Gall color is one small, but convenient, aspect of the panoply of cell biological changes required for gall development. We hypothesize that the product of each *bicycle* gene has its own unique set of targets in the plant and that the combined action of all *bicycle* gene products regulates many aspects of gall development. Testing this hypothesis will require the development of new methods to explore and manipulate this aphid-plant system.

### BICYCLE protein functions

*Dgc* likely encodes a protein that is deposited by aphids into gall tissue, and current evidence suggests that this protein specifically and dramatically results in the downregulation of seven anthocyanin biosynthetic genes. The mechanisms by which this novel aphid protein could alter plant transcription remains to be determined. The primary sequences of DGC and other BICYCLE proteins provide few clues to their molecular mode of action. Outside of the N-terminal secretion signal, BICYCLE proteins possess no similarity with previously reported proteins and display no conserved domains that might guide functional studies. The relatively well-conserved C-Y-C motif appears to define a pair of ∼50-80 aa domains in each protein and the paired cysteines may form disulfide bonds, which is commonly observed for secreted proteins. Secondary structure prediction methods provide little evidence for structural conservation across BICYCLE proteins and the extensive variation in the spacing between conserved cysteines further suggests that BICYCLE proteins may display structural heterogeneity. Identification of their molecular mode of action will require identification of BICYCLE protein binding targets.

### Evolution of *bicycle* genes

We were unable to detect any sequence homology between *bicycle* genes and previously reported genes, which is one reason that it is difficult to infer the molecular function of *bicycle* genes from sequence alone. It is possible that *bicycle* genes evolved *de novo* in an ancestor of gall-forming aphids, perhaps through capture of a 5’ exon encoding an N-terminal signal sequence. However, if *bicycle* genes experienced strong diversifying selection since their origin, perhaps in an ancestor of gall forming aphids about 280 MYA, then the rate of amino-acid substitution that we detected between two closely-related species would likely be sufficient to have eliminated sequence similarity that could be detected by homology-detection algorithms. Identifying the evolutionary antecedents of *bicycle* genes will likely require tracing their evolutionary history across genomes of related species. It may also be more fruitful to use the unusual gene structure of *bicycle* genes (Fig. S10F, G) to search for the antecedents of *bicycle* genes.

## Acknowledgements

We thank Jim Truman for performing the initial aphid salivary gland dissections and for teaching us the tricks, which enabled the entire project, Erika Gajda and Henry Horn for help collecting galls, Patrick Reilly for his genotyping pipeline and helpful conversations, Goran Ceric for getting all software packages to run on the Janelia compute cluster, Tom Dolafi for maintaining the Apollo genome annotation service, Mountain Lakes Biological Station for permission to collect specimens, and Jim Truman, Nicolas Frankel, Richard Mann, Brett Mensh, and Vanessa Ruta for helpful comments on the manuscript.

## STAR Methods

### Lead Contact

Further information and requests for resources and reagents should be directed to and will be fulfilled by the Lead Contact, David L. Stern

### Material Availability

This study did not generate new unique reagents.

### Data and Code Availability

All sequencing data generated during this study is available at the NCBI Short Read Archive and the accession numbers are provided for each sample in the tables below. The genomes are available at Genbank and the genomes and gene annotations (gff files) are available at FigShare.

All of the computer code we produced for this study is freely available at GitHub and links to each piece of code are provided in the detailed descriptions of methods.

---

**Table STAR Methods 1.** Details of biological samples, sequence files, and SRA accession numbers for genome sequencing.

| Sample # | Collection Date | Collection Location | Species | Library Prep Method | Sequencing Method | # Read pairs | Accession # |
| --- | --- | --- | --- | --- | --- | --- | --- |
| MP2 | 15-Jun-2016 | Morven Park, Leesburg, VA | *Hormaphis cornu* | 10X Chromium | Illumina PE151 | 408,093,791 | SRR11397593 |
| MP2 | 15-Jun-2016 | Morven Park, Leesburg, VA | *Hormaphis cornu* | Chicago | Illumina PE151 | 234,423,934 | SRR11397592 |
| 80 | 31-May-2017 | Gap Road, Rt. 651, Leesburg, VA | *Hormaphis cornu* | HiC | Illumina PE151 | 281,999,702 | SRR11397591 |
| ML97Red | 7-Jun-2018 | Mountain Lakes Biological Station, Pembroke, VA | *Hormaphis hamamelidis* | 10X Chromium | Illumina PE151 | 91,611,428 | SRR13258507 |
| Hamamelis virginiana nuclei | 17-Apr-2019 | Janelia Research Campus, Ashburn, VA | *Hamamelis virginiana* | 10X Chromium | Illumina PE150 | 303,926,536 | SRR11397589 |

**Table STAR Methods 2.** Details of biological samples, sequence files, and SRA accession numbers for H. cornu RNA-sequencing. WGB refers to Whole Gene Body library preparation method described in Supplemental Methods. SS refers to the Smart-SCRB library preparation methods described in (Cembrowski et al., 2018). JRC = Janelia Research Campus, Ashburn, VA.

| Sample ID | Collection Location | Collection Date | Stage | Generation | Organ | Library Prep Method | Sequencing Method | # Read pairs | Accession # |
| --- | --- | --- | --- | --- | --- | --- | --- | --- | --- |
| 1st_I_FundSG | JRC | 6-4-2016 | 1 | 1 | SG | WGB | Illumina PE100 | 12643792, 33584393 | SRR11428787 |
| 1st_II_FundSG | JRC | 6-4-2016 | 1 | 1 | SG | WGB | Illumina PE100 | 12412280, 29986671 | SRR11428786 |
| 1st_III_FundSG | JRC | 6-4-2016 | 1 | 1 | SG | WGB | Illumina PE100 | 9566012, 28830029 | SRR11428769 |
| early2nd_I_FundSG | JRC | 6-4-2016 | 2 | 1 | SG | WGB | Illumina PE100 | 11846803, 31258786 | SRR11428758 |
| early2nd_II_FundSG | JRC | 6-4-2016 | 2 | 1 | SG | WGB | Illumina PE100 | 14097211, 33244826 | SRR11428747 |
| early2nd_III_FundSG | JRC | 6-4-2016 | 2 | 1 | SG | WGB | Illumina PE100 | 10610077, 26423617 | SRR11428736 |
| late2nd_I_FundSG_rep1 | JRC | 6-4-2016 | 2 | 1 | SG | WGB | Illumina PE100 | 11783417, 29867606 | SRR11428725 |
| late2nd_II_FundSG_rep1 | JRC | 6-4-2016 | 2 | 1 | SG | WGB | Illumina PE100 | 14189826, 37094462 | SRR11428714 |
| late2nd_III_FundSG_rep1 | JRC | 6-4-2016 | 2 | 1 | SG | WGB | Illumina PE100 | 12137125, 30412534 | SRR11428779 |
| late2nd_IV_FundSG | JRC | 6-4-2016 | 2 | 1 | SG | WGB | Illumina PE100 | 34021474 | SRR11428778 |
| late2nd_V_FundSG | JRC | 6-4-2016 | 2 | 1 | SG | WGB | Illumina PE100 | 33255973 | SRR11428785 |
| late2nd_VI_FundSG | JRC | 6-4-2016 | 2 | 1 | SG | WGB | Illumina PE100 | 29588804 | SRR11428783 |
| 3rd_I_FundSG | JRC | 29-4-2016 | 2 | 1 | SG | WGB | Illumina PE100 | 30833668 | SRR11428777 |
| 3rd_II_FundSG | JRC | 29-4-2016 | 2 | 1 | SG | WGB | Illumina PE100 | 33651454 | SRR11428776 |
| 3rd_III_FundSG | JRC | 29-4-2016 | 2 | 1 | SG | WGB | Illumina PE100 | 40510300 | SRR11428775 |
| 4th_I_FundSG | JRC | 3-5-2016 | 2 | 1 | SG | WGB | Illumina PE100 | 36267453 | SRR11428774 |
| 4th_II_FundSG | JRC | 3-5-2016 | 2 | 1 | SG | WGB | Illumina PE100 | 37712709 | SRR11428773 |
| 4th_III_FundSG | JRC | 3-5-2016 | 2 | 1 | SG | WGB | Illumina PE100 | 37200793 | SRR11428772 |
| adult_I_FundSG | JRC | 3-5-2016 | 2 | 1 | SG | WGB | Illumina PE100 | 37826115 | SRR11428771 |
| adult_II_FundSG | JRC | 3-5-2016 | 2 | 1 | SG | WGB | Illumina PE100 | 31970558 | SRR11428770 |
| adult_III_FundSG | JRC | 4-5-2016 | 2 | 1 | SG | WGB | Illumina PE100 | 29795163 | SRR11428768 |
| whole_1st_I | JRC | 6-4-2016 | 1 | 1 | Whole | WGB | Illumina PE100 | 36313971 | SRR11428767 |
| whole_early 2nd I Fund | JRC | 6-4-2016 | 2 | 1 | Whole | WGB | Illumina PE100 | 44067290 | SRR11428766 |
| whole_late2nd I Fund | JRC | 6-4-2016 | 2 | 1 | Whole | WGB | Illumina PE100 | 32843185 | SRR11428765 |
| whole_v_late2nd I Fund | JRC | 6-4-2016 | 2 | 1 | Whole | WGB | Illumina PE100 | 35509383 | SRR11428764 |
| whole_3rd_I Fund | JRC | 29-4-2016 | 3 | 1 | Whole | WGB | Illumina PE100 | 39418696 | SRR11428763 |
| whole_3rd_I I Fund | JRC | 29-4-2016 | 3 | 1 | Whole | WGB | Illumina PE100 | 40421752 | SRR11428762 |
| whole_4th_I Fund | JRC | 3-5-2016 | 4 | 1 | Whole | WGB | Illumina PE100 | 38585734 | SRR11428761 |
| whole_adult I Fund | JRC | 4-5-2016 | Ad | 1 | Whole | WGB | Illumina PE100 | 42742613 | SRR11428760 |
| 2nd_gen_1st inst SG1 | JRC | 2-6-2018 | 1 | 2 | SG | WGB | Illumina PE150 | 11405653 | SRR11428759 |
| 2nd_gen_1st inst SG2 | JRC | 2-6-2018 | 1 | 2 | SG | WGB | Illumina PE150 | 10794548 | SRR11428757 |
| 2nd_gen_1st inst SG3 | JRC | 2-6-2018 | 1 | 2 | SG | WGB | Illumina PE150 | 12574061 | SRR11428756 |
| 2ndgen_3rdinst sg I 1 | JRC | 19-5-2016 | 3 | 2 | SG | WGB | Illumina PE100 | 20922005 | SRR11428755 |
| 2ndgen_3rdinst sg II 1 | JRC | 19-5-2016 | 3 | 2 | SG | WGB | Illumina PE100 | 19238796 | SRR11428754 |
| 2ndgen_3rdinst sg III 1 | JRC | 19-5-2016 | 3 | 2 | SG | WGB | Illumina PE100 | 19740444 | SRR11428753 |
| 2ndgen_3rdinst_whole_I 1 | JRC | 19-5-2016 | 3 | 2 | Whole | WGB | Illumina PE100 | 18321283 | SRR11428752 |
| 2ndgen_3rdinst_whole_I I 1 | JRC | 19-5-2016 | 3 | 2 | Whole | WGB | Illumina PE100 | 18971169 | SRR11428751 |
| 2ndgen_3rdinst_whole_I II 1 | JRC | 19-5-2016 | 3 | 2 | Whole | WGB | Illumina PE100 | 19463547 | SRR11428750 |
| 2nd_gen_Ad SG4 | JRC | 2-6-2018 | 3 | Ad | SG | WGB | Illumina PE150 | 12714848 | SRR11428749 |
| 2nd_gen_Ad SG5 | JRC | 2-6-2018 | 3 | Ad | SG | WGB | Illumina PE150 | 10858738 | SRR11428748 |
| 2nd_gen_Ad SG6 | JRC | 2-6-2018 | 3 | Ad | SG | WGB | Illumina PE150 | 13120645 | SRR11428746 |
| ex_bet_AdS G8 | JRC | 8-7-2018 | Ad | Ex Betula | SG | WGB | Illumina PE150 | 4914246, 13175533 | SRR11428745 |
| ex_bet_4thS G9 | JRC | 8-7-2018 | 4 | Ex Betula | SG | WGB | Illumina PE150 | 6611299, 18243185 | SRR11428744 |
| ex_bet_4thS G10 | JRC | 8-7-2018 | 4 | Ex Betula | SG | WGB | Illumina PE150 | 6578419, 17834081 | SRR11428743 |
| ex_bet_AdS G11 | JRC | 8-7-2018 | Ad | Ex Betula | SG | WGB | Illumina PE150 | 5627200, 10425494 | SRR11428742 |
| ex_bet_AdS G12 | JRC | 8-7-2018 | Ad | Ex Betula | SG | WGB | Illumina PE150 | 6915842, 13798518 | SRR11428741 |
| ex_bet_AdS<br>G13 | JRC | 8-7-2018 | Ad | Ex<br>Betula | SG | WGB | Illumina<br>PE150 | 7365598,<br>14951729 | SRR1142<br>8740 |
| ex_bet_nym<br>phs_15 | JRC | 8-7-2018 | 4 | Ex<br>Betula | SG | WGB | Illumina<br>PE150 | 11106332 | SRR1142<br>8739 |
| ex_bet_nym<br>phs_16 | JRC | 8-7-2018 | 4 | Ex<br>Betula | SG | WGB | Illumina<br>PE150 | 12481764 | SRR1142<br>8738 |
| sex_SG_19_<br>1 | JRC | 22-10-2019 | Nymp<br>h | Sexual | SG | WGB | Illumina<br>PE150 | 9662029 | SRR1142<br>8737 |
| sex_SG_19_<br>2 | JRC | 22-10-2019 | Nymp<br>h | Sexual | SG | WGB | Illumina<br>PE150 | 12690319 | SRR1142<br>8735 |
| sex_SG_19_<br>3 | JRC | 22-10-2019 | Nymp<br>h | Sexual | SG | WGB | Illumina<br>PE150 | 10875978 | SRR1142<br>8734 |
| sex_SG_19_<br>4 | JRC | 22-10-2019 | Nymp<br>h | Sexual | SG | WGB | Illumina<br>PE150 | 9540662 | SRR1142<br>8733 |
| sex_SG_19_<br>5 | JRC | 22-10-2019 | Nymp<br>h | Sexual | SG | WGB | Illumina<br>PE150 | 9632389 | SRR1142<br>8732 |
| D02 | JRC | 18-11-2018 | Nymp<br>h | Sexual | 3 pooled<br>heads | SS | Illumina<br>SE125 | 23182206 | SRR1142<br>8731 |
| E02 | JRC | 18-11-2018 | Nymp<br>h | Sexual | 3 pooled<br>bodies | SS | Illumina<br>SE125 | 23157666 | SRR1142<br>8730 |
| F02 | JRC | 18-11-2018 | Nymp<br>h | Sexual | 4 pooled<br>salivary<br>glands | SS | Illumina<br>SE125 | 24353653 | SRR1142<br>8729 |
| G02 | JRC | 18-11-2018 | Nymp<br>h | Sexual | 3 pooled<br>salivary<br>glands | SS | Illumina<br>SE125 | 21591985 | SRR1142<br>8728 |
| H02 | JRC | 18-11-2018 | Nymp<br>h | Sexual | 3 pooled<br>salivary<br>glands | SS | Illumina<br>SE125 | 17705013 | SRR1142<br>8727 |

**Table STAR Methods 3.** Details of biological samples, sequence files, and SRA accession numbers for *H. cornu* GWAS whole-genome re-sequencing.

| Sample # | Collection Location | Collection Date | Gall Color | Library Prep Method | Sequencing Method | # Read pairs | Accession # |
| --- | --- | --- | --- | --- | --- | --- | --- |
| B02L02-gr | Moore St & Guyot Ave., Princeton, NJ | 10-May-2017 | Green | Tn5 | Illumina PE150 | 6366491 | SRR11363255 |
| B03L03 | Moore St & Guyot Ave., Princeton, NJ | 10-May-2017 | Green | Tn5 | Illumina PE150 | 4593475 | SRR11363254 |
| GCL74 | Goose Creek Lane, Leesburg, VA | 31-May-2017 | Green | Tn5 | Illumina PE150 | 4809727 | SRR11363163 |
| HW03L02 | Herrontown Woods, Princeton, NJ | 31-May-2017 | Green | Tn5 | Illumina PE150 | 3884184 | SRR11363152 |
| HW03L07 | Herrontown Woods, Princeton, NJ | 31-May-2017 | Green | Tn5 | Illumina PE150 | 8469155 | SRR11363141 |
| HW04L01 | Herrontown Woods, Princeton, NJ | 31-May-2017 | Green | Tn5 | Illumina PE150 | 4589629 | SRR11363130 |
| HW04L07 | Herrontown Woods, Princeton, NJ | 31-May-2017 | Green | Tn5 | Illumina PE150 | 6432189 | SRR11363119 |
| HW05L02 | Herrontown Woods, Princeton, NJ | 31-May-2017 | Green | Tn5 | Illumina PE150 | 5734918 | SRR11363108 |
| HW06L01G01 | Herrontown Woods, Princeton, NJ | 31-May-2017 | Green | Tn5 | Illumina PE150 | 5168885 | SRR11363097 |
| HW07L01 | Herrontown Woods, Princeton, NJ | 31-May-2017 | Green | Tn5 | Illumina PE150 | 5796597 | SRR11363086 |
| HW08L03 | Herrontown Woods, Princeton, NJ | 31-May-2017 | Green | Tn5 | Illumina PE150 | 5326728 | SRR11363253 |
| HW09L05 | Herrontown Woods, Princeton, NJ | 31-May-2017 | Green | Tn5 | Illumina PE150 | 3783083 | SRR11363210 |
| HW10L01 | Herrontown Woods, Princeton, NJ | 31-May-2017 | Green | Tn5 | Illumina PE150 | 3390296 | SRR11363199 |
| HW11L01 | Herrontown Woods, Princeton, NJ | 31-May-2017 | Green | Tn5 | Illumina PE150 | 6969888 | SRR11363188 |
| HW02L05 | Herrontown Woods, Princeton, NJ | 31-May-2017 | Green | Tn5 | Illumina PE150 | 4714807 | SRR11363241 |
| HWL04 | Herrontown Woods, Princeton, NJ | 31-May-2017 | Green | Tn5 | Illumina PE150 | 5643363 | <u>SRR11363230</u> |
| HWL07 | Herrontown Woods, Princeton, NJ | 31-May-2017 | Green | Tn5 | Illumina PE150 | 4917264 | <u>SRR11363187</u> |
| JRC03 | Janelia Research Campus, Ashburn, VA | 3-Apr-2017 | Green | Tn5 | Illumina PE150 | 4863866 | <u>SRR11363176</u> |
| JRC04 | Janelia Research Campus, Ashburn, VA | 3-Apr-2017 | Green | Tn5 | Illumina PE150 | 4523085 | <u>SRR11363165</u> |
| JRC10 | Janelia Research Campus, Ashburn, VA | 3-Apr-2017 | Green | Tn5 | Illumina PE150 | 5182279 | <u>SRR11363164</u> |
| JRC12 | Janelia Research Campus, Ashburn, VA | 3-Apr-2017 | Green | Tn5 | Illumina PE150 | 4178879 | <u>SRR11363162</u> |
| JRC13 | Janelia Research Campus, Ashburn, VA | 3-Apr-2017 | Green | Tn5 | Illumina PE150 | 1985664 | <u>SRR11363161</u> |
| JRC15 | Janelia Research Campus, Ashburn, VA | 3-Apr-2017 | Green | Tn5 | Illumina PE150 | 4191677 | <u>SRR11363160</u> |
| JRC17 | Janelia Research Campus, Ashburn, VA | 3-Apr-2017 | Green | Tn5 | Illumina PE150 | 5230834 | <u>SRR11363159</u> |
| JRC19 | Janelia Research Campus, Ashburn, VA | 3-Apr-2017 | Green | Tn5 | Illumina PE150 | 4844552 | <u>SRR11363158</u> |
| JRC20 | Janelia Research Campus, Ashburn, VA | 3-Apr-2017 | Green | Tn5 | Illumina PE150 | 4750713 | <u>SRR11363157</u> |
| JRC27 | Janelia Research Campus, Ashburn, VA | 3-Apr-2017 | Green | Tn5 | Illumina PE150 | 4868486 | <u>SRR11363156</u> |
| MLR36 | Man Lu Ridge, Maryland | 16-May-2017 | Green | Tn5 | Illumina PE150 | 3547891 | <u>SRR11363155</u> |
| MLR37 | Man Lu Ridge, Maryland | 16-May-2017 | Green | Tn5 | Illumina PE150 | 3401601 | <u>SRR11363154</u> |
| MP55 | Morven Park, Leesburg, VA | 20-May-2017 | Green | Tn5 | Illumina PE150 | 4030936 | <u>SRR11363153</u> |
| MP56 | Morven Park, Leesburg, VA | 20-May-2017 | Green | Tn5 | Illumina PE150 | 5264763 | <u>SRR11363151</u> |
| OWR61 | Old Waterford Road,<br>Leesburg, VA | 20-May-2017 | Green | Tn5 | Illumina PE150 | 4227643 | <u>SRR11363150</u> |
| OWR73 | Old Waterford Road,<br>Leesburg, VA | 30-May-2017 | Green | Tn5 | Illumina PE150 | 4276012 | <u>SRR11363149</u> |
| POR29 | Point of Rocks,<br>Maryland | 16-May-2017 | Green | Tn5 | Illumina PE150 | 3962098 | <u>SRR11363148</u> |
| SM01L01 | St. Michael's Reserve,<br>Princeton, NJ | 1-Jun-2017 | Green | Tn5 | Illumina PE150 | 4883070 | <u>SRR11363147</u> |
| SM02L01 | St. Michael's Reserve,<br>Princeton, NJ | 1-Jun-2017 | Green | Tn5 | Illumina PE150 | 4381921 | <u>SRR11363146</u> |
| SM03L01 | St. Michael's Reserve,<br>Princeton, NJ | 1-Jun-2017 | Green | Tn5 | Illumina PE150 | 4311021 | <u>SRR11363145</u> |
| SM05L01 | St. Michael's Reserve,<br>Princeton, NJ | 1-Jun-2017 | Green | Tn5 | Illumina PE150 | 4506282 | <u>SRR11363144</u> |
| SM05L05 | St. Michael's Reserve,<br>Princeton, NJ | 1-Jun-2017 | Green | Tn5 | Illumina PE150 | 3972013 | <u>SRR11363143</u> |
| W02L01G01 | Woodfield Reservation,<br>Princeton, NJ | 26-May-2017 | Green | Tn5 | Illumina PE150 | 5347819 | <u>SRR11363142</u> |
| W03L01 | Woodfield Reservation,<br>Princeton, NJ | 26-May-2017 | Green | Tn5 | Illumina PE150 | 4536267 | <u>SRR11363140</u> |
| W05L03G02 | Woodfield Reservation,<br>Princeton, NJ | 26-May-2017 | Green | Tn5 | Illumina PE150 | 6397623 | <u>SRR11363139</u> |
| W06L02 | Woodfield Reservation,<br>Princeton, NJ | 26-May-2017 | Green | Tn5 | Illumina PE150 | 6315684 | <u>SRR11363138</u> |
| B01L01 | Moore St & Guyot Ave.,<br>Princeton, NJ | 10-May-2017 | Red | Tn5 | Illumina PE150 | 8135750 | <u>SRR11363137</u> |
| B01L03 | Moore St & Guyot Ave.,<br>Princeton, NJ | 10-May-2017 | Red | Tn5 | Illumina PE150 | 4429167 | <u>SRR11363136</u> |
| B02L01 | Moore St & Guyot Ave.,<br>Princeton, NJ | 10-May-2017 | Red | Tn5 | Illumina PE150 | 4620277 | <u>SRR11363135</u> |
| B02L02-red | Moore St & Guyot Ave.,<br>Princeton, NJ | 10-May-2017 | Red | Tn5 | Illumina PE150 | 5625715 | <u>SRR11363134</u> |
| B03L01 | Moore St & Guyot Ave.,<br>Princeton, NJ | 10-May-2017 | Red | Tn5 | Illumina PE150 | 10312327 | <u>SRR11363133</u> |
| HW02L02 | Herrontown Woods,<br>Princeton, NJ | 31-May-2017 | Red | Tn5 | Illumina PE150 | 4039449 | <u>SRR11363132</u> |
| HW03L01 | Herrontown Woods, Princeton, NJ | 31-May-2017 | Red | Tn5 | Illumina PE150 | 4130302 | <u>SRR11363131</u> |
| HW03L03 | Herrontown Woods, Princeton, NJ | 31-May-2017 | Red | Tn5 | Illumina PE150 | 3384712 | <u>SRR11363129</u> |
| HW03L04 | Herrontown Woods, Princeton, NJ | 31-May-2017 | Red | Tn5 | Illumina PE150 | 3904224 | <u>SRR11363128</u> |
| HW06L01G02 | Herrontown Woods, Princeton, NJ | 31-May-2017 | Red | Tn5 | Illumina PE150 | 5403059 | <u>SRR11363127</u> |
| HW07L02 | Herrontown Woods, Princeton, NJ | 31-May-2017 | Red | Tn5 | Illumina PE150 | 8799194 | <u>SRR11363126</u> |
| HW09L02 | Herrontown Woods, Princeton, NJ | 31-May-2017 | Red | Tn5 | Illumina PE150 | 3673337 | <u>SRR11363125</u> |
| HW09L03 | Herrontown Woods, Princeton, NJ | 31-May-2017 | Red | Tn5 | Illumina PE150 | 5235539 | <u>SRR11363124</u> |
| HW09L07 | Herrontown Woods, Princeton, NJ | 31-May-2017 | Red | Tn5 | Illumina PE150 | 8256988 | <u>SRR11363123</u> |
| HW10L02 | Herrontown Woods, Princeton, NJ | 31-May-2017 | Red | Tn5 | Illumina PE150 | 5492996 | <u>SRR11363122</u> |
| HWL03 | Herrontown Woods, Princeton, NJ | 31-May-2017 | Red | Tn5 | Illumina PE150 | 7814556 | <u>SRR11363121</u> |
| JRC01 | Janelia Research Campus, Ashburn, VA | 3-Apr-2017 | Red | Tn5 | Illumina PE150 | 4617289 | <u>SRR11363120</u> |
| JRC06 | Janelia Research Campus, Ashburn, VA | 3-Apr-2017 | Red | Tn5 | Illumina PE150 | 5709657 | <u>SRR11363118</u> |
| JRC11 | Janelia Research Campus, Ashburn, VA | 3-Apr-2017 | Red | Tn5 | Illumina PE150 | 4441152 | <u>SRR11363117</u> |
| JRC14 | Janelia Research Campus, Ashburn, VA | 3-Apr-2017 | Red | Tn5 | Illumina PE150 | 4255537 | <u>SRR11363116</u> |
| JRC16 | Janelia Research Campus, Ashburn, VA | 3-Apr-2017 | Red | Tn5 | Illumina PE150 | 7587374 | <u>SRR11363115</u> |
| JRC18 | Janelia Research Campus, Ashburn, VA | 3-Apr-2017 | Red | Tn5 | Illumina PE150 | 4944638 | <u>SRR11363114</u> |
| JRC22 | Janelia Research Campus, Ashburn, VA | 3-Apr-2017 | Red | Tn5 | Illumina PE150 | 6692470 | <u>SRR11363113</u> |
| JRC23 | Janelia Research Campus, Ashburn, VA | 3-Apr-2017 | Red | Tn5 | Illumina PE150 | 4475288 | <u>SRR11363112</u> |
| JRC25 | Janelia Research Campus, Ashburn, VA | 3-Apr-2017 | Red | Tn5 | Illumina PE150 | 5506970 | <u>SRR11363111</u> |
| JRC84 | Janelia Research Campus, Ashburn, VA | 21-May-2017 | Red | Tn5 | Illumina PE150 | 7362956 | <u>SRR11363110</u> |
| JRC85 | Janelia Research Campus, Ashburn, VA | 21-May-2017 | Red | Tn5 | Illumina PE150 | 7150172 | <u>SRR11363109</u> |
| JRC86 | Janelia Research Campus, Ashburn, VA | 21-May-2017 | Red | Tn5 | Illumina PE150 | 6592107 | <u>SRR11363107</u> |
| JRC87 | Janelia Research Campus, Ashburn, VA | 21-May-2017 | Red | Tn5 | Illumina PE150 | 6071483 | <u>SRR11363106</u> |
| JRC88 | Janelia Research Campus, Ashburn, VA | 21-May-2017 | Red | Tn5 | Illumina PE150 | 3476862 | <u>SRR11363105</u> |
| JRC89 | Janelia Research Campus, Ashburn, VA | 21-May-2017 | Red | Tn5 | Illumina PE150 | 6474867 | <u>SRR11363104</u> |
| JRC90 | Janelia Research Campus, Ashburn, VA | 21-May-2017 | Red | Tn5 | Illumina PE150 | 7186265 | <u>SRR11363103</u> |
| JRC91 | Janelia Research Campus, Ashburn, VA | 21-May-2017 | Red | Tn5 | Illumina PE150 | 6643361 | <u>SRR11363102</u> |
| JRC92 | Janelia Research Campus, Ashburn, VA | 21-May-2017 | Red | Tn5 | Illumina PE150 | 4674510 | <u>SRR11363101</u> |
| JRC93 | Janelia Research Campus, Ashburn, VA | 21-May-2017 | Red | Tn5 | Illumina PE150 | 5182706 | <u>SRR11363100</u> |
| JRC94 | Janelia Research | 21-May-2017 | Red | Tn5 | Illumina PE150 | 4946068 | <u>SRR11363099</u> |
|  | Campus,<br>Ashburn, VA |  |  |  |  |  |  |
| JRC95 | Janelia<br>Research<br>Campus,<br>Ashburn, VA | 21-May-2017 | Red | Tn5 | Illumina<br>PE150 | 7026186 | <u>SRR11363098</u> |
| JRC96 | Janelia<br>Research<br>Campus,<br>Ashburn, VA | 21-May-2017 | Red | Tn5 | Illumina<br>PE150 | 4946848 | <u>SRR11363096</u> |
| JRC97 | Janelia<br>Research<br>Campus,<br>Ashburn, VA | 21-May-2017 | Red | Tn5 | Illumina<br>PE150 | 6115771 | <u>SRR11363095</u> |
| OWR71 | Old Waterford<br>Road,<br>Leesburg, VA | 30-May-2017 | Red | Tn5 | Illumina<br>PE150 | 5324101 | <u>SRR11363094</u> |
| POR30 | Point of Rocks,<br>Maryland | 16-May-2017 | Red | Tn5 | Illumina<br>PE150 | 3709000 | <u>SRR11363093</u> |
| POR31 | Point of Rocks,<br>Maryland | 16-May-2017 | Red | Tn5 | Illumina<br>PE150 | 6515377 | <u>SRR11363092</u> |
| SM05L02 | St. Michael's<br>Reserve,<br>Princeton, NJ | 1-Jun-2017 | Red | Tn5 | Illumina<br>PE150 | 5328410 | <u>SRR11363091</u> |
| SM05L04 | St. Michael's<br>Reserve,<br>Princeton, NJ | 1-Jun-2017 | Red | Tn5 | Illumina<br>PE150 | 4428657 | <u>SRR11363090</u> |
| W02L02 | Woodfield<br>Reservation,<br>Princeton, NJ | 31-May-2017 | Red | Tn5 | Illumina<br>PE150 | 4068904 | <u>SRR11363089</u> |
| W02L06 | Woodfield<br>Reservation,<br>Princeton, NJ | 31-May-2017 | Red | Tn5 | Illumina<br>PE150 | 3336967 | <u>SRR11363088</u> |
| W05L05 | Woodfield<br>Reservation,<br>Princeton, NJ | 31-May-2017 | Red | Tn5 | Illumina<br>PE150 | 4073188 | <u>SRR11363087</u> |

**Table STAR Methods 4.** Details of biological samples, sequence files, and SRA accession numbers for H. cornu GWAS targeted re-sequencing of approximately 800 kbp of Chromosome 1.

| Sample # | Collection Location | Collection Date | Gall Color | Library Prep Method | Sequencing Method | # Read pairs | Accession # |
| --- | --- | --- | --- | --- | --- | --- | --- |
| B01L02 | Moore St & Guyot Ave., Princeton, NJ | 10-May-2017 | Green | Tn5 | Illumina PE150 | 214163 | SRR11363085 |
| B02L02-gr | Moore St & Guyot Ave., Princeton, NJ | 10-May-2017 | Green | Tn5 | Illumina PE150 | 190639 | SRR11363084 |
| B03L03 | Moore St & Guyot Ave., Princeton, NJ | 10-May-2017 | Green | Tn5 | Illumina PE150 | 212907 | SRR11363083 |
| GCL74 | Goose Creek Lane, Leesburg, VA | 31-May-2017 | Green | Tn5 | Illumina PE150 | 306230 | SRR11363082 |
| HW03L02 | Herrontown Woods, Princeton, NJ | 31-May-2017 | Green | Tn5 | Illumina PE150 | 291669 | SRR11363081 |
| HW03L07 | Herrontown Woods, Princeton, NJ | 31-May-2017 | Green | Tn5 | Illumina PE150 | 306531 | SRR11363080 |
| HW04L07 | Herrontown Woods, Princeton, NJ | 31-May-2017 | Green | Tn5 | Illumina PE150 | 270871 | SRR11363079 |
| HW05L02 | Herrontown Woods, Princeton, NJ | 31-May-2017 | Green | Tn5 | Illumina PE150 | 262310 | SRR11363078 |
| HW06L01G01 | Herrontown Woods, Princeton, NJ | 31-May-2017 | Green | Tn5 | Illumina PE150 | 269379 | SRR11363077 |
| HW07L01 | Herrontown Woods, Princeton, NJ | 31-May-2017 | Green | Tn5 | Illumina PE150 | 237951 | SRR11363076 |
| HW08L03 | Herrontown Woods, Princeton, NJ | 31-May-2017 | Green | Tn5 | Illumina PE150 | 257010 | SRR11363252 |
| HW09L05 | Herrontown Woods, Princeton, NJ | 31-May-2017 | Green | Tn5 | Illumina PE150 | 211921 | SRR11363219 |
| HW10L01 | Herrontown Woods, Princeton, NJ | 31-May-2017 | Green | Tn5 | Illumina PE150 | 241629 | SRR11363218 |
| HW11L01 | Herrontown Woods, Princeton, NJ | 31-May-2017 | Green | Tn5 | Illumina PE150 | 237027 | SRR11363217 |
| HWL02L05 | Herrontown Woods, Princeton, NJ | 31-May-2017 | Green | Tn5 | Illumina PE150 | 272737 | SRR11363216 |
| HWL04 | Herrontown Woods, Princeton, NJ | 31-May-2017 | Green | Tn5 | Illumina PE150 | 249525 | SRR11363215 |
| HWL07 | Herrontown Woods, Princeton, NJ | 31-May-2017 | Green | Tn5 | Illumina PE150 | 304184 | SRR11363214 |
| JH01L04G01 | Jockey Hollow, Morristown, NJ | 12-Jun-2017 | Green | Tn5 | Illumina PE150 | 225593 | SRR11363213 |
| JH02L04G02 | Jockey Hollow, Morristown, NJ | 3-Apr-2017 | Green | Tn5 | Illumina PE150 | 241035 | SRR11363212 |
| JRC03 | Janelia Research Campus, Ashburn, VA | 3-Apr-2017 | Green | Tn5 | Illumina PE150 | 270579 | SRR11363211 |
| JRC04 | Janelia Research Campus, Ashburn, VA | 3-Apr-2017 | Green | Tn5 | Illumina PE150 | 291029 | SRR11363209 |
| JRC12 | Janelia Research Campus, Ashburn, VA | 3-Apr-2017 | Green | Tn5 | Illumina PE150 | 291716 | SRR11363208 |
| JRC15 | Janelia Research Campus, Ashburn, VA | 3-Apr-2017 | Green | Tn5 | Illumina PE150 | 278157 | SRR11363207 |
| JRC19 | Janelia Research Campus, Ashburn, VA | 3-Apr-2017 | Green | Tn5 | Illumina PE150 | 296693 | SRR11363206 |
| JRC27 | Janelia Research Campus, Ashburn, VA | 3-Apr-2017 | Green | Tn5 | Illumina PE150 | 289882 | SRR11363205 |
| MLR36 | Man Lu Ridge Rd., Jefferson, MD | 16-May-2017 | Green | Tn5 | Illumina PE150 | 237264 | SRR11363204 |
| MLR37 | Man Lu Ridge Rd., Jefferson, MD | 16-May-2017 | Green | Tn5 | Illumina PE150 | 215201 | SRR11363203 |
| MP55 | Morven Park, Leesburg, VA | 20-May-2017 | Green | Tn5 | Illumina PE150 | 233325 | SRR11363202 |
| MP56 | Morven Park, Leesburg, VA | 20-May-2017 | Green | Tn5 | Illumina PE150 | 201670 | SRR11363201 |
| OWR61 | Old Waterford Road, Leesburg, VA | 20-May-2017 | Green | Tn5 | Illumina PE150 | 175533 | SRR11363200 |
| OWR73 | Old Waterford Road, Leesburg, VA | 20-May-2017 | Green | Tn5 | Illumina PE150 | 132423 | SRR11363198 |
| POR29 | Point of Rocks, MD | 16-May-2017 | Green | Tn5 | Illumina PE150 | 236262 | SRR11363197 |
| SM01L01 | St. Michael's Reserve, Princeton, NJ | 1-Jun-2017 | Green | Tn5 | Illumina PE150 | 325832 | SRR11363196 |
| SM02L01 | St. Michael's Reserve, Princeton, NJ | 1-Jun-2017 | Green | Tn5 | Illumina PE150 | 313663 | SRR11363195 |
| SM03L01 | St. Michael's Reserve, Princeton, NJ | 1-Jun-2017 | Green | Tn5 | Illumina PE150 | 257124 | SRR11363194 |
| SM05L01 | St. Michael's Reserve, Princeton, NJ | 1-Jun-2017 | Green | Tn5 | Illumina PE150 | 292603 | SRR11363193 |
| SM05L05 | St. Michael's Reserve, Princeton, NJ | 1-Jun-2017 | Green | Tn5 | Illumina PE150 | 231169 | SRR11363192 |
| W01L01 | Woodfield Reservation, Princeton, NJ | 26-May-2017 | Green | Tn5 | Illumina PE150 | 300257 | SRR11363191 |
| W02L01G01 | Woodfield Reservation, Princeton, NJ | 26-May-2017 | Green | Tn5 | Illumina PE150 | 315314 | SRR11363190 |
| W03L01 | Woodfield Reservation, Princeton, NJ | 26-May-2017 | Green | Tn5 | Illumina PE150 | 313024 | SRR11363189 |
| W05L03G02 | Woodfield Reservation, Princeton, NJ | 26-May-2017 | Green | Tn5 | Illumina PE150 | 223027 | SRR11363251 |
| W06L02 | Woodfield Reservation, Princeton, NJ | 26-May-2017 | Green | Tn5 | Illumina PE150 | 303327 | SRR11363250 |
| B01L01 | Moore St. & Guyot Ave., Princeton, NJ | 10-May-2017 | Red | Tn5 | Illumina PE150 | 303803 | SRR11363249 |
| B01L03 | Moore St & Guyot Ave., Princeton, NJ | 10-May-2017 | Red | Tn5 | Illumina PE150 | 323593 | SRR11363248 |
| B02L01 | Moore St & Guyot Ave., Princeton, NJ | 10-May-2017 | Red | Tn5 | Illumina PE150 | 256759 | SRR11363247 |
| B02L02 | Moore St & Guyot Ave., Princeton, NJ | 10-May-2017 | Red | Tn5 | Illumina PE150 | 305206 | SRR11363246 |
| B03L01 | Moore St & Guyot Ave., Princeton, NJ | 10-May-2017 | Red | Tn5 | Illumina PE150 | 319941 | SRR11363245 |
| HW02L02 | Herrontown woods, NJ | 31-May-2017 | Red | Tn5 | Illumina PE150 | 333873 | SRR11363244 |
| HW03L01 | Herrontown woods, NJ | 31-May-2017 | Red | Tn5 | Illumina PE150 | 258120 | SRR11363243 |
| HW03L03 | Herrontown woods, NJ | 31-May-2017 | Red | Tn5 | Illumina PE150 | 275293 | SRR11363242 |
| HW03L04 | Herrontown woods, NJ | 31-May-2017 | Red | Tn5 | Illumina PE150 | 299356 | SRR11363240 |
| HW06L01G02 | Herrontown woods, NJ | 31-May-2017 | Red | Tn5 | Illumina PE150 | 260553 | SRR11363239 |
| HW06L02G03 | Herrontown woods, NJ | 31-May-2017 | Red | Tn5 | Illumina PE150 | 300112 | SRR11363238 |
| HW07L02 | Herrontown woods, NJ | 31-May-2017 | Red | Tn5 | Illumina PE150 | 259311 | SRR11363237 |
| HW09L02 | Herrontown woods, NJ | 31-May-2017 | Red | Tn5 | Illumina PE150 | 260464 | SRR11363236 |
| HW09L03 | Herrontown woods, NJ | 31-May-2017 | Red | Tn5 | Illumina PE150 | 247150 | SRR11363235 |
| HW09L07 | Herrontown woods, NJ | 31-May-2017 | Red | Tn5 | Illumina PE150 | 252787 | SRR11363234 |
| HW10L02 | Herrontown woods, NJ | 31-May-2017 | Red | Tn5 | Illumina PE150 | 254752 | SRR11363233 |
| HWL03 | Herrontown woods, NJ | 31-May-2017 | Red | Tn5 | Illumina PE150 | 254968 | SRR11363232 |
| JRC01 | Janelia Research Campus, Ashburn, VA | 3-Apr-2017 | Red | Tn5 | Illumina PE150 | 252369 | SRR11363231 |
| JRC06 | Janelia Research Campus, Ashburn, VA | 3-Apr-2017 | Red | Tn5 | Illumina PE150 | 317542 | SRR11363229 |
| JRC11 | Janelia Research Campus, Ashburn, VA | 3-Apr-2017 | Red | Tn5 | Illumina PE150 | 304971 | SRR11363228 |
| JRC14 | Janelia Research Campus, Ashburn, VA | 3-Apr-2017 | Red | Tn5 | Illumina PE150 | 327441 | SRR11363227 |
| JRC16 | Janelia Research Campus, Ashburn, VA | 3-Apr-2017 | Red | Tn5 | Illumina PE150 | 253464 | SRR11363226 |
| JRC18 | Janelia Research Campus, Ashburn, VA | 3-Apr-2017 | Red | Tn5 | Illumina PE150 | 362357 | SRR11363225 |
| JRC22 | Janelia Research Campus, Ashburn, VA | 3-Apr-2017 | Red | Tn5 | Illumina PE150 | 226260 | SRR11363224 |
| JRC23 | Janelia Research Campus, Ashburn, VA | 3-Apr-2017 | Red | Tn5 | Illumina PE150 | 338219 | SRR11363223 |
| JRC25 | Janelia Research Campus, Ashburn, VA | 3-Apr-2017 | Red | Tn5 | Illumina PE150 | 235476 | SRR11363222 |
| JRC84 | Janelia Research Campus, Ashburn, VA | 21-May-2017 | Red | Tn5 | Illumina PE150 | 244087 | SRR11363110 |
| JRC85 | Janelia Research Campus, Ashburn, VA | 21-May-2017 | Red | Tn5 | Illumina PE150 | 242809 | SRR11363220 |
| JRC86 | Janelia Research Campus, Ashburn, VA | 21-May-2017 | Red | Tn5 | Illumina PE150 | 277938 | SRR11363186 |
| JRC87 | Janelia Research Campus, Ashburn, VA | 21-May-2017 | Red | Tn5 | Illumina PE150 | 294687 | SRR11363185 |
| JRC88 | Janelia Research Campus, Ashburn, VA | 21-May-2017 | Red | Tn5 | Illumina PE150 | 283813 | SRR11363184 |
| JRC89 | Janelia Research Campus, Ashburn, VA | 21-May-2017 | Red | Tn5 | Illumina PE150 | 293816 | SRR11363183 |
| JRC90 | Janelia Research Campus, Ashburn, VA | 21-May-2017 | Red | Tn5 | Illumina PE150 | 307410 | SRR11363182 |
| JRC91 | Janelia Research Campus, Ashburn, VA | 21-May-2017 | Red | Tn5 | Illumina PE150 | 301578 | SRR11363181 |
| JRC92 | Janelia Research Campus, Ashburn, VA | 21-May-2017 | Red | Tn5 | Illumina PE150 | 274484 | SRR11363180 |
| JRC93 | Janelia Research Campus, Ashburn, VA | 21-May-2017 | Red | Tn5 | Illumina PE150 | 251875 | SRR11363179 |
| JRC94 | Janelia Research Campus, Ashburn, VA | 21-May-2017 | Red | Tn5 | Illumina PE150 | 246812 | SRR11363178 |
| JRC95 | Janelia Research Campus, Ashburn, VA | 21-May-2017 | Red | Tn5 | Illumina PE150 | 255648 | SRR11363177 |
| JRC96 | Janelia Research Campus, Ashburn, VA | 21-May-2017 | Red | Tn5 | Illumina PE150 | 228158 | SRR11363175 |
| JRC97 | Janelia Research Campus, Ashburn, VA | 21-May-2017 | Red | Tn5 | Illumina PE150 | 242492 | SRR11363174 |
| OWR71 | Old Waterford Road, Leesburg, VA | 30-May-2017 | Red | Tn5 | Illumina PE150 | 271725 | SRR11363173 |
| POR30 | Point of Rocks, MD | 16-May-2017 | Red | Tn5 | Illumina PE150 | 264484 | SRR11363172 |
| POR31 | Point of Rocks, MD | 16-May-2017 | Red | Tn5 | Illumina PE150 | 132066 | SRR11363171 |
| SM05L02 | St. Michael's preserve, NJ | 1-Jun-2017 | Red | Tn5 | Illumina PE150 | 317520 | SRR11363170 |
| SM05L04 | St. Michael's preserve, NJ | 1-Jun-2017 | Red | Tn5 | Illumina PE150 | 327054 | SRR11363169 |
| W02L02 | Woodfield reservation, NJ | 31-May-2017 | Red | Tn5 | Illumina PE150 | 272076 | SRR11363168 |
| W02L06 | Woodfield reservation, NJ | 31-May-2017 | Red | Tn5 | Illumina PE150 | 266667 | SRR11363167 |
| W05L05 | Woodfield reservation, NJ | 31-May-2017 | Red | Tn5 | Illumina PE150 | 233816 | SRR11363166 |

**Table STAR Methods 5.** Details of biological samples, sequence files, and SRA accession numbers for *H. hamamelidis* whole-genome re-sequencing. MLBS = Mountain Lakes Biological Station, Pembroke, Virginia.

| Sample # | Collection Location | Collection Date | Library Prep Method | Sequencing Method | # Read pairs | Accession # |
| --- | --- | --- | --- | --- | --- | --- |
| GCC77 | Goose Creek Lane, Leesburg, VA | 31-May-2017 | Tn5 | Illumina PE150 | 7327275 | SRR13258155 |
| ML10 Gr | MLBS | 7-Jul-2018 | Tn5 | Illumina PE150 | 7333060 | SRR13258154 |
| ML14 Gr | MLBS | 7-Jul-2018 | Tn5 | Illumina PE150 | 7868181 | SRR13258160 |
| ML15 Gr | MLBS | 7-Jul-2018 | Tn5 | Illumina PE150 | 6656988 | SRR13258022 |
| ML17 Gr | MLBS | 7-Jul-2018 | Tn5 | Illumina PE150 | 7010451 | SRR13257979 |
| ML18 Gr | MLBS | 7-Jul-2018 | Tn5 | Illumina PE150 | 6481897 | SRR13258033 |
| ML2 Gr | MLBS | 7-Jul-2018 | Tn5 | Illumina PE150 | 7344032 | SRR13258044 |
| ML21 Gr | MLBS | 7-Jul-2018 | Tn5 | Illumina PE150 | 7166893 | SRR13257983 |
| ML23 Gr | MLBS | 7-Jul-2018 | Tn5 | Illumina PE150 | 7503826 | SRR13258124 |
| ML24 Gr | MLBS | 7-Jul-2018 | Tn5 | Illumina PE150 | 4641363 | SRR13257994 |
| ML27 Gr | MLBS | 7-Jul-2018 | Tn5 | Illumina PE150 | 6152152 | SRR13258153 |
| ML28 Gr | MLBS | 7-Jul-2018 | Tn5 | Illumina PE150 | 7763943 | SRR13258063 |
| ML30 Gr | MLBS | 7-Jul-2018 | Tn5 | Illumina PE150 | 6873459 | SRR13258068 |
| ML33 Gr | MLBS | 7-Jul-2018 | Tn5 | Illumina PE150 | 5378458 | SRR13258073 |
| ML34 Gr | MLBS | 7-Jul-2018 | Tn5 | Illumina PE150 | 6891338 | SRR13258084 |
| ML36 Gr | MLBS | 7-Jul-2018 | Tn5 | Illumina PE150 | 228271 | SRR13258094 |
| ML38 Gr | MLBS | 7-Jul-2018 | Tn5 | Illumina PE150 | 7212647 | SRR13258102 |
| ML40 Gr | MLBS | 7-Jul-2018 | Tn5 | Illumina PE150 | 6162598 | SRR13258113 |
| ML41 Gr | MLBS | 7-Jul-2018 | Tn5 | Illumina PE150 | 6170559 | SRR13258002 |
| ML43 Gr | MLBS | 7-Jul-2018 | Tn5 | Illumina PE150 | 7580980 | SRR13258012 |
| ML44 Gr | MLBS | 7-Jul-2018 | Tn5 | Illumina PE150 | 7912117 | SRR13258161 |
| ML48 Gr | MLBS | 7-Jul-2018 | Tn5 | Illumina PE150 | 5620965 | SRR13258162 |
| ML49 Gr | MLBS | 7-Jul-2018 | Tn5 | Illumina PE150 | 4744505 | SRR13257971 |
| ML50_Gr | MLBS | 7-Jul-2018 | Tn5 | Illumina<br>PE150 | 7383271 | SRR13258015 |
| ML51_Gr | MLBS | 7-Jul-2018 | Tn5 | Illumina<br>PE150 | 7607244 | SRR13258016 |
| ML53_Gr | MLBS | 7-Jul-2018 | Tn5 | Illumina<br>PE150 | 5317613 | SRR13258017 |
| ML55_Gr | MLBS | 7-Jul-2018 | Tn5 | Illumina<br>PE150 | 7580879 | SRR13258018 |
| ML57_Gr | MLBS | 7-Jul-2018 | Tn5 | Illumina<br>PE150 | 1605322 | SRR13258019 |
| ML6_Gr | MLBS | 7-Jul-2018 | Tn5 | Illumina<br>PE150 | 7702032 | SRR13258020 |
| ML60_Gr | MLBS | 7-Jul-2018 | Tn5 | Illumina<br>PE150 | 7457247 | SRR13258021 |
| ML61_Gr | MLBS | 7-Jul-2018 | Tn5 | Illumina<br>PE150 | 7359768 | SRR13258023 |
| ML62_Gr | MLBS | 7-Jul-2018 | Tn5 | Illumina<br>PE150 | 7278081 | SRR13258024 |
| ML65_Gr | MLBS | 7-Jul-2018 | Tn5 | Illumina<br>PE150 | 7448093 | SRR13258025 |
| ML70_Gr | MLBS | 7-Jul-2018 | Tn5 | Illumina<br>PE150 | 6912997 | SRR13257972 |
| ML71_Gr | MLBS | 7-Jul-2018 | Tn5 | Illumina<br>PE150 | 8049317 | SRR13257973 |
| ML73_Gr | MLBS | 7-Jul-2018 | Tn5 | Illumina<br>PE150 | 7124219 | SRR13257974 |
| ML77_Gr | MLBS | 7-Jul-2018 | Tn5 | Illumina<br>PE150 | 8407556 | SRR13257975 |
| ML79_Gr | MLBS | 7-Jul-2018 | Tn5 | Illumina<br>PE150 | 8352011 | SRR13257976 |
| ML82_Gr | MLBS | 7-Jul-2018 | Tn5 | Illumina<br>PE150 | 6974944 | SRR13257977 |
| MLR38 | Man Lu Ridge Rd.,<br>Maryland | 16-May-2017 | Tn5 | Illumina<br>PE150 | 7732998 | SRR13257978 |
| MLR40 | Man Lu Ridge Rd.,<br>Maryland | 16-May-2017 | Tn5 | Illumina<br>PE150 | 8121428 | SRR13257980 |
| MLR50 | Man Lu Ridge Rd.,<br>Maryland | 16-May-2017 | Tn5 | Illumina<br>PE150 | 6605756 | SRR13257981 |
| MLR51 | Man Lu Ridge Rd.,<br>Maryland | 16-May-2017 | Tn5 | Illumina<br>PE150 | 7442782 | SRR13257982 |
| MLR52 | Man Lu Ridge Rd.,<br>Maryland | 16-May-2017 | Tn5 | Illumina<br>PE150 | 7346231 | SRR13258026 |
| MLR53 | Man Lu Ridge Rd.,<br>Maryland | 16-May-2017 | Tn5 | Illumina<br>PE150 | 7679601 | SRR13258027 |
| MLR54 | Man Lu Ridge Rd.,<br>Maryland | 16-May-2017 | Tn5 | Illumina<br>PE150 | 6710199 | SRR13258028 |
| OWR57 | Old Waterford<br>Road, Leesburg,<br>VA | 22-May-2017 | Tn5 | Illumina<br>PE150 | 7307248 | SRR13258029 |
| OWR70 | Old Waterford<br>Road, Leesburg,<br>VA | 30-May-2017 | Tn5 | Illumina<br>PE150 | 6821266 | SRR13258030 |
| ML10_Red | MLBS | 7-Jul-2018 | Tn5 | Illumina<br>PE150 | 7157138 | SRR13258031 |
| ML12_Red | MLBS | 7-Jul-2018 | Tn5 | Illumina<br>PE150 | 135505 | SRR13258032 |
| ML13_Red | MLBS | 7-Jul-2018 | Tn5 | Illumina<br>PE150 | 8118063 | SRR13258034 |
| ML16_Red | MLBS | 7-Jul-2018 | Tn5 | Illumina<br>PE150 | 9041960 | SRR13258035 |
| ML18_Red | MLBS | 7-Jul-2018 | Tn5 | Illumina<br>PE150 | 8826653 | SRR13258036 |
| ML19_Red | MLBS | 7-Jul-2018 | Tn5 | Illumina<br>PE150 | 8143654 | SRR13258037 |
| ML2_Red | MLBS | 7-Jul-2018 | Tn5 | Illumina<br>PE150 | 7175142 | SRR13258038 |
| ML20_Red | MLBS | 7-Jul-2018 | Tn5 | Illumina<br>PE150 | 8225262 | SRR13258039 |
| ML22_Red | MLBS | 7-Jul-2018 | Tn5 | Illumina<br>PE150 | 8664565 | SRR13258040 |
| ML23_Red | MLBS | 7-Jul-2018 | Tn5 | Illumina<br>PE150 | 8102497 | SRR13258041 |
| ML24_Red | MLBS | 7-Jul-2018 | Tn5 | Illumina<br>PE150 | 7884498 | SRR13258042 |
| ML25_Red | MLBS | 7-Jul-2018 | Tn5 | Illumina<br>PE150 | 8547869 | SRR13258043 |
| ML27_Red | MLBS | 7-Jul-2018 | Tn5 | Illumina<br>PE150 | 7831540 | SRR13258045 |
| ML29_Red | MLBS | 7-Jul-2018 | Tn5 | Illumina<br>PE150 | 7930016 | SRR13258129 |
| ML3_Red | MLBS | 7-Jul-2018 | Tn5 | Illumina<br>PE150 | 7532536 | SRR13258128 |
| ML32_Red | MLBS | 7-Jul-2018 | Tn5 | Illumina<br>PE150 | 7411318 | SRR13258046 |
| ML35_Red | MLBS | 7-Jul-2018 | Tn5 | Illumina<br>PE150 | 7957062 | SRR13258047 |
| ML36_Red | MLBS | 7-Jul-2018 | Tn5 | Illumina<br>PE150 | 8051736 | SRR13258048 |
| ML37_Red | MLBS | 7-Jul-2018 | Tn5 | Illumina<br>PE150 | 8015006 | SRR13258049 |
| ML38_Red | MLBS | 7-Jul-2018 | Tn5 | Illumina<br>PE150 | 8549973 | SRR13258050 |
| ML39_Red | MLBS | 7-Jul-2018 | Tn5 | Illumina<br>PE150 | 7014367 | SRR13258051 |
| ML41_Red | MLBS | 7-Jul-2018 | Tn5 | Illumina<br>PE150 | 7008886 | SRR13258052 |
| ML42_Red | MLBS | 7-Jul-2018 | Tn5 | Illumina<br>PE150 | 6255046 | SRR13257984 |
| ML44_Red | MLBS | 7-Jul-2018 | Tn5 | Illumina<br>PE150 | 8196265 | SRR13257985 |
| ML45_Red | MLBS | 7-Jul-2018 | Tn5 | Illumina<br>PE150 | 7370732 | SRR13257986 |
| ML46_Red | MLBS | 7-Jul-2018 | Tn5 | Illumina<br>PE150 | 7101764 | SRR13258053 |
| ML47_Red | MLBS | 7-Jul-2018 | Tn5 | Illumina<br>PE150 | 7935806 | SRR13258054 |
| ML52_Red | MLBS | 7-Jul-2018 | Tn5 | Illumina<br>PE150 | 8092411 | SRR13258055 |
| ML54_Red | MLBS | 7-Jul-2018 | Tn5 | Illumina PE150 | 5607371 | SRR13258056 |
| ML56_Red | MLBS | 7-Jul-2018 | Tn5 | Illumina PE150 | 7892227 | SRR13258121 |
| ML57_Red | MLBS | 7-Jul-2018 | Tn5 | Illumina PE150 | 6630514 | SRR13258122 |
| ML58_Red | MLBS | 7-Jul-2018 | Tn5 | Illumina PE150 | 7561525 | SRR13258123 |
| ML59_Red | MLBS | 7-Jul-2018 | Tn5 | Illumina PE150 | 7637387 | SRR13258125 |
| ML62_Red | MLBS | 7-Jul-2018 | Tn5 | Illumina PE150 | 7912262 | SRR13258126 |
| ML63_Red | MLBS | 7-Jul-2018 | Tn5 | Illumina PE150 | 6922716 | SRR13258127 |
| ML64_Red | MLBS | 7-Jul-2018 | Tn5 | Illumina PE150 | 7470732 | SRR13257987 |
| ML65_Red | MLBS | 7-Jul-2018 | Tn5 | Illumina PE150 | 7195503 | SRR13257988 |
| ML66_Red | MLBS | 7-Jul-2018 | Tn5 | Illumina PE150 | 7596370 | SRR13257989 |
| ML67_Red | MLBS | 7-Jul-2018 | Tn5 | Illumina PE150 | 7351721 | SRR13257990 |
| ML68_Red | MLBS | 7-Jul-2018 | Tn5 | Illumina PE150 | 6448176 | SRR13257991 |
| ML69_Red | MLBS | 7-Jul-2018 | Tn5 | Illumina PE150 | 7633376 | SRR13257992 |
| ML70_Red | MLBS | 7-Jul-2018 | Tn5 | Illumina PE150 | 8179649 | SRR13257993 |
| ML9_Red | MLBS | 7-Jul-2018 | Tn5 | Illumina PE150 | 6874469 | SRR13257995 |
| MLR32 | Man Lu Ridge Rd., Maryland | 16-May-2017 | Tn5 | Illumina PE150 | 5102731 | SRR13257996 |
| MLR34 | Man Lu Ridge Rd., Maryland | 16-May-2017 | Tn5 | Illumina PE150 | 6204711 | SRR13257997 |
| MLR43 | Man Lu Ridge Rd., Maryland | 16-May-2017 | Tn5 | Illumina PE150 | 7370217 | SRR13257998 |
| MLR49 | Man Lu Ridge Rd., Maryland | 16-May-2017 | Tn5 | Illumina PE150 | 6120506 | SRR13258057 |
| OWR58 | Old Waterford Road, Leesburg, VA | 22-May-2017 | Tn5 | Illumina PE150 | 7305751 | SRR13258058 |

**Table STAR Methods 6.** Details of biological samples, sequence files, and SRA accession numbers for H. hamamelidis targeted genome re-sequencing of approximately 800 kbp of Chromosome 1. MLBS = Mountain Lakes Biological Station, Pembroke, Virginia.

| Sample # | Collection Location | Collection Date | Library Prep Method | Sequencing Method | # Read pairs | Accession # |
| --- | --- | --- | --- | --- | --- | --- |
| GCC77 | Goose Creek Lane, Leesburg, Virginia | 31-May-2017 | Tn5 | Illumina PE150 | 314257 | SRR13258059 |
| ML10_Gr | MLBS | 7-Jul-2018 | Tn5 | Illumina PE150 | 334744 | SRR13258060 |
| ML14_Gr | MLBS | 7-Jul-2018 | Tn5 | Illumina PE150 | 325866 | SRR13258061 |
| ML15 Gr | MLBS | 7-Jul-2018 | Tn5 | Illumina<br>PE150 | 298479 | SRR13258062 |
| ML17 Gr | MLBS | 7-Jul-2018 | Tn5 | Illumina<br>PE150 | 313916 | SRR13258152 |
| ML18 Gr | MLBS | 7-Jul-2018 | Tn5 | Illumina<br>PE150 | 290694 | SRR13258151 |
| ML2 Gr | MLBS | 7-Jul-2018 | Tn5 | Illumina<br>PE150 | 331929 | SRR13258150 |
| ML21 Gr | MLBS | 7-Jul-2018 | Tn5 | Illumina<br>PE150 | 329166 | SRR13258149 |
| ML23 Gr | MLBS | 7-Jul-2018 | Tn5 | Illumina<br>PE150 | 327831 | SRR13258148 |
| ML24 Gr | MLBS | 7-Jul-2018 | Tn5 | Illumina<br>PE150 | 178226 | SRR13258147 |
| ML27 Gr | MLBS | 7-Jul-2018 | Tn5 | Illumina<br>PE150 | 275723 | SRR13258146 |
| ML28 Gr | MLBS | 7-Jul-2018 | Tn5 | Illumina<br>PE150 | 346212 | SRR13258145 |
| ML30 Gr | MLBS | 7-Jul-2018 | Tn5 | Illumina<br>PE150 | 300476 | SRR13258144 |
| ML33 Gr | MLBS | 7-Jul-2018 | Tn5 | Illumina<br>PE150 | 235769 | SRR13258143 |
| ML34 Gr | MLBS | 7-Jul-2018 | Tn5 | Illumina<br>PE150 | 313858 | SRR13258064 |
| ML36 Gr | MLBS | 7-Jul-2018 | Tn5 | Illumina<br>PE150 | 6195 | SRR13258142 |
| ML38 Gr | MLBS | 7-Jul-2018 | Tn5 | Illumina<br>PE150 | 312165 | SRR13258141 |
| ML40 Gr | MLBS | 7-Jul-2018 | Tn5 | Illumina<br>PE150 | 262504 | SRR13258140 |
| ML41 Gr | MLBS | 7-Jul-2018 | Tn5 | Illumina<br>PE150 | 266728 | SRR13258065 |
| ML43 Gr | MLBS | 7-Jul-2018 | Tn5 | Illumina<br>PE150 | 334740 | SRR13258066 |
| ML44 Gr | MLBS | 7-Jul-2018 | Tn5 | Illumina<br>PE150 | 355990 | SRR13258139 |
| ML48 Gr | MLBS | 7-Jul-2018 | Tn5 | Illumina<br>PE150 | 76870 | SRR13258138 |
| ML49 Gr | MLBS | 7-Jul-2018 | Tn5 | Illumina<br>PE150 | 219732 | SRR13258137 |
| ML50 Gr | MLBS | 7-Jul-2018 | Tn5 | Illumina<br>PE150 | 319064 | SRR13258067 |
| ML51 Gr | MLBS | 7-Jul-2018 | Tn5 | Illumina<br>PE150 | 335990 | SRR13258069 |
| ML53 Gr | MLBS | 7-Jul-2018 | Tn5 | Illumina<br>PE150 | 240432 | SRR13258136 |
| ML55 Gr | MLBS | 7-Jul-2018 | Tn5 | Illumina<br>PE150 | 334902 | SRR13258135 |
| ML57 Gr | MLBS | 7-Jul-2018 | Tn5 | Illumina<br>PE150 | 89944 | SRR13258134 |
| ML6 Gr | MLBS | 7-Jul-2018 | Tn5 | Illumina<br>PE150 | 340415 | SRR13258070 |
| ML60 Gr | MLBS | 7-Jul-2018 | Tn5 | Illumina<br>PE150 | 283159 | SRR13258071 |
| ML61 Gr | MLBS | 7-Jul-2018 | Tn5 | Illumina<br>PE150 | 317924 | SRR13258133 |
| ML62 Gr | MLBS | 7-Jul-2018 | Tn5 | Illumina<br>PE150 | 312738 | SRR13258132 |
| ML65 Gr | MLBS | 7-Jul-2018 | Tn5 | Illumina<br>PE150 | 332910 | SRR13258131 |
| ML70 Gr | MLBS | 7-Jul-2018 | Tn5 | Illumina<br>PE150 | 313079 | SRR13258072 |
| ML71 Gr | MLBS | 7-Jul-2018 | Tn5 | Illumina<br>PE150 | 353342 | SRR13258074 |
| ML73 Gr | MLBS | 7-Jul-2018 | Tn5 | Illumina<br>PE150 | 309115 | SRR13258075 |
| ML77 Gr | MLBS | 7-Jul-2018 | Tn5 | Illumina<br>PE150 | 374346 | SRR13258076 |
| ML79 Gr | MLBS | 7-Jul-2018 | Tn5 | Illumina<br>PE150 | 367693 | SRR13258077 |
| ML82 Gr | MLBS | 7-Jul-2018 | Tn5 | Illumina<br>PE150 | 280482 | SRR13258078 |
| MLR38 | Man Lu Ridge Rd.,<br>Maryland | 16-May-2017 | Tn5 | Illumina<br>PE150 | 331238 | SRR13258079 |
| MLR40 | Man Lu Ridge Rd.,<br>Maryland | 16-May-2017 | Tn5 | Illumina<br>PE150 | 356281 | SRR13258080 |
| MLR50 | Man Lu Ridge Rd.,<br>Maryland | 16-May-2017 | Tn5 | Illumina<br>PE150 | 290972 | SRR13258081 |
| MLR51 | Man Lu Ridge Rd.,<br>Maryland | 16-May-2017 | Tn5 | Illumina<br>PE150 | 327369 | SRR13258082 |
| MLR52 | Man Lu Ridge Rd.,<br>Maryland | 16-May-2017 | Tn5 | Illumina<br>PE150 | 327149 | SRR13258083 |
| MLR53 | Man Lu Ridge Rd.,<br>Maryland | 16-May-2017 | Tn5 | Illumina<br>PE150 | 323702 | SRR13258085 |
| MLR54 | Man Lu Ridge Rd.,<br>Maryland | 16-May-2017 | Tn5 | Illumina<br>PE150 | 285992 | SRR13258086 |
| OWR57 | Old Waterford<br>Road, Leesburg,<br>VA | 22-May-2017 | Tn5 | Illumina<br>PE150 | 306545 | SRR13258087 |
| OWR70 | Old Waterford<br>Road, Leesburg,<br>VA | 30-May-2017 | Tn5 | Illumina<br>PE150 | 302774 | SRR13258088 |
| ML10 Red | MLBS | 7-Jul-2018 | Tn5 | Illumina<br>PE150 | 329939 | SRR13258089 |
| ML12 Red | MLBS | 7-Jul-2018 | Tn5 | Illumina<br>PE150 | 3091 | SRR13258090 |
| ML13 Red | MLBS | 7-Jul-2018 | Tn5 | Illumina<br>PE150 | 365121 | SRR13258091 |
| ML16 Red | MLBS | 7-Jul-2018 | Tn5 | Illumina<br>PE150 | 402478 | SRR13258092 |
| ML18 Red | MLBS | 7-Jul-2018 | Tn5 | Illumina<br>PE150 | 395314 | SRR13258093 |
| ML19 Red | MLBS | 7-Jul-2018 | Tn5 | Illumina<br>PE150 | 369502 | SRR13258156 |
| ML2 Red | MLBS | 7-Jul-2018 | Tn5 | Illumina<br>PE150 | 323168 | SRR13258095 |
| ML20 Red | MLBS | 7-Jul-2018 | Tn5 | Illumina<br>PE150 | 360596 | SRR13258096 |
| ML22 Red | MLBS | 7-Jul-2018 | Tn5 | Illumina<br>PE150 | 380225 | SRR13258097 |
| ML23 Red | MLBS | 7-Jul-2018 | Tn5 | Illumina<br>PE150 | 355840 | SRR13258098 |
| ML24 Red | MLBS | 7-Jul-2018 | Tn5 | Illumina<br>PE150 | 360261 | SRR13258157 |
| ML25 Red | MLBS | 7-Jul-2018 | Tn5 | Illumina<br>PE150 | 381256 | SRR13258158 |
| ML27 Red | MLBS | 7-Jul-2018 | Tn5 | Illumina<br>PE150 | 354745 | SRR13258099 |
| ML29 Red | MLBS | 7-Jul-2018 | Tn5 | Illumina<br>PE150 | 360523 | SRR13258100 |
| ML3 Red | MLBS | 7-Jul-2018 | Tn5 | Illumina<br>PE150 | 341628 | SRR13258101 |
| ML32 Red | MLBS | 7-Jul-2018 | Tn5 | Illumina<br>PE150 | 332324 | SRR13258159 |
| ML35 Red | MLBS | 7-Jul-2018 | Tn5 | Illumina<br>PE150 | 353534 | SRR13258103 |
| ML36 Red | MLBS | 7-Jul-2018 | Tn5 | Illumina<br>PE150 | 369572 | SRR13258104 |
| ML37 Red | MLBS | 7-Jul-2018 | Tn5 | Illumina<br>PE150 | 349811 | SRR13258105 |
| ML38 Red | MLBS | 7-Jul-2018 | Tn5 | Illumina<br>PE150 | 393709 | SRR13258106 |
| ML39 Red | MLBS | 7-Jul-2018 | Tn5 | Illumina<br>PE150 | 304153 | SRR13258107 |
| ML41 Red | MLBS | 7-Jul-2018 | Tn5 | Illumina<br>PE150 | 312756 | SRR13258108 |
| ML42 Red | MLBS | 7-Jul-2018 | Tn5 | Illumina<br>PE150 | 281446 | SRR13258109 |
| ML44 Red | MLBS | 7-Jul-2018 | Tn5 | Illumina<br>PE150 | 360792 | SRR13258110 |
| ML45 Red | MLBS | 7-Jul-2018 | Tn5 | Illumina<br>PE150 | 331967 | SRR13258111 |
| ML46 Red | MLBS | 7-Jul-2018 | Tn5 | Illumina<br>PE150 | 306892 | SRR13258112 |
| ML47 Red | MLBS | 7-Jul-2018 | Tn5 | Illumina<br>PE150 | 312341 | SRR13258114 |
| ML52 Red | MLBS | 7-Jul-2018 | Tn5 | Illumina<br>PE150 | 355827 | SRR13258115 |
| ML54 Red | MLBS | 7-Jul-2018 | Tn5 | Illumina<br>PE150 | 220263 | SRR13258116 |
| ML56 Red | MLBS | 7-Jul-2018 | Tn5 | Illumina<br>PE150 | 341262 | SRR13258117 |
| ML57 Red | MLBS | 7-Jul-2018 | Tn5 | Illumina<br>PE150 | 287334 | SRR13258118 |
| ML58 Red | MLBS | 7-Jul-2018 | Tn5 | Illumina<br>PE150 | 337310 | SRR13258119 |
| ML59 Red | MLBS | 7-Jul-2018 | Tn5 | Illumina<br>PE150 | 342148 | SRR13258120 |
| ML62 Red | MLBS | 7-Jul-2018 | Tn5 | Illumina<br>PE150 | 360053 | SRR13257999 |
| ML63 Red | MLBS | 7-Jul-2018 | Tn5 | Illumina<br>PE150 | 303988 | SRR13258000 |
| ML64 Red | MLBS | 7-Jul-2018 | Tn5 | Illumina<br>PE150 | 326164 | SRR13258001 |
| ML65 Red | MLBS | 7-Jul-2018 | Tn5 | Illumina<br>PE150 | 308072 | SRR13258003 |
| ML66 Red | MLBS | 7-Jul-2018 | Tn5 | Illumina<br>PE150 | 334547 | SRR13258004 |
| ML67 Red | MLBS | 7-Jul-2018 | Tn5 | Illumina<br>PE150 | 313769 | SRR13258005 |
| ML68 Red | MLBS | 7-Jul-2018 | Tn5 | Illumina<br>PE150 | 285472 | SRR13258006 |
| ML69 Red | MLBS | 7-Jul-2018 | Tn5 | Illumina<br>PE150 | 335029 | SRR13258007 |
| ML70 Red | MLBS | 7-Jul-2018 | Tn5 | Illumina<br>PE150 | 368484 | SRR13258008 |
| ML9 Red | MLBS | 7-Jul-2018 | Tn5 | Illumina<br>PE150 | 311326 | SRR13258009 |
| MLR32 | Man Lu Ridge Rd.,<br>Maryland | 16-May-2017 | Tn5 | Illumina<br>PE150 | 206644 | SRR13258010 |
| MLR34 | Man Lu Ridge Rd.,<br>Maryland | 16-May-2017 | Tn5 | Illumina<br>PE150 | 271798 | SRR13258011 |
| MLR43 | Man Lu Ridge Rd.,<br>Maryland | 16-May-2017 | Tn5 | Illumina<br>PE150 | 320317 | SRR13258130 |
| MLR49 | Man Lu Ridge Rd.,<br>Maryland | 16-May-2017 | Tn5 | Illumina<br>PE150 | 256744 | SRR13258013 |
| OWR58 | Old Waterford<br>Road, Leesburg,<br>VA | 22-May-2017 | Tn5 | Illumina<br>PE150 | 320996 | SRR13258014 |

**Table STAR Methods 7.** Details of biological samples, sequence files, and SRA accession numbers for *Hormaphis cornu* salivary gland RNA-sequencing from fundatrices from red and green galls. JRC = Janelia Research Campus.

| Sample ID | Collection Location | Collection Date | Gall color | Library Prep Method | Sequencing Method | # Read pairs | Accession # |
| --- | --- | --- | --- | --- | --- | --- | --- |
| AKG_4 | JRC | 28-Apr-2019 | Green | WGB | Illumina PE 150 | 11607393 | SRR11428726 |
| DSG_6 | JRC | 28-Apr-2019 | Green | WGB | Illumina PE 150 | 12179326 | SRR11428724 |
| DSG_7 | JRC | 30-Apr-2019 | Green | WGB | Illumina PE 150 | 13321660 | SRR11428723 |
| DSG_8 | JRC | 30-Apr-2019 | Green | WGB | Illumina PE 150 | 10433415 | SRR11428722 |
| DSG_9 | JRC | 30-Apr-2019 | Green | WGB | Illumina PE 150 | 14776468 | SRR11428721 |
| DSG_10 | JRC | 30-Apr-2019 | Green | WGB | Illumina PE 150 | 13105789 | SRR11428720 |
| DSG_11 | JRC | 01-May-2019 | Green | WGB | Illumina PE 150 | 12867296 | SRR11428719 |
| DSG_13 | JRC | 01-May-2019 | Green | WGB | Illumina PE 150 | 10772840 | SRR11428718 |
| DSR_11 | JRC | 01-May-2019 | Red | WGB | Illumina PE 150 | 15042624 | SRR11428717 |
| DSR_12 | JRC | 01-May-2019 | Red | WGB | Illumina PE 150 | 11051413 | SRR11428716 |
| DSR_13 | JRC | 01-May-2019 | Red | WGB | Illumina PE 150 | 15396180 | SRR11428715 |
| DSR_15 | JRC | 01-May-2019 | Red | WGB | Illumina PE 150 | 18975841 | SRR11428784 |
| DSR_16 | JRC | 01-May-2019 | Red | WGB | Illumina PE 150 | 22465588 | SRR11428782 |
| DSR_17 | JRC | 01-May-2019 | Red | WGB | Illumina PE 150 | 15618318 | SRR11428781 |
| DSR_18 | JRC | 01-May-2019 | Red | WGB | Illumina PE 150 | 16229009 | SRR11428780 |
| AKR1 | JRC | 28-Apr-2019 | Red | WGB | Illumina PE 150 | 14353107 | SRR13258428 |
| AKR2 | JRC | 28-Apr-2019 | Red | WGB | Illumina PE 150 | 17302776 | SRR13258427 |
| AKR3 | JRC | 28-Apr-2019 | Red | WGB | Illumina PE 150 | 24214993 | SRR13258426 |
| AKR4 | JRC | 28-Apr-2019 | Red | WGB | Illumina PE 150 | 12914802 | SRR13258425 |
| DSR5 | JRC | 28-Apr-2019 | Red | WGB | Illumina PE 150 | 16153051 | SRR13258424 |
| DSR6 | JRC | 28-Apr-2019 | Red | WGB | Illumina PE 150 | 10998399 | SRR13258423 |
| DSR7 | JRC | 28-Apr-2019 | Red | WGB | Illumina PE 150 | 10264369 | SRR13258422 |
| DSR_8 | JRC | 28-Apr-2019 | Red | WGB | Illumina PE 150 | 19769187 | SRR13258421 |
| DSR_9 | JRC | 28-Apr-2019 | Red | WGB | Illumina PE 150 | 18487599 | SRR13258420 |
| DSR | JRC | 28-Apr-2019 | Red | WGB | Illumina PE 150 | 9649227 | SRR13258419 |

**Table STAR Methods 8.** Details of biological samples, sequence files, and SRA accession numbers for Hamamelis virginiana RNA-sequencing from galls and leaf around galls. JRC = Janelia Research Campus, Ashburn, VA.

| Sample ID | Collection Location | Collection Date | Library prep method | Sequencing method | # Read pairs | Accession # |
| --- | --- | --- | --- | --- | --- | --- |
| Gall_9 | JRC | 11-Apr-2017 | Nugen Universal Plus mRNA-seq | Illumina PE150 | 14453904 | SRR11362686 |
| Gall_11 | JRC | 11-Apr-2017 | Nugen Universal Plus mRNA-seq | Illumina PE150 | 7224527 | SRR11362685 |
| Early_gall_18 | JRC | 12-Apr-2017 | Nugen Universal Plus mRNA-seq | Illumina PE150 | 25279196 | SRR11362642 |
| Gall_20 | JRC | 12-Apr-2017 | Nugen Universal Plus mRNA-seq | Illumina PE150 | 11692754 | SRR11362631 |
| Gall_22 | JRC | 12-Apr-2017 | Nugen Universal Plus mRNA-seq | Illumina PE150 | 10990345 | SRR11362620 |
| Gall_26 | JRC | 14-Apr-2017 | Nugen Universal Plus mRNA-seq | Illumina PE150 | 6419861 | SRR13258463 |
| Gall_27 | JRC | 14-Apr-2017 | Nugen Universal Plus mRNA-seq | Illumina PE150 | 7086413 | SRR13258462 |
| Gall_28 | JRC | 14-Apr-2017 | Nugen Universal Plus mRNA-seq | Illumina PE150 | 7618914 | SRR13258461 |
| Gall_29 | JRC | 14-Apr-2017 | Nugen Universal Plus mRNA-seq | Illumina PE150 | 5633325 | SRR13258460 |
| Gall_31 | JRC | 14-Apr-2017 | Nugen Universal Plus mRNA-seq | Illumina PE150 | 12324002 | SRR11362672 |
| Early_gall_33 | JRC | 14-Apr-2017 | Nugen Universal Plus mRNA-seq | Illumina PE150 | 31823857 | SRR11362661 |
| gall_35 | JRC | 14-Apr-2017 | Nugen Universal Plus mRNA-seq | Illumina PE150 | 11104190 | SRR11362652 |
| Gall_42 | JRC | 15-Apr-2017 | Nugen Universal Plus mRNA-seq | Illumina PE150 | 6199468 | SRR13258459 |
| Gall_44 | JRC | 15-Apr-2017 | Nugen Universal Plus mRNA-seq | Illumina PE150 | 11104190 | SRR11362683 |
| Gall_46 | JRC | 15-Apr-2017 | Nugen Universal Plus mRNA-seq | Illumina PE150 | 11965583 | SRR11362684 |
| Gall_47 | JRC | 15-Apr-2017 | Nugen Universal Plus mRNA-seq | Illumina PE150 | 18509404 | SRR11362682 |
| Bottom_gall_51 | JRC | 17-Apr-2017 | Nugen Universal Plus mRNA-seq | Illumina PE150 | 13291659 | SRR11362650 |
| Top_gall_52 | JRC | 17-Apr-2017 | Nugen Universal Plus mRNA-seq | Illumina PE150 | 11679869 | SRR11362649 |
| Bottom_gall_53 | JRC | 17-Apr-2017 | Nugen Universal Plus mRNA-seq | Illumina PE150 | 11661968 | SRR11362648 |
| Top_gall_54 | JRC | 17-Apr-2017 | Nugen Universal Plus mRNA-seq | Illumina PE150 | 12277949 | SRR11362647 |
| Bottom_gall_55 | JRC | 17-Apr-2017 | Nugen Universal Plus mRNA-seq | Illumina PE150 | 10705594 | SRR11362646 |
| Top_gall_56 | JRC | 17-Apr-2017 | Nugen Universal Plus mRNA-seq | Illumina PE150 | 12245151 | SRR11362645 |
| Bottom_gall_57 | JRC | 17-Apr-2017 | Nugen Universal Plus mRNA-seq | Illumina PE150 | 5688881 | SRR11362644 |
| Top_gall_58 | JRC | 17-Apr-2017 | Nugen Universal Plus mRNA-seq | Illumina PE150 | 8195240 | SRR11362643 |
| Gall_60 | JRC | 17-Apr-2017 | Nugen Universal Plus mRNA-seq | Illumina PE150 | 7075731 | SRR13258458 |
| Gall_61 | JRC | 17-Apr-2017 | Nugen Universal Plus mRNA-seq | Illumina PE150 | 5226728 | SRR13258472 |
| Gall_62 | JRC | 17-Apr-2017 | Nugen Universal Plus mRNA-seq | Illumina PE150 | 3985629 | SRR13258471 |
| Gall_63 | JRC | 17-Apr-2017 | Nugen Universal Plus mRNA-seq | Illumina PE150 | 6194899 | SRR13258470 |
| Gall_65 | JRC | 17-Apr-2017 | Nugen Universal Plus mRNA-seq | Illumina PE150 | 10541245 | SRR13258469 |
| Gall_66 | JRC | 17-Apr-2017 | Nugen Universal Plus mRNA-seq | Illumina PE150 | 5324437 | SRR13258468 |
| Apical_gall_118 | JRC | 22-Apr-2017 | Nugen Universal Plus mRNA-seq | Illumina PE150 | 8305160 | SRR13258474 |
| Apical_gall_122 | JRC | 22-Apr-2017 | Nugen Universal Plus mRNA-seq | Illumina PE150 | 3971025 | SRR13258473 |
| Basal_gall_116 | JRC | 22-Apr-2017 | Nugen Universal Plus mRNA-seq | Illumina PE150 | 10405803 | SRR13258465 |
| Basal_gall_120 | JRC | 22-Apr-2017 | Nugen Universal Plus mRNA-seq | Illumina PE150 | 4887393 | SRR13258464 |
| Medial_gall_117 | JRC | 22-Apr-2017 | Nugen Universal Plus mRNA-seq | Illumina PE150 | 3452858 | SRR13258467 |
| Medial_gall_121 | JRC | 22-Apr-2017 | Nugen Universal Plus mRNA-seq | Illumina PE150 | 5465679 | SRR13258466 |
| Leaf_around_gall_10 | JRC | 11-Apr-2017 | Nugen Universal Plus mRNA-seq | Illumina PE150 | 3055859 | SRR11362625 |
| Leaf_around_gall_12 | JRC | 11-Apr-2017 | Nugen Universal Plus mRNA-seq | Illumina PE150 | 2924715 | SRR11362624 |
| Leaf_around_gall_19 | JRC | 11-Apr-2017 | Nugen Universal Plus mRNA-seq | Illumina PE150 | 39720279 | SRR11362623 |
| Leaf_around_gall_21 | JRC | 12-Apr-2017 | Nugen Universal Plus mRNA-seq | Illumina PE150 | 4346238 | SRR11362622 |
| Leaf_around_gall_23 | JRC | 12-Apr-2017 | Nugen Universal Plus mRNA-seq | Illumina PE150 | 9527411 | SRR11362621 |
| Leaf_around_gall_30 | JRC | 14-Apr-2017 | Nugen Universal Plus mRNA-seq | Illumina PE150 | 10162178 | SRR11362619 |
| Leaf_around_gall_34 | JRC | 14-Apr-2017 | Nugen Universal Plus mRNA-seq | Illumina PE150 | 34938887 | SRR11362681 |
| Leaf_around_gall_38 | JRC | 14-Apr-2017 | Nugen Universal Plus mRNA-seq | Illumina PE150 | 5195676 | SRR11362680 |
| Leaf_around_gall_43 | JRC | 15-Apr-2017 | Nugen Universal Plus mRNA-seq | Illumina PE150 | 11911791 | SRR11362679 |
| Leaf_around_gall_45 | JRC | 15-Apr-2017 | Nugen Universal Plus mRNA-seq | Illumina PE150 | 9012995 | SRR11362678 |
| Leaf_around_gall_49 | JRC | 15-Apr-2017 | Nugen Universal Plus mRNA-seq | Illumina PE150 | 2405452 | SRR11362677 |
| Leaf_around_gall_50 | JRC | 17-Apr-2017 | Nugen Universal Plus mRNA-seq | Illumina PE150 | 4591989 | SRR11362676 |
| Leaf_around_gall_59 | JRC | 17-Apr-2017 | Nugen Universal Plus mRNA-seq | Illumina PE150 | 6517483 | SRR11362675 |
| Leaf_around_gall_64 | JRC | 17-Apr-2017 | Nugen Universal Plus mRNA-seq | Illumina PE150 | 4871569 | SRR11362674 |
| Leaf_around_gall_67 | JRC | 17-Apr-2017 | Nugen Universal Plus mRNA-seq | Illumina PE150 | 4352513 | SRR11362673 |
| Leaf_around_gall_119 | JRC | 22-Apr-2017 | Nugen Universal Plus mRNA-seq | Illumina PE150 | 4505696 | <u>SRR11362671</u> |
| Leaf_around_gall_123 | JRC | 22-Apr-2017 | Nugen Universal Plus mRNA-seq | Illumina PE150 | 2655968 | <u>SRR11362670</u> |

**Table STAR Methods 9.** Details of biological samples, sequence files, and SRA accession numbers for *Hamamelis virginiana* RNA-sequencing from green and red galls.

| Sample ID | Collection Location | Collection Date | Leaf Sample | Gall color | Library Prep Method | Sequencing Method | # Read pairs | Accession # |
| --- | --- | --- | --- | --- | --- | --- | --- | --- |
| 20JRC105R | JRC | 08-May-2020 | 1 | Red | WGB | Illumina SE75 | 5813855 | SRR13258378 |
| 20JRC104G | JRC | 08-May-2020 | 1 | Green | WGB | Illumina SE75 | 6618273 | SRR13258369 |
| 20JRC107R | JRC | 08-May-2020 | 2 | Red | WGB | Illumina SE75 | 7330505 | SRR13258344 |
| 20JRC106G | JRC | 08-May-2020 | 2 | Green | WGB | Illumina SE75 | 6999471 | SRR13258368 |
| 20JRC108R | JRC | 08-May-2020 | 3 | Red | WGB | Illumina SE75 | 7444078 | SRR13258376 |
| 20JRC109G | JRC | 08-May-2020 | 3 | Green | WGB | Illumina SE75 | 7711014 | SRR13258350 |
| 20JRC110R | JRC | 08-May-2020 | 4 | Red | WGB | Illumina SE75 | 6961512 | SRR13258372 |
| 20JRC111G | JRC | 09-May-2020 | 4 | Green | WGB | Illumina SE75 | 6765543 | SRR13258346 |
| 20JRC113R | JRC | 08-May-2020 | 5 | Red | WGB | Illumina SE75 | 6707106 | SRR13258371 |
| 20JRC112G | JRC | 09-May-2020 | 5 | Green | WGB | Illumina SE75 | 7274181 | SRR13258345 |
| 20JRC116R | JRC | 08-May-2020 | 7 | Red | WGB | Illumina SE75 | 12806892 | SRR13258370 |
| 20JRC1076R | JRC | 08-May-2020 | 8 | Red | WGB | Illumina SE75 | 9172629 | SRR13258377 |
| 20JRC1077R | JRC | 08-May-2020 | 8 | Red | WGB | Illumina SE75 | 9280201 | SRR13258366 |
| 20JRC1074G | JRC | 08-May-2020 | 8 | Green | WGB | Illumina SE75 | 9200050 | SRR13258367 |
| 20JRC1075G | JRC | 08-May-2020 | 8 | Green | WGB | Illumina SE75 | 9001911 | SRR13258365 |
| 20JRC1079R | JRC | 08-May-2020 | 9 | Red | WGB | Illumina SE75 | 8765890 | SRR13258355 |
| 20JRC1078G | JRC | 08-May-2020 | 9 | Green | WGB | Illumina SE75 | 8534470 | SRR13258364 |
| 20JRC1081R | JRC | 08-May-2020 | 10 | Red | WGB | Illumina SE75 | 7555292 | SRR13258343 |
| 20JRC1080G | JRC | 08-May-2020 | 10 | Green | WGB | Illumina SE75 | 8840709 | SRR13258363 |
| 20JRC1083R | JRC | 08-May-2020 | 11 | Red | WGB | Illumina SE75 | 9657276 | SRR13258342 |
| 20JRC1082G | JRC | 08-May-2020 | 11 | Green | WGB | Illumina SE75 | 8984115 | SRR13258362 |
| 20JRC1085R | JRC | 08-May-2020 | 12 | Red | WGB | Illumina SE75 | 9542403 | SRR13258341 |
| 20JRC1086R | JRC | 08-May-2020 | 13 | Red | WGB | Illumina SE75 | 9508095 | SRR13258340 |
| 20JRC1087G | JRC | 08-May-2020 | 13 | Green | WGB | Illumina SE75 | 9552857 | SRR13258361 |
| 20JRC1089R | JRC | 08-May-2020 | 14 | Red | WGB | Illumina SE75 | 8580538 | SRR13258339 |
| 20JRC1088G | JRC | 08-May-2020 | 14 | Green | WGB | Illumina SE75 | 9348175 | SRR13258360 |
| 20JRC1091R | JRC | 08-May-2020 | 15 | Red | WGB | Illumina SE75 | 10193238 | SRR13258375 |
| 20JRC1090G | JRC | 09-May-2020 | 15 | Green | WGB | Illumina SE75 | 9782456 | SRR13258359 |
| 20JRC1092G | JRC | 09-May-2020 | 15 | Green | WGB | Illumina SE75 | 10354726 | SRR13258358 |
| 20JRC1093R | JRC | 08-May-2020 | 16 | Red | WGB | Illumina SE75 | 10632771 | SRR13258374 |
| 20JRC1094G | JRC | 09-May-2020 | 16 | Green | WGB | Illumina SE75 | 9767489 | SRR13258357 |
| 20JRC1095G | JRC | 09-May-2020 | 16 | Green | WGB | Illumina SE75 | 9957641 | SRR13258356 |
| 20JRC1096G | JRC | 09-May-2020 | 16 | Green | WGB | Illumina SE75 | 9938437 | SRR13258354 |
| 20JRC1097G | JRC | 09-May-2020 | 16 | Green | WGB | Illumina SE75 | 8222414 | SRR13258353 |
| 20JRC1100R | JRC | 08-May-2020 | 17 | Red | WGB | Illumina SE75 | 5354681 | SRR13258373 |
| 20JRC1098G | JRC | 09-May-2020 | 17 | Green | WGB | Illumina SE75 | 5890682 | SRR13258352 |
| 20JRC1099G | JRC | 09-May-2020 | 17 | Green | WGB | Illumina SE75 | 6057780 | SRR13258351 |
| 20JRC1101G | JRC | 09-May-2020 | 17 | Green | WGB | Illumina SE75 | 7237058 | SRR13258349 |
| 20JRC1102G | JRC | 08-May-2020 | 18 | Green | WGB | Illumina SE75 | 4943324 | SRR13258348 |
| 20JRC1103G | JRC | 08-May-2020 | 18 | Green | WGB | Illumina SE75 | 7407432 | SRR13258347 |

## Method Details

### Imaging of leaves and fundatrices inside developing galls

Young *Hamamelis virginiana* (witch hazel) leaves or leaves with early stage galls of *Hormaphis cornu* were fixed in Phosphate Buffered Saline (PBS: 137 mM NaCl, 2.7 mM KCl, 10 mM Na2HPO4, 1.8 mM KH2PO4, pH 7.4) containing 0.1% Triton X-100, 2% paraformaldehyde and 0.5% glutaraldehyde (paraformaldehyde and glutaraldehyde were EM grade from Electron Microscopy Services) at room temperature for two hours without agitation to prevent the disruption of the aphid stylet inserted into leaf tissue. Fixed leaves or galls were washed in PBS containing 0.1% Triton X-100, hand cut into small sections (∼10 mm2), and embedded in 7% agarose for subsequent sectioning into 0.3 mm thick sections using a Leica Vibratome (VT1000s). Sectioned plant tissue was stained with Calcofluor White (0.1 mg/mL, Sigma-Aldrich, F3543) and Congo Red (0.25 mg/mL, Sigma-Aldrich, C6767) in PBS containing 0.1% Triton X-100 with 0.5% DMSO and Escin (0.05 mg/mL, Sigma-Aldrich, E1378) at room temperature with gentle agitation for 2 days. Stained sections were washed with PBS containing 0.1% Triton X-100. Soft tissues were digested and cleared to reduce light scattering during subsequent imaging using a mixture of 0.25 mg/mL collagenase/dispase (Roche #10269638001) and 0.25 mg/mL hyaluronidase (Sigma Aldrich #H3884) in PBS containing 0.1% Triton X-100 for 5 hours at 37°C. To avoid artifacts and warping caused by osmotic shrinkage of soft tissue and agarose, samples were gradually dehydrated in glycerol (2% to 80%) and then ethanol (20% to 100 %) (Ott 2008) and mounted in methyl salicylate (Sigma-Aldrich, M6752) for imaging. Serial optical sections were obtained at 2 µm intervals on a Zeiss 880 confocal microscope with a Plan-Apochromat 10x/0.45 NA objective, at 1 µm with a LD-LCI 25x/0.8 NA objective or at 0.5 µm with a Plan-Apochromat 40x/0.8 NA objective. Maximum projections of confocal stacks or rotation of images were carried out using FIJI (Schindelin et al. 2012).

### *Hormaphis cornu* genome sequencing, assembly, and annotation

We collected *H. cornu* aphids from a single gall for genome sequencing (Table STAR Methods 1). All aphids within the gall were presumed to be clonal offspring of a single fundatrix, because all *H. cornu* galls we have ever examined contain only a single fundatrix and the ostiole of the galls was closed at the time we collected this gall, so there is little chance of inter-gall migration. High molecular weight (HMW) DNA was prepared by gently grinding aphids with a plastic pestle against the inside wall of a 2 mL Eppendorf tube in 1 mL of 0.5% SDS, 200 mM Tris-HCl pH 8, 25 mM EDTA, 250 mM NaCl with 10 uL of 1 mg/mL RNAse A. Sample was incubated at 37°C for 1 hour and then 30uL of 10 mg/mL ProteinaseK was added and the sample was incubated for an additional 1 hr at 50°C with gentle agitation at 300 RPM. 1 mL of Phenol:Chloroform:Isoamyl alcohol (25:24:1) was added and the sample was centrifuged at 16,000 RCF for 10 min. The supernatant was removed to a new 2mL Eppendorf tube and the Phenol:Chloroform:Isoamyl alcohol extraction was repeated. The supernatant was removed to a new 2 mL tube and 2.5 X volumes of absolute ethanol were added. The sample was centrifuged at 16,000 RCF for 15 min and then washed with fresh 70% ethanol. All ethanol was removed with a pipette and the sample was air dried for approximately 15 minutes and DNA was resuspended in 50 uL TE. This sample was sent to HudsonAlpha Institute for Biotechnology for genome sequencing.

DNA quality control, library preparation, and Chromium 10X linked read sequencing were performed by HudsonAlpha Institute for Biotechnology. Most of the mass of the HMW DNA appeared greater than approximately 50 kb on a pulsed field gel and paired end sequencing on an Illumina HiSeq X10 yielded 816M reads. The genome was assembled using *Supernova* (Weisenfeld et al. 2017) using 175M reads, which generated the best genome N50 of a range of values tested. This 10X genome consisted of 21,072 scaffolds of total length 319.352 MB. The genome scaffold N50 was 839.101 KB and the maximum scaffold length was 3.495 MB.

We then contracted with Dovetail Genomics to apply Chicago (*in vitro* proximity ligation) and HiC (*in vivo* proximity ligation) to generate larger scaffolds (https://dovetailgenomics.com/ga_tech_overview/). We submitted HMW gDNA from the same sample used for 10X genome sequencing for Chicago and a separate sample of frozen aphids for HiC (Table STAR Methods 1). The Dovetail genome consisted of 11,244 scaffolds of total length 320.34 MB with a scaffold N50 of 36.084 Mb. This genome, named hormaphis_cornu_26Sep2017_PoQx8, contains 9 main scaffolds, each longer than 17.934 Mb, which appear to represent the expected 9 chromosomes of *H. cornu* (Blackman and Eastop 1994). This assembly also includes the circular genome of the bacterial endosymbiont *Buchnera aphidicola* of 643,259 bp. Assembly and *BUSCO* analysis statistics (Simao et al. 2015) using the *gVolante* web interface (Nishimura et al., 2017) with the Arthropod gene set are shown below.

| **Assembly statistics** | **hormaphis_cornu_26Sep2017_PoQx8** |
| --- | --- |
| # of scaffolds: | 11242 |
| Total length: | 320,336,030 |
| Longest sequence: | 60,222,264 |
| N50 sequence length: | 36,083,769 |
| Sum length of sequences > 1M | 297,831,669 (93.0% of total length) |
| Sum length of sequences > 10M | 294,347,208 (91.9% of total length) |

| **BUSCO Analysis** | **hormaphis_cornu_26Sep2017_PoQx8** |
| --- | --- |
| Total # of core genes queried | 1066 |
| # of complete core genes detected | 1026 (96.25%) |
| # of complete and partial core genes detected | 1038 (97.37%) |
| # of missing core genes: | 28 (2.63%) |
| Average # of orthologs per core genes: | 1.02 |
| % of detected core genes that have more than 1 ortholog: | 2.34 |

As further checks on the quality of this genome assembly, we examined the K-mer spectra and examined the HiC contact map. The K-mer spectra comparing kmers from the 10X genome sequencing and the Dovetail assembled genome shown below was calculated using KAT (Mapleson et al. 2017). As shown below, there are two peaks for multiplicity 1X, as expected when sequencing a heterozygous individual, and the vast majority of sequence reads map uniquely to the assembled genome 1X, revealing few duplications.

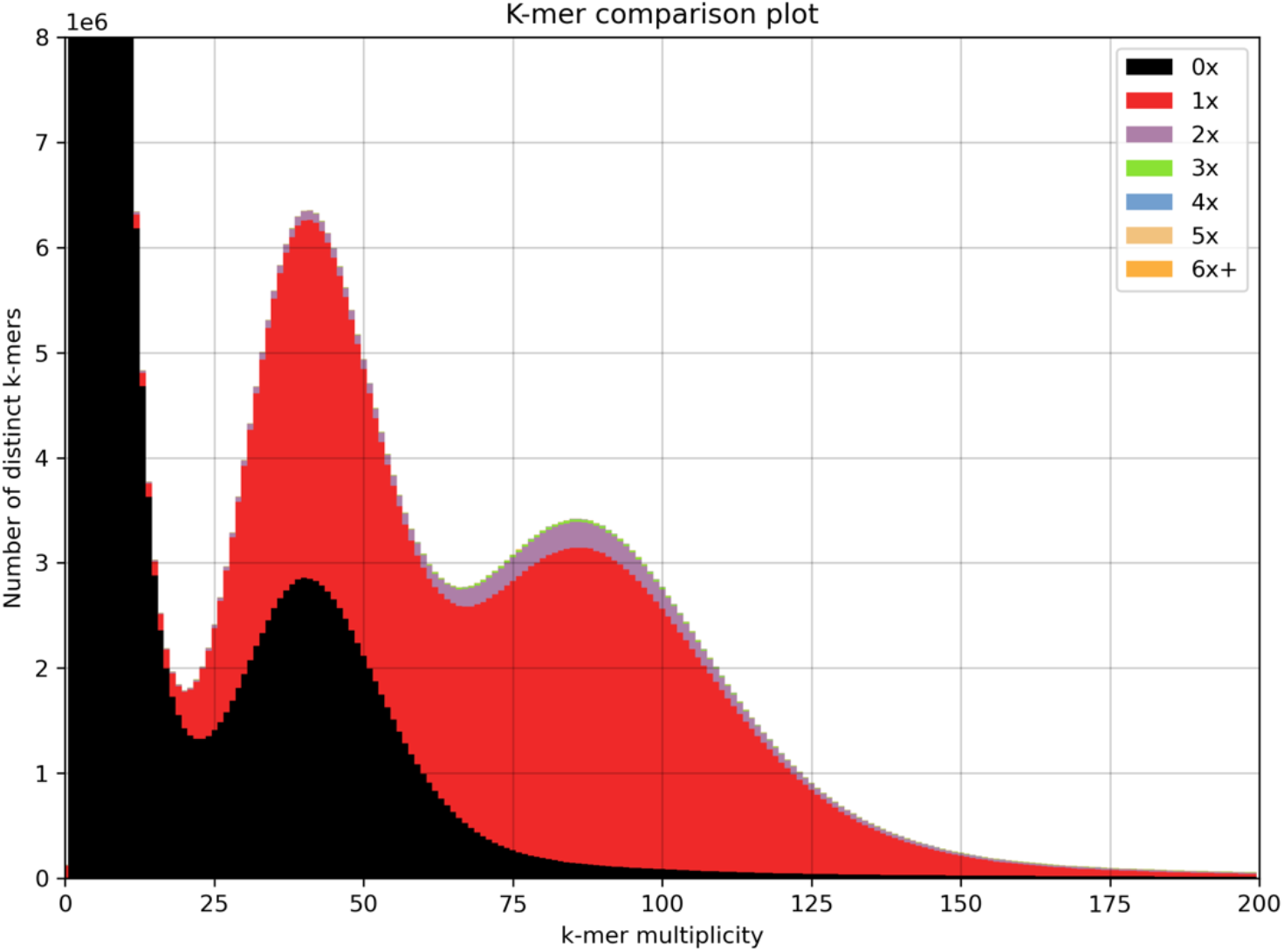

The HiC contact map, shown below, reveals that the great majority of read pairs map to approximately the same location within each of the nine major scaffolds. The X and Y axes indicate the mapping position for the first and second read of each read pair and the color indicates the number of read pairs mapping to each bin. Only scaffolds greater than 1 Mbp are shown.

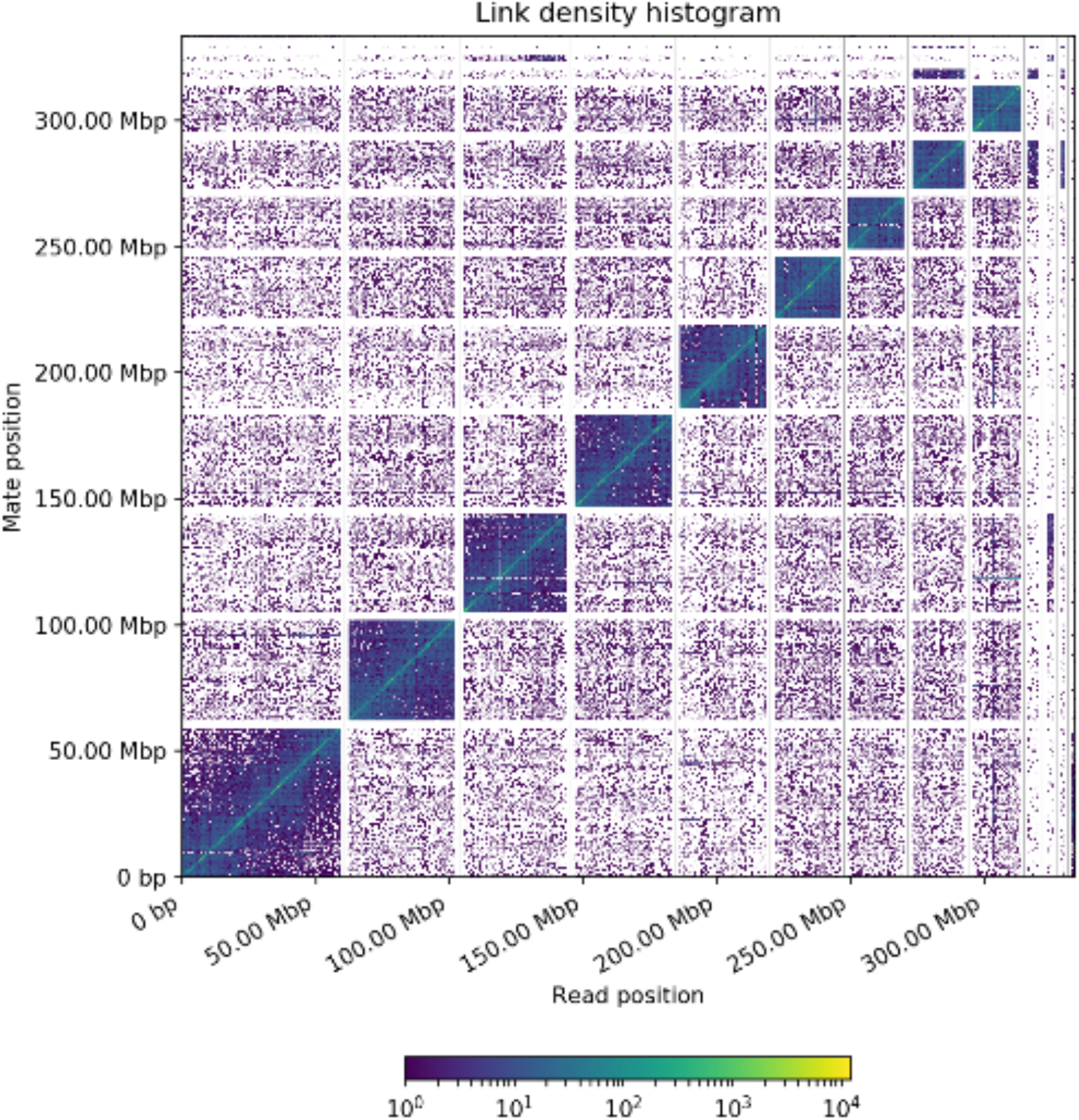

We annotated this genome for protein-coding genes using RNA-seq data collected from salivary glands and carcasses of many stages of the *H. cornu* life cycle (Table STAR Methods 2) using *BRAKER* (Altschul et al. 1990; Barnett et al. 2011; Camacho et al. 2009; Hoff et al. 2016, 2019; Li 2013; Lomsadze, Burns, and Borodovsky 2014; Stanke et al. 2006). To increase the efficiency of mapping RNA-seq reads for differential expression analysis, we predicted 3’ UTRs using *UTRme* (Radío et al. 2018). We found that *UTRme* sometimes predicted UTRs within introns. We therefore applied a custom R script to remove such predicted UTRs. Later, after discovering the *bicycle* genes, we manually annotated all predicted *bicycle* genes, including 5’ and 3’ UTRs, in *APOLLO* (Lewis et al. 2002) by examining evidence from RNAseq reads aligned to the genome with the *Integrative Genomics Viewer* (Robinson et al. 2011; Thorvaldsdóttir, Robinson, and Mesirov 2013). We found that the start sites of many *bicycle* genes were incorrectly annotated by *BRAKER* at a downstream methionine, inadvertently excluding predicted putative signal peptides from these genes. RNAseq evidence often supported transcription start sites that preceded an upstream methionine and these exons were corrected in *APOLLO*. The combined collection of 18,895 automated and 687 manually curated gene models (19,582 total) were used for all subsequent analyses of *H. cornu* genomic data.

### Genome-wide association study of aphids inducing red and green galls

Galls produced by *H. cornu* were collected in the early summer (Table STAR Methods 3) and dissected by making a single vertical cut down the side of each gall with a razor blade to expose the aphids inside. DNA was extracted using the Zymo ZR-96 Quick gDNA kit from the foundress of each gall, because foundresses do not appear to travel between galls. We performed tagmentation of genomic DNA derived from 47 individuals from red galls and 43 from green galls using barcoded adaptors compatible with the Illumina sequencing platform (Hennig et al. 2018). Tagmented samples were pooled without normalization, PCR amplified for 14 cycles, and sequenced on an Illumina NextSeq 500 to generated paired end 150 bp reads to an average depth of 2.9X genomic coverage. The average sequencing depth before filtering was calculated by multiplying the number of read pairs generated by *SAMtools flagstat* version 1.3 (Li et al. 2009) by the read length of 150bp, then dividing by the total genome size (323,839,449bp).

We performed principle component analysis on the genome-wide polymorphism data to detect any potential population structure that might confound GWAS study. Reads were mapped using *bwa mem* version 0.7.17-r1188 (Li 2013) and joint genotyped using *SAMtools mpileup* version 1.3, with the flag *-ugf,* followed by *BCFtools call* version 1.9 (Li 2011), with the flag *-m*. Genotype calls were then filtered for quality and missingness using *BCFtools filter* and *view* version 1.9, where only SNPs with MAF > 0.05, QUAL > 20, and genotyped in at least 80% of the individuals were kept. To limit the number of SNPs for computational efficiency, the SNPs were additionally thinned using *VCFtools –thin* version 0.1.15 to exclude any SNPs within 1000bp of each other. PCA was performed using the *snpgdsPCA* function from the R package *SNPRelate* version 1.20.1 in *R* version 3.6.1 (Zheng et al. 2012).

We performed genome-wide association mapping with these low coverage data by mapping reads with *bwa mem* version 0.7.17-r1188, and then calculated the likelihood of association with gall color with *SAMtools mpileup* version 0.1.19 and *BCFtools view-vcs* version 0.1.19 using BAM files as the input. Association for each SNP was measured by the likelihood-ratio test (LRT) value in the INFO field of the output VCF file, which is a one-degree association test p-value. This method calculates association likelihoods using genotype likelihoods rather than hard genotype calls, ameliorating the issue of low-confidence genotype calls resulting from low-coverage data (Li 2011). The false discovery rate was set as the Bonferroni corrected value for 0.05, which was calculated as 0.05 / 50,957,130 (the total number of SNPs in the genome-wide association mapping).

### Enrichment and sequencing of the genomic region containing highly significant GWAS hits

The low coverage GWAS identified multiple linked SNPS on chromosome 1 that were strongly associated with gall color (Figure 2A). To identify all candidate SNPs in this genomic region and to generate higher-confidence GWAS calls, we enriched this genomic region from a library of pooled tagmented samples of fundatrix DNA from red and green galls using custom designed Arbor Bioscience *MyBaits* for a 800,290 bp region on chromosome 1 spanning the highest scoring GWAS SNP (40,092,625 - 40,892,915 bp). This enriched library was sequenced on an Illumina NextSeq 550 generating paired-end 150 bp reads and resulted in usable re-sequencing data for 48 red gall-producing individuals and 42 green gall-producing individuals, with average pre-filtered sequencing depth of 58.2X.

We mapped reads with *bwa mem* version 0.7.17-r1188 and sorted bam files with *SAMtools sort* version 1.7, marked duplicate reads with *Picard MarkDuplicates* version 2.18.0 (http://broadinstitute.github.io/picard/), re-aligned indels using *GATK IndelRealigner* version 3.4 (McKenna et al. 2010), and called variants using *SAMtools mpileup version* 1.7 and *BCFtools call* version 1.7 (https://github.com/SAMtools/bcftools). This genotyping pipeline is available at https://github.com/YourePrettyGood/PseudoreferencePipeline (thereafter referred to as *PseudoreferencePipeline*). SNPs were quality filtered from the VCF file using *BCFtools view* version 1.7 at DP > 10 and MQ > 40 and merged using *BCFtools merge*.

For PCA analysis, the joint genotype calls were filtered for quality and missingness using *BCFtools filter* and *view* version 1.9, where only SNPs with MAF > 0.05 and genotyped in at least 80% of the individuals were retained. PCA was performed using the *snpgdsPCA* function from the R package *SNPRelate* version 1.20.1 in *Rstudio* version 3.6.1.

Association testing was performed using *PLINK* version 1.90 (Purcell et al. 2007) with minor allele frequency filtered at MAF > 0.2. We did not apply a Hardy-Weinberg equilibrium filter because the samples were not randomly collected from nature. Red galls are rare in our population and we oversampled fundatrices from red galls to roughly match the number of fundatrices sampled from green galls. Results of the GWAS were plotted using the *plotManhattan* function of *Sushi* version 1.24.0 (Phanstiel et al. 2014).

To calculate LD for the 45kbp region surrounding the 11 GWAS SNPs in Fig. S5B, we extracted positions chromosome1:40,475,000 – 40,520,000 from the merged VCF and retained SNPs that both exhibited MAF > 0.05 and were genotyped in at least 80% of the samples using *BCFtools view* and *filter* version 1.9.

To plot LD for the entire target enrichment in Fig. S6D, we filtered the VCF for only SNPs with MAF>0.2 and genotyped in at least 80% of the samples using *bcftools view* and *filter* version 1.9, and thinned the resulting SNP set using *VCFtools–thin* version 0.1.15 to exclude any SNPs within 500bp of each other. We further merged back the 11 significant GWAS SNPs using *bcftools concat*, since the thinning process could have removed one or more of these SNPs. We also removed SNPs in regions where the *H. cornu* reference genome did not align with the genome of the sister species *H. hamamelidis* using *BEDTools intersect* version 2.29.2 (Quinlan and Hall 2010).

The LD heatmaps were generated using the R packages *vcfR* version 1.10.0 (Knaus and Grünwald 2017), *snpStats* version 1.36.0 (Clayton 2019) and *LDheatmap* version 0.99.8 (Shin et al. 2006) in *Rstudio* version 3.6.1. The R code used to generate the LD heatmap figure was adapted from code provided at sfustatgen.github.io/LDheatmap/articles/vcfOnLDheatmap.html. The gene models were plotted using the *plotGenes* function from the R package *Sushi* version 1.24.0.

### Lack of evidence for chromosomal aberrations

To identify possible chromosomal rearrangements or transposable elements that might be linked to the GWAS SNPs, we first trimmed adapters from the *H. cornu* target enrichment data using *Trim Galore!* version 0.6.5 and *cutadapt* version 2.7 (Martin 2011). The trimming pipeline is available at github.com/YourePrettyGood/ParsingPipeline. We then mapped the reads to the *H. cornu* reference genome with *bwa mem* version 0.7.17, sorted BAM files with *SAMtools sort* version 1.9, and marked duplicate reads with *Picard MarkDuplicates* version 2.22.7 (http://broadinstitute.github.io/picard/), all done with the *MAP* function of the *PseudoreferencePipeline*. The analysis includes 43 high coverage red individuals and 42 high coverage green individuals. The five individuals isolated from red galls that did not carry the associated GWAS SNPs in *dgc* were excluded since the genetic basis for their gall coloration is unknown.

We subset the BAM file for each individual to contain only the target enrichment region on chromosome 1 from 40,092,625 to 40,892,915 bp using *SAMtools view* version 1.9 and generated merged BAM file for each color using *SAMtools merge*. The discordant reads were then extracted from each BAM file using *SAMtools view* with flag *-F 1286* and the percentage of discordant reads was calculated as the ratio of the number of discordant reads over the total number of mapped reads for each 5000 bp window.

To further explore the possibility that chromosomal aberrations near the GWAS signal might differ between red- and green-gall producing individuals, we plotted the mapping locations of discordant reads in the 100 kbp region near the 11 GWAS hits (40,450,000 - 40,550,000 bp) for red individuals, since the *H. cornu* reference was made from a green individual. We obtained the read ID for all the discordant reads within the 100kbp region and extracted all occurrences of these reads from the whole genome BAM file, regardless of their mapping location. We then extracted the paired-end reads from the discordant reads BAM file using *bedtools bamtofastq* version 2.29.2 and used *bwa mem* to map these reads as single-end reads for read 1 and read 2 separately to a merged reference containing the *H. cornu* genome and the 343 *Acyrthosiphon pisum* transposable elements annotated in *RepBase* (Jurka 1998). We then removed duplicates and sorted the BAM file using *SAMtools rmdup* and *sort* and determined the mapping location of all discordant reads using *SAMtools view*. We masked the windows from chromosome 1: 40,400,000 - 40,499,999 bp and 40,500,000 - 40,599,999 bp in the genome-wide scatter plot of discordant reads mapping because the majority of the discordant reads are expected to map to these regions and displaying their counts would obscure potential signals in the rest of the genome.

### Large scale survey of 11 *dgc* SNPs associated with gall color

Aphids were collected from red and green galls as described above for the GWAS study directly into Zymo DNA extraction buffer and ground with a plastic pestle. DNA was prepared using the ZR-96 Quick gDNA kit. We developed qPCR assays and amplicon-seq assays to genotype all individuals at all 11 SNPs (primers, etc below). PCR amplicon products were barcoded and samples were pooled for sequencing on an Illumina platform.

Adaptors were trimmed from amplicon reads using *Trim Galore!* version 0.6.5 and *cutadapt* version 2.7. The wrapper pipeline is available at github.com/YourePrettyGood/ParsingPipeline. We mapped reads to a 34 kbp region of chromosome 1 of the *H. cornu* genome that includes the amplicon SNPs (40,477,000 – 40, 511,000 bp) with *bwa mem* version 0.7.17, sorted BAM files with *SAMtools sort* version 1.9, and re-aligned indels using *GATK IndelRealigner* version 3.4. No marking of duplicates was done given the nature of amplicon sequencing data. To maximize genotyping efficiency and improve accuracy, we performed variant calling with two distinct pipelines: *SAMtools mpileup* version 1.7 (Li et al., 2009) plus *BCFtools call* version 1.7, and *GATK HaplotypeCaller* version 3.4. The mapping and indel re-alignment pipelines are available as part of the *MAP* (with flag *only_bwa*) and *IR* functions of the *PseudoreferencePipeline*. Using the same *PseudoreferencePipeline*, variant calling was performed using the *MPILEUP* function of *BCFtools* and *HC* function of *GATK*. FASTA sequences for each individual, where the genotyped SNPs were updated in the reference space, were then generated for both *BCFtools* and *GATK* variant calls using the above *PseudoreferencePipeline*’s *PSEUDOFASTA* function, with flags “*MPILEUP, no_markdup*” and “*HC, no_markdup*” respectively. The *BCFtools* SNP updating pipeline used *bcftools filter*, *query*, and *consensus* version 1.9, and we masked all sites where MQ <= 20 or QUAL <= 26 or DP <= 5. The HC SNP updating pipeline used *GATK SelectVariants* and *FastaAlternateReferenceMaker* version 3.4, and we masked all sites where MQ < 50, DP <= 5, GQ < 90 or RGQ < 90.

We then merged the variant calls from *BCFtools* and *GATK*, as well as the qPCR genotyping results, and manually identified all missing or discrepant genotypes. We manually curated these missing or discrepant genotype calls from the indel realigned BAM files using the following criteria: for heterozygous calls, the site has to have at least two reads supporting each allele, and for homozygous calls, the site has to have at least ten reads supporting the allele and no reads supporting alternative alleles.

### RNA-seq of salivary glands from aphids inducing red and green galls

We dissected salivary glands from fundatrices isolated from green and red galls in PBS, gently pipetted the salivary glands from the dissection tray in < 0.5uL volume of PBS, and deposited glands into 3 uL of Smart-seq2 lysis buffer (0.2%Triton-X 100, 0.1 U/uL RNasin® Ribonuclease Inhibitor). RNA-seq libraries were prepared with a single-cell RNA-seq method developed by the Janelia Quantitative Genomics core facility and described previously (Cembrowski et al. 2018).

RNAseq libraries were prepared as described above for red and green gall samples except that the entire 3uL sample of salivary glands in lysis buffer was provided as input. Barcoded samples were pooled and sequenced on an Illumina NextSeq 550. We detected 9.0 million reads per sample on average. We replaced the original oligonucleotides with the following oligonucleotides to generate unstranded reads from the entire transcript:

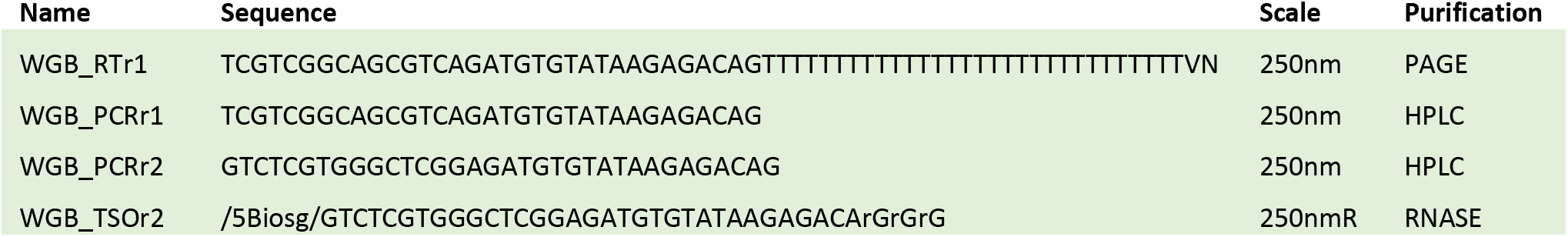

Samples were PCR amplified for 18 cycles and the library was prepared using ¼ of the standard Nextera XT sample size and 150 pg of cDNA.

### Differential expression analyses of fundatrix salivary glands from red and green galls

All differential expression analyses for plant and aphid samples were performed in *R* version 3.6.1 (R Core Team 2018) and all *R* Notebooks are provided on Github (will be posted prior to publication). Adapters were trimmed using *cutadapt* version 2.7 and read counts per transcript were calculated by mapping reads to the genome with *hisat2* version 2.1.0 (Kim, Langmead, and Salzberg 2015) and counting reads per gene with *htseq-count* version 0.12.4 (Anders, Pyl, and Huber 2015). In R, technical replicates were examined and pooled, since all replicates were very similar to each other. We performed exploratory data analysis using interactive multidimensional scaling plots, using the command *glMDSPlot* from the package *Glimma* (Su et al. 2017), and outlier samples were excluded from subsequent analyses. Differentially expressed genes were identified using the *glmQLFTest* and associated functions of the package *edgeR* (Robinson, McCarthy, and Smyth 2009). Volcano plots were generated using the *EnhancedVolcano* command from the package *EnhancedVolcano* version 1.4.0 (Blighe, Rana, and Lewis 2020). Mean-Difference (MD) plots and multidimensional scaling plots were generated using the *plotMD* and *plotMDS* functions, respectively, from the package *limma* version 3.42.0 (Ritchie et al. 2015).

### *Hamamelis virginiana* genome sequencing, assembly, and annotation

Leaves from a single tree of *Hamamelis virginiana* were sampled from the Janelia Research Campus forest as follows. Branches containing leaves that were less than 50% expanded were wrapped with aluminum foil and harvested after 40 hours. Leaves were cleared of obvious contamination, including aphids and other insects, and then plunged into liquid N_2_. Samples were stored at −80°C and sent to the Arizona Genomics Institute, University of Arizona on dry ice, which prepared HMW DNA from nuclei isolated from the frozen leaves. The Janelia Quantitative Genomics core facility generated a 10X chromium linked-read library from this DNA and sequenced the library on an Illumina NextSeq 550 to generate 608M linked reads.

The *H. virginiana* genome was assembled with the *supernova* commands *run* and *mkoutput* version 2.1.1, with options minsize=1000 and style=pseudohap (Weisenfeld et al. 2017). We used 332M reads in the assembly to achieve raw coverage of 56X as recommended by the supernova instruction manual. BUSCO completeness analysis (Simao et al. 2015) was performed using the *gVolante* Web interface (Nishimura, Hara, and Kuraku 2017) using the plants database. Genome assembly and BUSCO statistics are reported below.

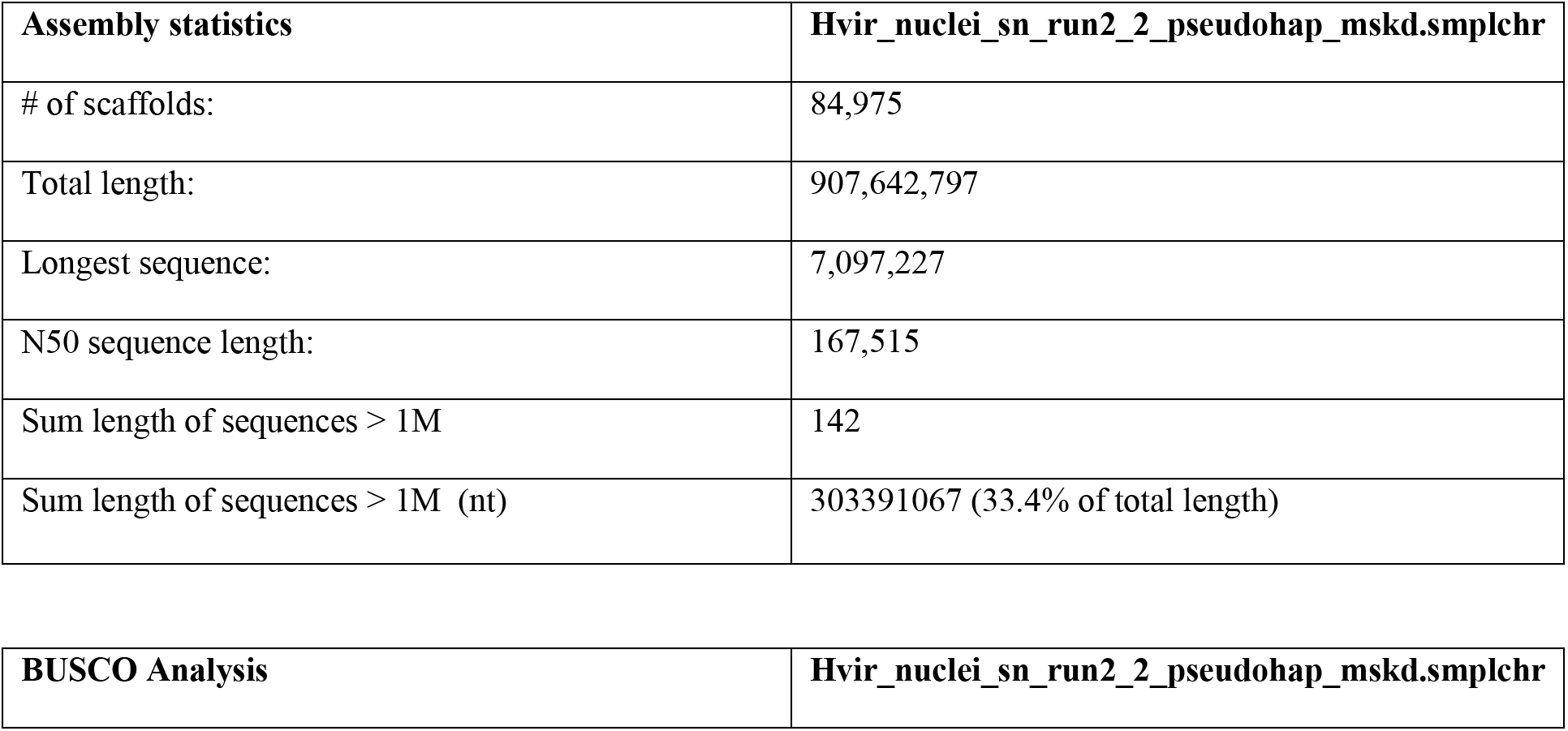
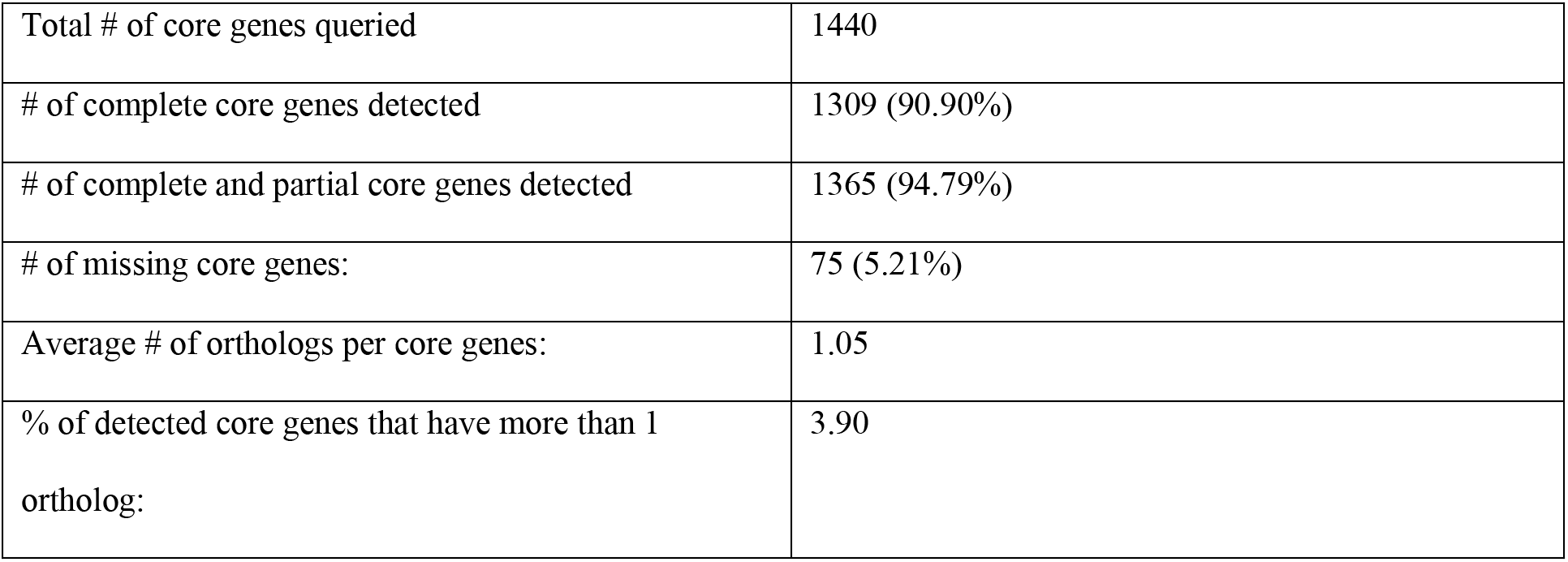

The assembled genome reference was repeat masked with soft masking using *RepeatMasker* version 4.0.9 (Smit n.d.). Twenty-five RNA-seq libraries from galls and leaves were used for genome annotation. RNA-seq reads were adapter trimmed using *cutadapt* version 2.7 and mapped to the genome using *HISAT2* version 2.1.0. Genome annotation was performed with *BRAKER* version 2.1.4 using the RNAseq data to provide intron hints (Altschul et al. 1990; Barnett et al. 2011; Camacho et al. 2009; Hoff et al. 2016, 2019; Li et al. 2009; Lomsadze et al. 2014; Ritchie et al. 2015; Stanke et al. 2006, 2008) and 3’ UTRs were predicted using *UTRme* (Radío et al. 2018).

### *H. virginiana* RNA extraction and RNA-seq library preparation

RNA was extracted from frozen *H. virginiana* leaf or gall tissue as follows. Plant tissue frozen at −80°C was placed into ZR BashingBead Lysis Tubes (pre-chilled in liquid N_2_) and pulverized to a fine powder in a Talboys High Throughput homogenizer (Troemer) with minimal thawing. Powdered plant tissue was suspended in extraction buffer (100 mM Tris-HCl, pH 7.5, 25 mM EDTA, 1.5 M NaCl, 2% (w/v) Hexadecyltrimethylammonium bromide, 10% Polyvinylpyrrolidone (w/v) and 0.3% (v/v) β-mercaptoethanol) and heated to 55°C for 8 min followed by centrifugation at 13,000 x g for 5 minutes at room temperature to remove insoluble debris (Jordon-Thaden et al. 2015). Total RNA was extracted from the supernatant using the Quick-RNA Plant Miniprep Kit (Zymo Research) with the inclusion of in-column DNAse I treatment. RNA-seq libraries were prepared with the Universal Plus mRNA-Seq kit (Nugen).

### Differential expression analysis of red versus green galls

We performed differential expression analysis on red and green galls by collecting paired red and green gall samples from the same leaves. In total, we collected 17 red galls and 23 green galls from 17 leaves. RNA was prepared as described above for plant material and RNA-seq libraries were prepared with a single-cell RNA-seq method modified for plant material described above.

These RNAseq libraries contained on average 4.4 million mapped reads per sample. Reads were quality trimmed and mapped to the transcriptome as described above. Only genes with greater than 1 count per million in at least 15 samples were included in subsequent analyses. Red and green samples clustered together in a principal components analysis (Fig. S9A, B) and no samples were identified as outliers. The expression analysis model included the effect of leaf blocking.

### Gall pigment extraction and analysis

Frozen gall tissue was ground to a powder under liquid nitrogen. We first tested for the presence of carotenoids by dehydrating gall tissue with methanol and extracting with hexane/acetone (Zhong et al. 2019). However, the lipophilic extract was colorless and all color remained in the polar phase and pellet, indicating that carotenoids do not contribute to red gall color.

We therefore next tested for presence of anthocyanins. Approximately 20 mg of ground gall tissue was suspended in 100 µL methanol (Optima grade, Fisher Scientific) and 400 µl of 5% aqueous formic acid (Optima grade, Fisher Scientific), vortexed for 30 seconds and centrifuged at 8000 x *g* for 2 min at 10° C. The supernatant containing pigment was filtered using a 0.2 µm, 13 mm diameter PTFE syringe filter to remove debris. Colorless pellet was discarded. Authentic anthocyanin pigment standards for malvidin 3,5-diglucoside chloride (Sigma Aldrich, St. Louis, MO, USA) and peonidin-3,5-diglucoside chloride (Carbosynth LLC, San Diego, CA, USA) were prepared at 1mg/mL in 5% aqueous formic acid.

Pigment separation and identification alongside standards was performed on a reverse phase C18 column (Acquity Plus BEH, 50 mm × 2.1 mm, 1.7 μm particle size, Waters, Milford, MA) using an Agilent 1290 UHPLC coupled to an Agilent 6545 quadrupole time-of-flight mass spectrometer (Agilent Technologies, Santa Clara, CA, USA) using an ESI probe in positive ion mode. Five µl of filtered pigment extract or a 1:100 dilution of anthocyanin standard was injected Solvent (A) consisted of 5% aqueous formic acid and (B) 1:99 water/acetonitrile acidified with 5% aqueous formic acid (*v/v*) (B). The gradient conditions were as follows: 1 min hold at 0% B, 4 min linear increase to 20% B, 5 min linear increase to 40% B, ramp up to 95% B in 0.1 min, hold at 95% B for 5 min, return to 0% B and hold for 0.9 min. The column flow rate was 0.3 mL/min, and the column temperature 30° C. The MS source parameters for initial anthocyanin detection were as follows: capillary = 4000 V, nozzle = 2000 V, gas temperature = 350° C, gas flow = 13 L/min, nebulizer = 30 psi, sheath gas temperature = 400° C, sheath gas flow = 12 L/min. DAD detection at 300 nm and 520 nm, and MS scanning from 50-1700 m/z at a rate of 2 spectra per second. Iterative fragmentation, followed by targeted MS/MS experiments, were performed using a collision energy = 35. Authentic standards confirmed the presence of peonidin-3,5-diglucoside and malvidin 3,5-diglucoside. The remaining anthocyanin species were identified using UV-Vis spectra, retention time relative to the other species in the sample, [M]^+^ precursor ions and aglycone fragment ions matching the respective entries in the RIKEN database (Sawada et al. 2012).

### Differential expression analysis of galls versus leaves

We performed RNA seq on 36 gall samples and 17 adjacent leaf samples. These gall samples did not overlap with the gall samples used in red versus green gall comparison described earlier. For larger galls, RNA was isolated separately from basal, medial, and apical gall regions (Table TAR Methods S8). Libraries were sequenced on an Illumina NextSeq 550 to generate 150 bp paired-end reads with an average of 8.1 million mapped reads per sample. Only genes expressed at greater than 1 count per million in at least 18 samples were included in subsequent analysis. Gall and leaf samples clustered separately in a principal components analysis (Fig. S9C, D) and no samples were excluded as outliers. We included only samples where there are paired gall and leaf samples from the same leaf and potential leaf effects were modeled in the different expression analysis.

To facilitate Gene Ontology (GO) analysis, the UniProt IDs of the differentially expressed genes were obtained by mapping the coding sequence of the *H. virginana* genome to the UniProt / Swiss-Prot database (Bateman 2019) using Protein-Protein BLAST 2.7.1 (Altschul et al. 1990; Camacho et al. 2009) and by extracting the differentially expressed genes (Fig. S9E). We used the WebGestalt 2019 webtool (Liao et al. 2019) to perform GO analysis on the differentially expressed genes.

### Differential expression analysis of *H. cornu* organs and life stages

RNAseq libraries were generated for fundatrix salivary glands (N = 20) and whole bodies (N = 8), G2 salivary glands (N = 6) and carcasses (N = 3), G5 salivary glands (N = 6) and carcasses (N = 2), and G7 salivary glands (N = 5). Libraries were generated as described above for salivary glands except that RNA samples of carcasses and whole bodies were prepared using the Arcturus PicoPure RNA Isolation Kit (Applied Biosystems). Only genes expressed with at least 1 count per million in at least 29 samples were included in subsequent analyses.

### Bioinformatic identification of *bicycle* genes in *H. cornu*

Genes that were upregulated specifically in the salivary glands of the fundatrix generation were prime candidates for inducing galls. We therefore identified genes that were upregulated both in salivary glands of fundatrices versus sexuals and in fundatrix salivary glands. These differentially expressed genes were then separated into genes with and without homologs containing some functional annotation. Homologs with previous functional annotations were identified using three methods: we performed (1) translated query-protein (*blastx*) and (2) protein-protein (*blastp*) based homology searches using BLAST 2.7.1 (Altschul et al. 1990; Camacho et al. 2009) against the UniProt / Swiss-Prot database (Bateman 2019) (Bateman 2019), and (3) Hidden-Markov based searches with the predicted proteins using *hmmscan* in *HMMER* version 3.1b2 (Eddy 2011) against the *pfam* database (Finn et al. 2014). For all predicted proteins, we also searched for secretion signal peptides using *SignalP-5.0* (Almagro Armenteros et al. 2019) and for transmembrane domains using *tmhmm* version 2.0 (Krogh et al. 2001). Gene Ontology analysis of genes with annotations that were enriched in fundatrix salivary glands was performed by searching for *Drosophila melanogaster* homologs of differentially expressed genes and using these *D. melanogaster* homologs as input into gene ontology analysis.

To determine whether any of these genes without detectable homologs in existing protein databases were homologous to each another, we performed sensitive homology searches of all-against-all of these genes using *jackhmmer* in *HMMER* version 3.1b2. We performed hierarchical clustering on the quantitative results of the *jackhmmer* analysis by first calculating distances amongst genes with the *dist* function using method *canberra* and clustering using the *hclust* function with method *ward.D2*, both from the library stats in *R* (R Core Team 2018). We aligned sequences of the clustered homologs using *MAFFT* version 7.419 with default parameters (Katoh 2002; Katoh and Standley 2013), trimmed aligned sequences using *trimAI* (Capella-Gutiérrez, Silla-Martínez, and Gabaldón 2009) with parameters -*gt* 0.50, and generated sequence logos by importing alignments using the functions *read.alignment* and *ggseqlogo* in the *R* packages *seqinr* (Charif and Lobry 2007) and *ggseqlogo* (Wagih 2017). After identification of the *bicycle* genes, we searched for additional *bicycle* genes in the entire *H. cornu* genome, which might not have been enriched in fundatrix salivary glands, using *jackhmmer* followed by hierarchical clustering to identify additional putative homologs. As described above, we manually annotated all these putative *bicycle* genes.

### Differential expression analysis of a single *H. cornu* fundatrix with a *dgc*^Green^ genotype inducing a red gall

Approximately 2.1% of fundatrices inducing red galls were homozygous for ancestral “green” alleles at all of the 11 *dgc* SNPs (Fig. 2E). Five such individuals were found in our original GWAS study and we did not observe any variants in the *dgc* gene region that were associated specifically with these individuals, suggesting that they carried variants elsewhere in the genome that caused them to generate red galls. Since isolating salivary glands from fundatrices is challenging and time consuming, we were unable to systematically examine transcriptome changes in the salivary glands of the rare individuals homozygous for *dgc*^Green^ that induced red galls. However, we fortuitously isolated salivary glands from one fundatrix from a red gall that was homozygous for *dgc*^Green^. We performed whole-transcriptome sequencing of the salivary glands from this one individual and compared expression levels of all genes in the *bicycle* gene paralog group to which *dgc* belongs. We also examined the *dgc* transcripts produced by this individual and found no exonic SNPs in the *dgc* transcripts produced by this individual, indicating that this individual probably expressed a functional *dgc* transcript.

### *H. cornu* and *H. hamamelidis* polymorphism and divergence measurements in GWAS region

To summarize the polymorphism and divergence patterns in *H. cornu* and *H. hamamelidis* in the target enrichment region, we generated SNP updated FASTA sequences for each individual by mapping the *H. cornu* reads to the *H. cornu* genome reference and the *H. hamamelidis* reads to a *H. hamamelidis* SNP-updated reference genome in *H. cornu* genome coordinate space. We used the same genotyping pipeline as described in the enrichment region GWAS, using the *PseudoreferencePipeline*, and masked sites were DP <= 10, MQ <= 20.0, or QUAL <= 29.5. High coverage individuals of *H. cornu* (n=90) and *H. hamamelidis* (N=92) were included in this analysis, including both of the color phenotypes. Polymorphism within each species and divergence (Dxy) between species were calculated using a custom script *calculateDiversity* (available at https://github.com/YourePrettyGood/DyakInversions/tree/master/tools). Windowed measurements of each of these statistics are generated with the custom script *nonOverlappingWindows.cpp*, using window size 3000bp (script available at github.com/YourePrettyGood/RandomScripts). Sites that are missing genotypes due to masking in 50% or more of the samples are not included in the windowed average and windows where 50% or more of the sites are missing are excluded from the plot in Fig. S13E.

### Whole-genome polymorphism and divergence measurements in *H. cornu* and *H. hamamelidis*

To examine genome-wide pattern of polymorphism and divergence, we used the same whole-genome sequencing data as in the genome-wide association study for gall color. This data set includes samples from fundatrices isolated from 43 green galls and 47 red galls for *H. cornu* and 48 galls for *H. hamamelidis*. Reads were mapped using *bwa mem* version 0.7.17-r1188. The *H. cornu* reads were mapped to the *H. cornu* reference genome and the *H. hamamelidis* reads were mapped to the *H. hamamelidis* SNPs updated reference genome, which are in the same coordinate space. We joint genotyped each species separately using *bcftools* version 1.9 *mpileup* and *call-m* to generate raw, multi-sample VCF for each species. Sites filtering for QUAL >= 20 was then done using *bcftools filter*, and insertions and deletions sites were removed using *VCFtools* version 0.1.16 *–remove-indels*. To generate a BED file of sites lacking genotypes for each sample, *bcftools view* with flag *-g^miss* was used to extract only the genotyped sites for each sample, *bcftools query* was used to generate the location of all SNPs and reference calls, and *bedtools* version 2.29.2 *complement* was used to generate the complementary positions to these genotyped calls as the positions to be masked. Consensus sequences were generated for each sample using *bcftools consensus*, with the flag *–iupac-codes*.

Polymorphism within each species and divergence (Dxy) between species were calculated using a custom script *calculateDiversity* (available at https://github.com/YourePrettyGood/DyakInversions/tree/master/tools). Windowed measurements of each of these statistics were generated with the custom script *nonOverlappingWindows.cpp*, using window size 1000bp (script available at github.com/YourePrettyGood/RandomScripts). Windows were designated as “*bicycle*” if they overlapped in coordinates with any *bicycle* genes; otherwise they were designated as “non-*bicycle*”. Sites masked in 80% or more of the samples were excluded in the windowed average and windows with 80% or more of the sites missing were excluded from genome-wide summary statistic calculations. This genotyping rate filter is more lenient to accommodate the higher amount of missing data due to the shallower sequencing depth.

### Bioinformatic analysis of *bicycle* gene structure and evolution

The *H. cornu* median exon size and number of exons per *bicycle* gene were calculated from the GFF annotation file Augustus.updated_w_annots.21Aug20.gff3. The *bicycle* genes alignment for the divergence tree was generated using *FastTree* version 2.1.11 (Price, Dehal, and Arkin 2010) and plotted using *ggtree* version 2.2.2 (Yu 2020) in R.

To calculate DnDs between *H. cornu* and *H. hamamelidis*, we first generated a masked *H. hamamelidis* reference genome by updating the *H. cornu* reference genome with *H. hamamelidis* SNPs. We randomly subset *H. hamamelidis* reads from 150bp PE 10X linked reads sequencing data to roughly 65X coverage using *seqtk* version 1.3 *sample -s100* (https://github.com/lh3/seqtk) and trimmed adapters using *Long Ranger* version 2.2.2 *basic* (10X Genomics). We then mapped the *H. hamamelidis* reads to the *H. cornu* reference and updated the SNPs using the *PseudoreferencePipeline*. Sites where MQ <= 20, QUAL <= 26 or DP <= 5 were masked, resulting in 21.0% of the genome being masked. This *H. hamamelidis* reference genome was used in the DnDs calculation.

DnDs between all orthologous genes in *H. cornu* and *H. hamamelidis* was calculated using the *codeml* function from the *PAML* package, version 4.9j (Yang 2007). The CDS sequences were extracted from both reference genomes for each gene using constructCDSesFromGFF3.pl and we split all degenerate bases into the two alleles to generate two pseudo-haplotypes using *fakeHaplotype.pl -s 42* (both scripts available at github.com/YourePrettyGood/RandomScripts). Haplotype 1 was used as input into *codeml* for both species. The settings used in *codeml* control file are: runmode=-2, seqtype=1, CodonFreq=0, clock=0, model=0, NSsites=0, fix_kappa=0, kappa=2, fix_omega=0, omega=1, Small_Diff=0.5e-6, method=0, fix_blength=0. Then, to evaluate whether DnDs is significantly different from 1 for each gene, we additionally ran a null model with the same control file, except with fix_omega=1. We then calculated the likelihood scores between the two models as 2×(lnL1-lnL2) and compared it to the 95^th^ percentile of the chi-square distribution with 1 degree of freedom. Mann-Whitney U test comparing DnDs of *bicycle* and non-*bicycle* genes was performed using the *wilcox.test* function in *R* version 3.6.1.

To perform McDonald-Kreitman tests for *bicycle* genes, we generated population sample of the *bicycle* coding regions for *H. cornu* by mapping RNA-seq data from 21 fundatrix salivary gland samples, which provided high coverage. We performed genotyping using the *STAR*, *IRRNA*, and *HC* functions in the *PseudoreferencePipeline*. This pipeline uses *STAR* version 2.7.3a for mapping and *IndelRealigner* and *HaplotypeCaller* in *GATK* version 3.4 for indel realignment and variant calling. SNP-updated FASTA sequences were generated for each individual from the genotype VCF using the *PSEUDOFASTA* function of the *PseudoreferencePipeline* without any additional masking. Then, for each individual, we split all degenerate bases into the two alleles to generate two pseudo-haplotypes using *fakeHaplotype.pl-s 42*. We calculated the number of synonymous and nonsynonymous polymorphisms (P_S_ and P_N_, respectively) and divergent substitutions (D_S_ and D_N_, respectively) using *Polymorphorama* version 6 (https://ib.berkeley.edu/labs/bachtrog/data/polyMORPHOrama/polyMORPHOrama.html) (Andolfatto 2007) for multiple categories of genes significantly over-expressed in the fundatrix salivary gland. The fraction of non-synonymous divergent substitutions that were fixed by positive selection (*α*; Fay, Wyckoff, and Wu 2002; Smith and Eyre-Walker 2002) was estimated using the Cochran-Mantel-Haenszel test framework (Shapiro et al. 2007). Using the *mantelhaen.test* function in *R* version 3.6.1, we estimated a common odds ratio 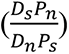 across a series of McDonald-Kreitman tables (McDonald and Kreitman 1991), and estimated 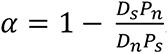. A MAF filter of 0.03 was applied to the polymorphism counts to reduce the downward bias in the estimator due to segregating weakly deleterious amino acid polymorphisms(Fay et al. 2002).

### Genome-wide tests of selection

To scan for genome-wide signatures of positive selection in *H. cornu* and *H. hamamelidis*, we used the joint-genotyped VCF for each species as described above. We filtered for QUAL >= 20 using *bcftools filter,* and 80% maximum missing genotype and biallelic sites using *VCFtools* version 0.1.16 *-max-missing 0.2*, *-max-alleles 2*, and *-min-alleles 2*. We then performed the selection scan using *SweeD-P* version 3.1 (Pavlidis et al. 2013), with one grid point per kilobase and *-folded* flag.

To determine the significance cutoff for the composite likelihood ratio (CLR) output by *SweeD*, we simulated 10 Mbp regions under neutrality for 100 haplotypes for each species using *MaCS* version 0.4f (Chen, Marjoram, and Wall 2008). The effective population size was set to 1,174,297 and 1,115,822 for *H. cornu* and *H. hamamelidis*, respectively. The mutation and recombination rates were set to 2.8e-9 and 1.1e-8, respectively, per bp per generation. The simulation output was converted to multi-sample VCF format using a custom script and run through *SweeD-P* using the same parameters as for the observed data. We ran ten iterations of the simulation for each species and combined the results to account for stochasticity of the simulations. The CLR cutoffs for the observed data were chosen as the 99^th^ quantiles of the CLR values from the neutral simulations of their respective species.

## Supplementary Text

### Chromosomal aberrations cannot explain association of 11 SNPs with gall color

We considered the possibility that DNA aberrations linked to the 11 associated SNPs might explain the red-green gall polymorphism. We found no evidence for large insertions, deletions, or other chromosomal aberrations in this region (Fig. S1E-I). In addition, we performed a genome-free association study (Rahman et al. 2018) and found strong linkage only for a subset of the 11 SNPs identified in the original GWAS (Fig. S3), further supporting the inference that these SNPs, and not other genetic changes, are associated with gall color.

### A small fraction of fundatrices likely carry variants not linked to *dgc* that induce red galls

Two percent of fundatrices from red galls were homozygous for the *dgc*^Green^ alleles at all 11 SNPs (Fig. 2E). These individuals do not carry exonic SNPs in *dgc* that would alter *dgc* function. We fortuitously performed RNA-seq on the salivary glands from one of these rare fundatrices and found that *dgc* was expressed at high levels (Fig. S7C), which suggests that the rare individuals that make red galls and are homozygous for *dgc*^Green^ alleles harbor variants at loci other than *dgc* that generate red galls.

### Gene ontology analysis of plant genes differentially expression in galls is consistent with the observed cell biology of gall development

Genes associated with cell division were strongly upregulated in galls (Fig. S9F), consistent with the extensive growth observed in gall tissue (Figure 1G-I). Genes associated with chloroplasts and photosynthesis were strongly downregulated in gall tissue (Fig. S9F), probably reflecting the reprogramming of plant leaf tissue from nutrient exporters to nutrient sinks (Inbar, Eshel, and Wool 1995). We also observed downregulation of genes related to immune response and multiple defense mechanisms in gall tissue (Fig. S9F), which may reflect suppression of the plant’s defense responses to infection. Some of these patterns are similar to patterns of differential expression observed in other gall systems (Hearn et al. 2019; Martinson, Werren, and Egan 2020). The pattern of differential expression does not appear to reflect a plant wound response, because wound response genes are not significantly over-expressed in gall tissue (Chi-square P-value = 0.367) and are biased toward under-represented.

### Evidence for roles of homologs of genes previously proposed as gall effector genes in *H. cornu*

Previously, the SSGP-71s effector gene family was described from the Hessian fly genome (Zhao et al. 2015) and it was suggested that these encode E3-ligase-mimicking effector proteins. We identified four homologs of the SSGP-71s effector gene family in *H. cornu* by searching *H. cornu* protein sequences using a hidden marker model search with a protein profile generated using the 426 SSGP-71 genes from Document S2, SSGP-71s tab, from Zhao et al. 2015 (hmmsearch with HMMER v3.2.1). However, none of these genes were differentially expressed in the fundatrix salivary gland. We also identified seven ubiquitin E3-ligase genes (Zhao et al. 2015) in *H. cornu*, two of which are upregulated in fundatrix salivary gland. However, none of these seven genes contain N-terminal secretion signals, making it unlikely that they could be injected into plants. Thus, these genes are unlikely to contribute to *H. cornu* gall development.

Ring domain proteins usually mediate E2 ubiquitin ligase activity, which can modify protein function or target proteins for proteosomal degradation (Metzger et al. 2014). The genome of a galling phylloxeran, a species from the sister family to aphids, has been shown to encode secreted RING domain proteins (Zhao, Rispe, and Nabity 2019), although it is not yet clear how the phylloxeran secreted RING domain proteins act and whether they are involved in gall-specific processes. We identified two *H. cornu* secreted RING domain proteins based on their similarity to proteins in the PFAM database (Finn et al. 2014), but only one was upregulated weakly in fundatrix salivary glands. Thus, secreted RING domain proteins are unlikely to contribute to *H. cornu* gall development.

### Characterization of *H. cornu* and *H. hamamelidis*

The sister species *H. cornu* and *H. hamamelidis* produce similar looking galls in adjacent geographic areas and were long confused as populations of a single species (von Dohlen and Stoetzel 1991). However, *H. cornu* exhibits a life cycle where aphids alternate between *Hamamelis virginiana* and *Betula nigra* (Fig. 4A), whereas *H. hamamelidis* does not host alternate to *B. nigra* and displays a truncated life cycle, where the offspring of the fundatrix develop as sexuparae (the equivalent of G6 in Fig. 4A) and deposit sexuals on *H. virginiana* in the autumn. Populations of *H. cornu* tend to live in the lowlands with prolonged summers and *H. hamamelidis* tend to live in the highlands and more northern latitudes, which experience shorted summers. In some locations, both species can be found making galls on the same trees.

Previous mtDNA sequencing supported the hypothesis that these are distinct species (von Dohlen, Kurosu, and Aoki 2002). Genome-wide divergence (Dxy) between these species was estimated to be 0.0189 and polymorphism was 0.0132 and 0.0125 for *H. cornu* and *H. hamamelidis*, respectively. Assuming a mutation rate *μ*=2.8e-9 (Keightley et al. 2014), we estimated effective population sizes using *θ* = 4*N_e_μ* for *H. cornu* and *H. hamamelidis* as 1,174,297 and 1,115,822, respectively.

## QUANTIFICATION AND STATISTICAL ANALYSIS

All quantitative methods and statistical analyses were explained in the **Method Details** section

