## Supplementary Materials for "A novel family of secreted insect proteins linked to plant gall development"

**This PDF file includes:**

Figs. S1 to S16
Tables S1 to S2

**Other Supplementary Materials for this manuscript include the following:**

Movie S1

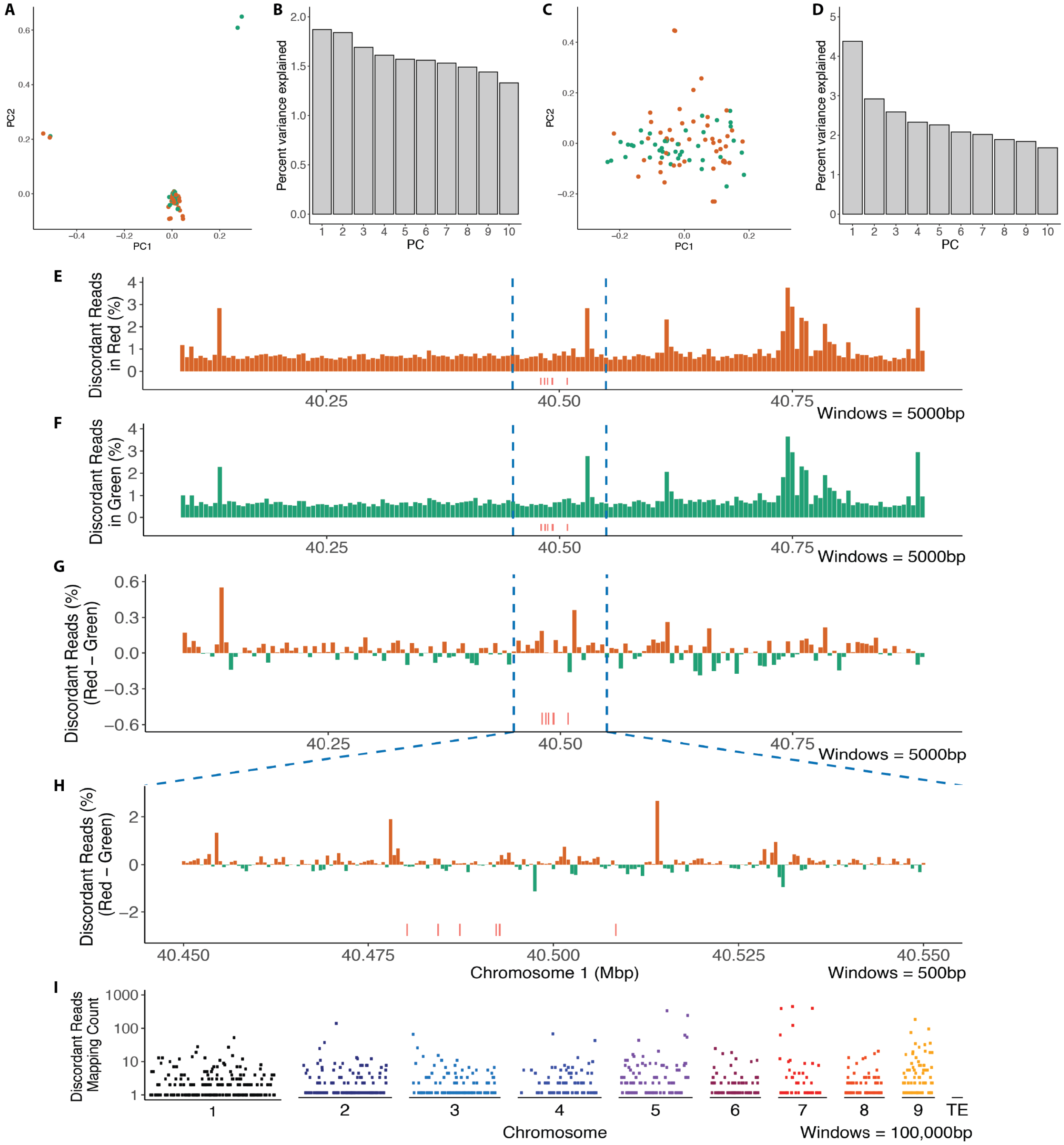

**Fig. S1. Supporting genetic evidence that the 11 SNPs located near *dgc* are the only genetic variants strongly associated with gall color.**

(**A-D**) The first two components of a principal components analysis (PCA) of genome-wide variation (**A**) and of variation in an 800 kbp window centered on *dgc* (**C**) reveal no large scale association of genetic variation with gall color. The top ten principal components of the genome-wide (**B**) and targeted (**D**) PCA each explain less than 2% and 5%, respectively, of the genetic variance.

(**E-I**) Distribution of mapped, unpaired Illumina sequencing reads provides no evidence for common large-scale genomic aberrations in aphids from either green (above) or red (below) galls. Discordant reads from fundatrices making red (**E**) and green (**F**) galls from the entire ~800kbp resequenced region are rare and are not specifically associated with gall color (**G**). Close examination of the genomic region harboring SNPs strongly associated with gall color (red vertical lines) shows that discordant reads are rare and not linked to the associated SNPs. Discordant reads do not map preferentially anywhere in the genome (**I**).

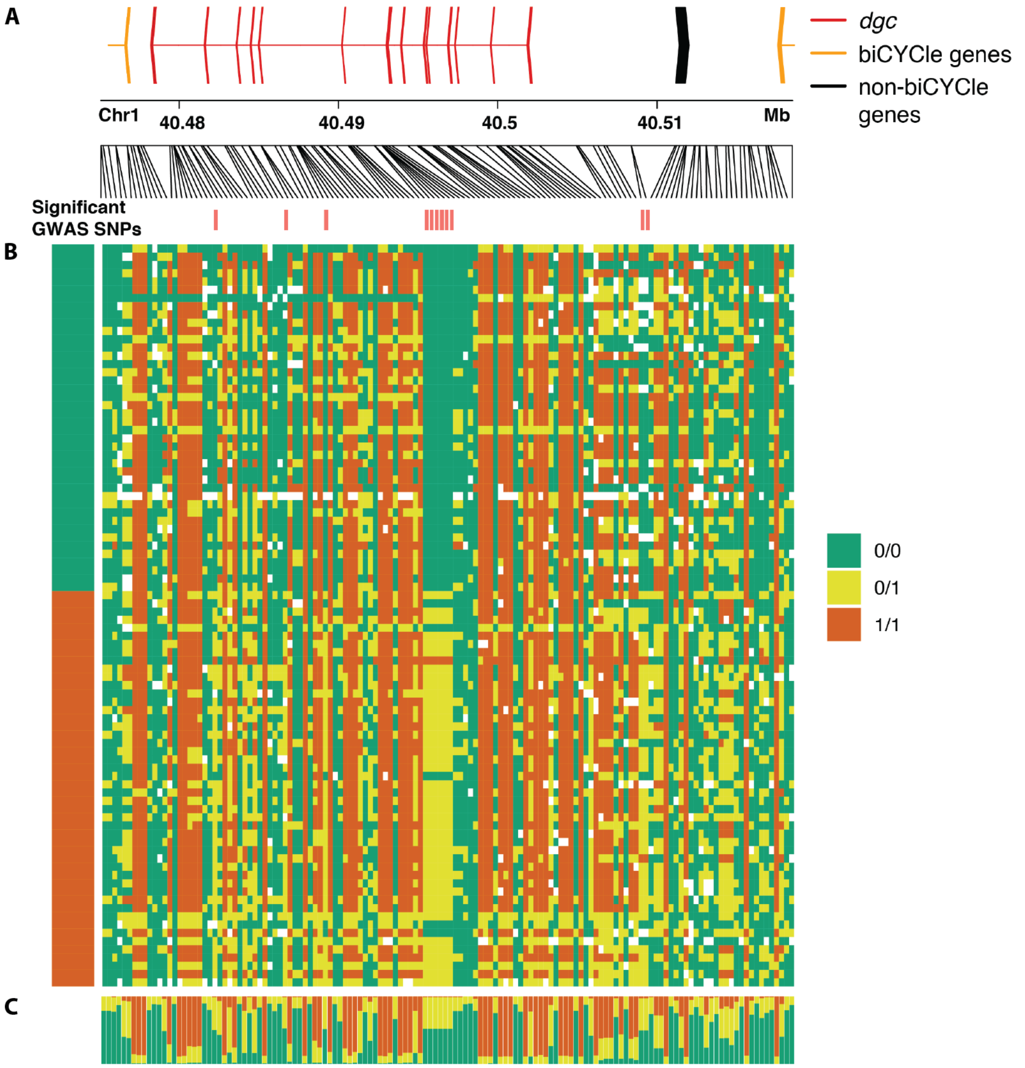

**Fig. S2. Genotypes of SNPs in the *dgc* gene region.**
(**A**) Gene models for the region ~40.475-40.52 of Chromosome 1. *Dgc* is labelled in red, other *bicycle* genes are labelled in yellow.
(**B**) Genotypes for each of the 90 individuals (42 from green galls, 48 from red galls; 4 low coverage individuals were excluded) from the original GWAS inferred from approximately 60X sequencing coverage for this region. Each row represents the genotypes at each of 698 SNPs. Genotypes are color coded as homozygous reference alleles (green), heterozygous (yellow), homozygous non-reference allele (orange). Data were thinned to exclude any SNPs within 500bp of each other.
(**C**) Frequencies of each genotype at each variable position.

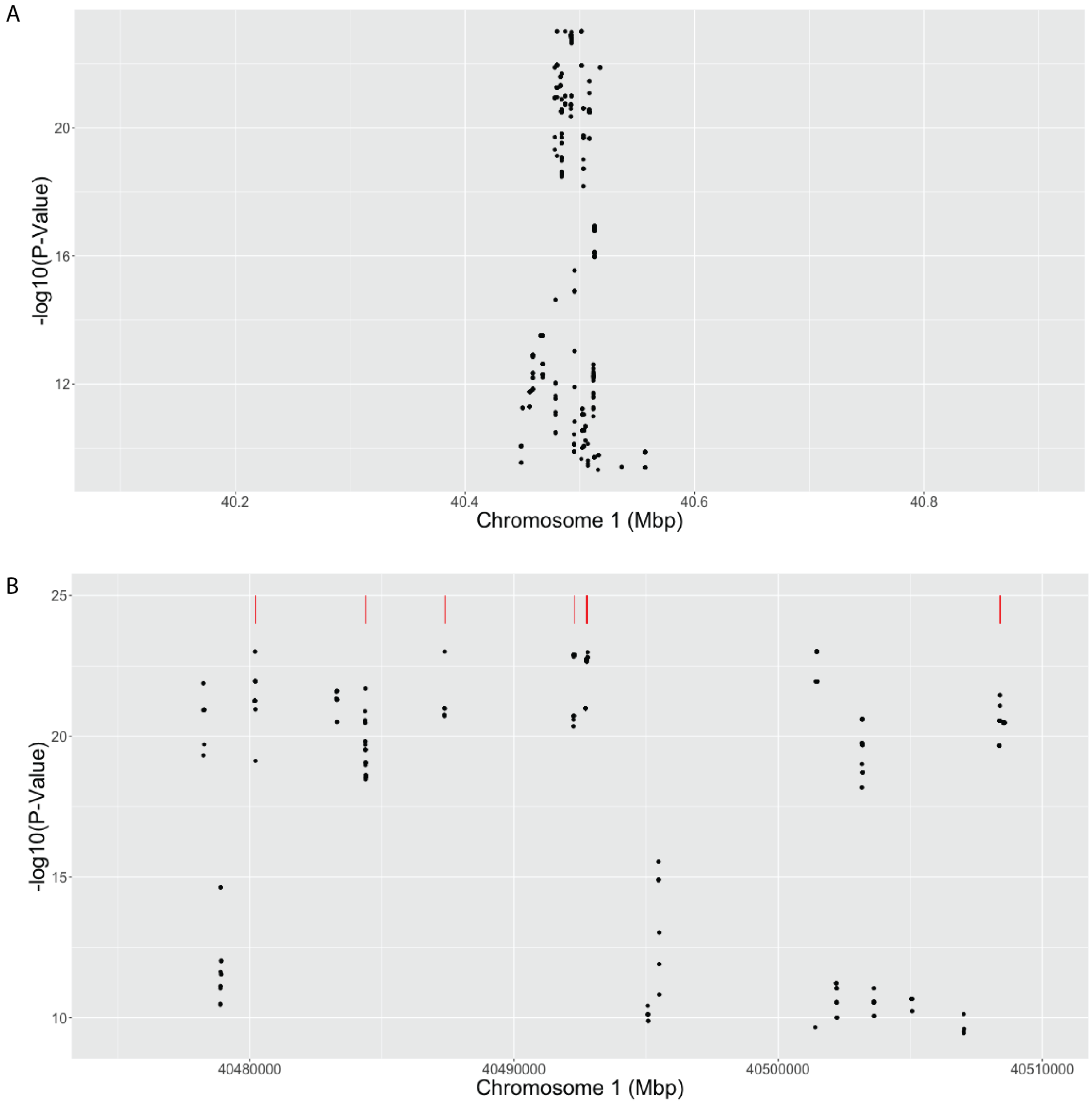

**Fig. S3. An alignment-free association test identifies most of the same associated SNPs as the GWAS.**

(**A**) Distribution of significant kmers associated with gall color detected by the alignment-free association test in the approximately 800 kb re-sequenced region centered on *dgc* (approximately 40.48-40.5 Mbp). The most significant kmers are found only located within and close to *dgc*. Non-significant kmers are not shown.

(**B**) Region from approximately 40.475-40.51 of plot (**A**), showing the finer-scale distribution of significant kmers. The red lines indicate SNPs detected in the original GWAS. All of the GWAS SNPs are confirmed by the alignment-free association test. The absence of significant kmers elsewhere in this region suggests that there are no genome rearrangements of transposon insertions that are associated with gall color.

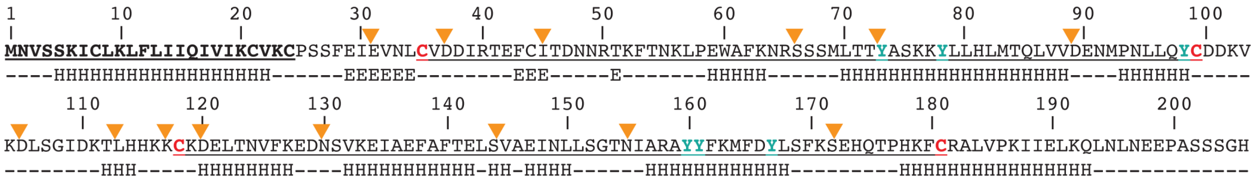

**Fig. S4. Predicted amino-acid sequence of g16073.**
Several secondary structure predictions are shown, including *SignalP* 5.0 prediction (Almagro Armenteros et al. 2019) of an N-terminal secretion signal from positions 1-24 (bold, underlined) and *jpred4* secondary structure prediction (Drozdetskiy et al. 2015) below primary sequence (H = helical, E = extended). Cysteines and tyrosines are colored red and blue, respectively, and the two CYC motifs are underlined. Orange triangles above protein sequence indicate intron-exon boundaries.

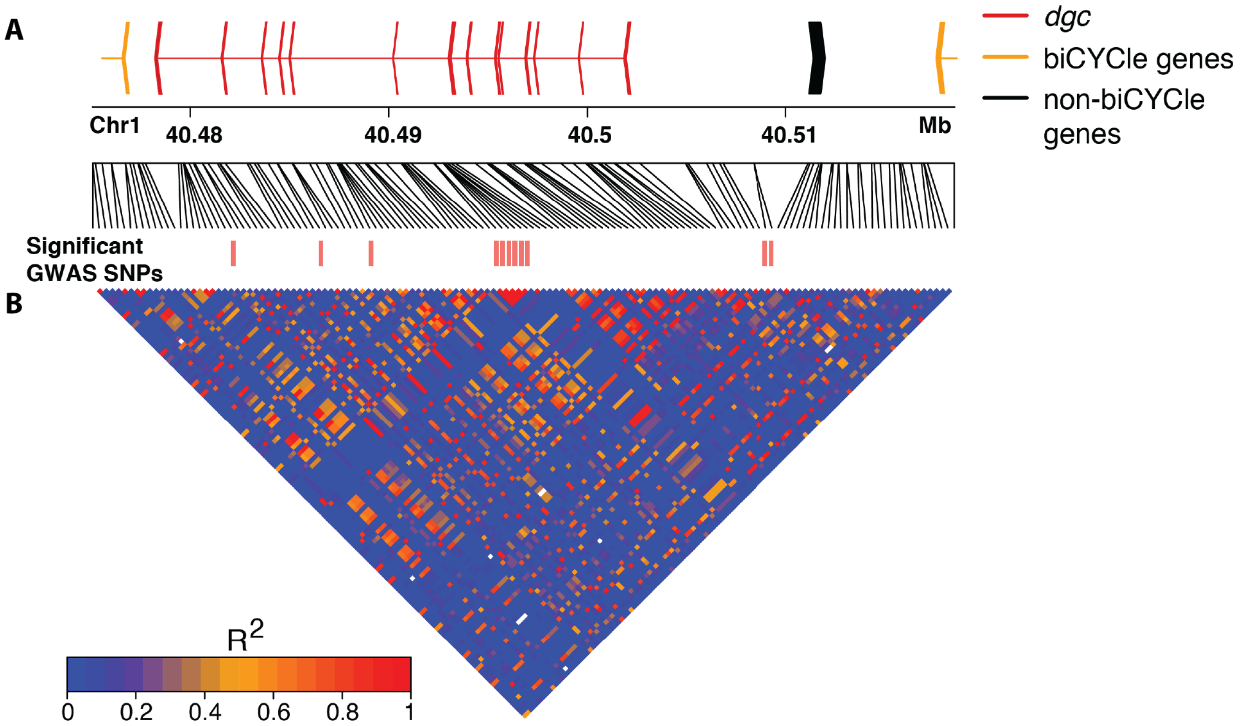

**Fig. S5. Linkage disequilibrium in *dgc* gene region.**

(**A**) Gene models for the region ~40.475-40.52 of Chromosome 1. *Dgc* is labelled in red, other *bicycle* genes are labelled in yellow.

(**B**) Linkage disequilibrium between all sites. Strong linkage disequilibrium is observed between the 11 SNPs linked to gall color (labelled as "Significant GWAS SNPs" above plot), but not between these SNPs and other variable sites.

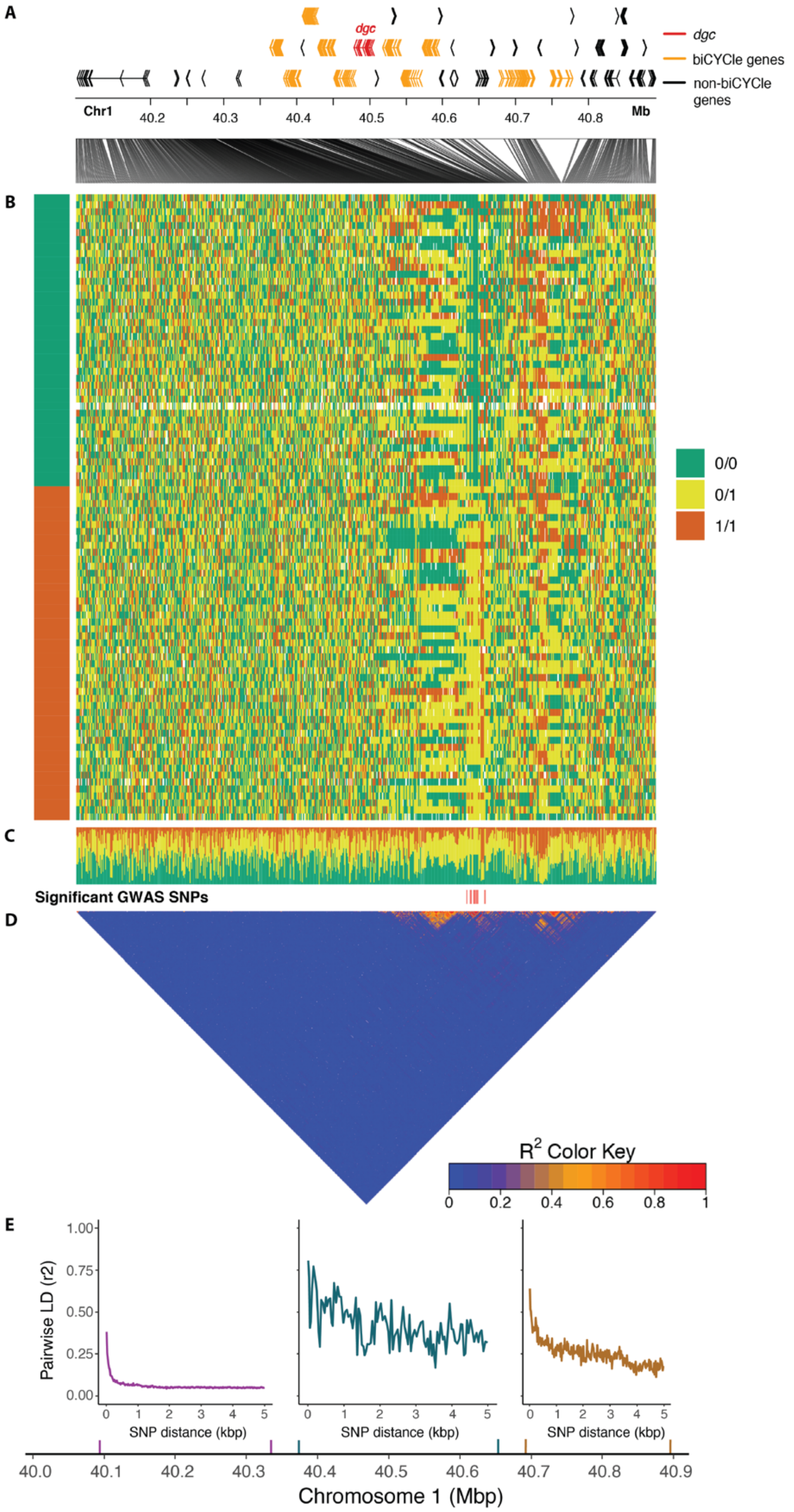

**Fig. S6. No evidence for long-distance linkage disequilibrium among variable sites in an 800 kbp region centered on *dgc*.**

(**A**) Gene models for the region ~40.1-40.9 of Chromosome 1. *Dgc* is labelled in red, other *bicycle* genes are labelled in yellow.
(**B**) Genotypes for each of the 90 individuals (42 from green galls, 48 from red galls; 4 low coverage individuals were excluded) from the original GWAS inferred from approximately 60X sequencing coverage for this region. Each row represents the genotypes at each of 698 SNPs. Genotypes are color coded as homozygous reference alleles (green), heterozygous (yellow), homozygous non-reference allele (orange). Data were thinned to exclude any SNPs within 500bp of each other.
(**C**) Frequencies of each genotype at each variable position.
(**D**) Linkage disequilibrium between all sites. Strong linkage disequilibrium is observed between the 11 SNPs linked to gall color (labelled as "Significant GWAS SNPs" above plot), but not between these SNPs and other variable sites.

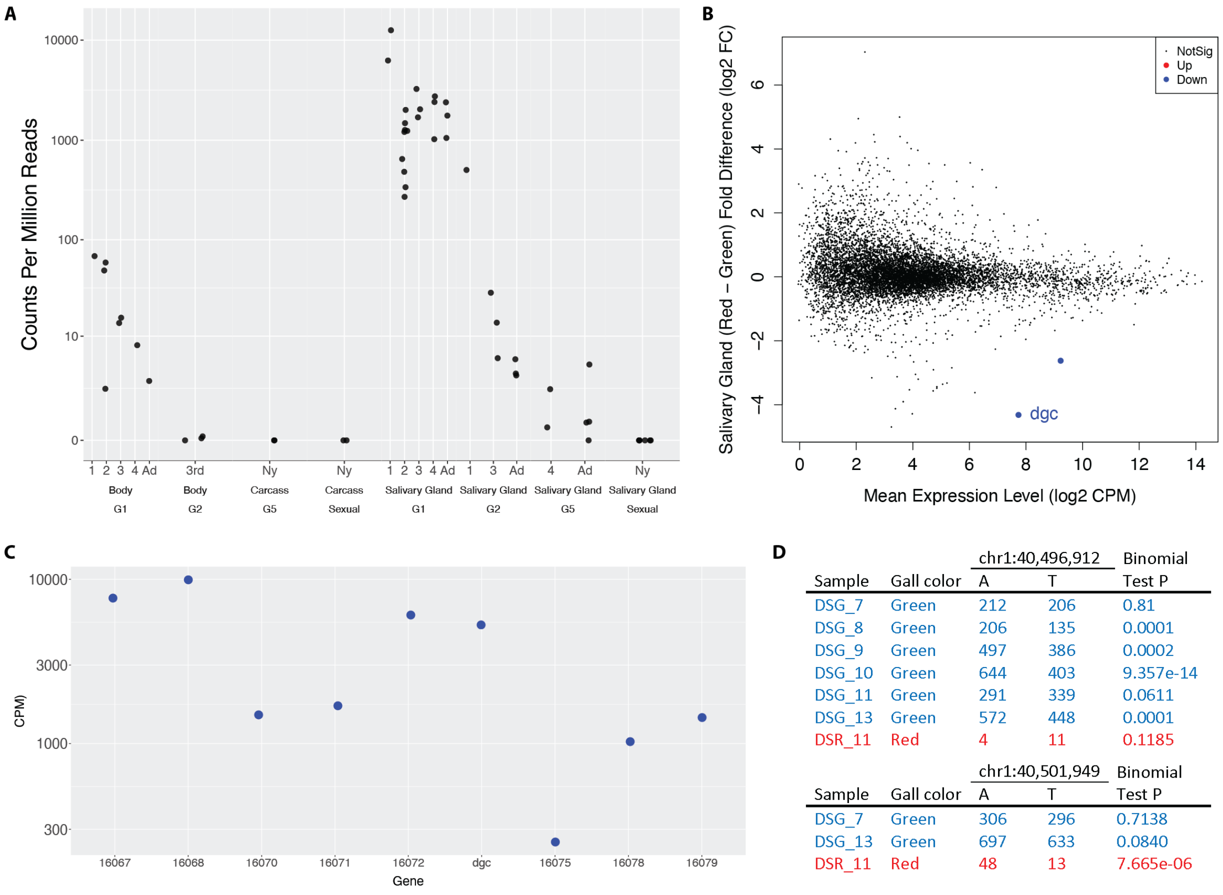

| | | chr1:40,496,912 | | Binomial |
|---|---|---|---|---|
| Sample | Gall color | A | T | Test P |
| DSG_7 | Green | 212 | 206 | 0.81 |
| DSG_8 | Green | 206 | 135 | 0.0001 |
| DSG_9 | Green | 497 | 386 | 0.0002 |
| DSG_10 | Green | 644 | 403 | 9.357e-14 |
| DSG_11 | Green | 291 | 339 | 0.0611 |
| DSG_13 | Green | 572 | 448 | 0.0001 |
| DSR_11 | Red | 4 | 11 | 0.1185 |

| | | chr1:40,501,949 | | Binomial |
|---|---|---|---|---|
| Sample | Gall color | A | T | Test P |
| DSG_7 | Green | 306 | 296 | 0.7138 |
| DSG_13 | Green | 697 | 633 | 0.0840 |
| DSR_11 | Red | 48 | 13 | 7.665e-06 |

**Fig. S7. Expression of *dgc***

(**A**) Expression of *dgc* in salivary glands, whole bodies, or carcasses (body minus salivary glands) throughout the *H. cornu* life cycle. Salivary glands were dissected from multiple nymphal stages and adults of four generations representing major morphs of the life cycle, G1 (fundatrix), G2, G5, and sexuals. *Dgc* is expressed at highest levels in salivary glands of fundatrices. *Dgc* expression declines in salivary glands through the instars of G2 animals and later generations and was not detected in salivary glands of sexuals. Expression observed in full bodies of G1 animals (fundatrices) probably reflect expression in the salivary glands and expression was not observed in carcasses.
(**B**) MD plot showing that *dgc* is the only gene differentially expressed in salivary glands of fundatrices sampled from red versus green galls and is strongly downregulated. The mean expression level of *dgc* illustrates that it is among the most highly expressed genes in salivary glands. Aphids from red and green galls were genotyped and carried *dgc*$^{\text{Red}}$/*dgc*$^{\text{Green}}$ and *dgc*$^{\text{Green}}$/*dgc*$^{\text{Green}}$ genotypes at all 11 SNPs associated with gall color, respectively.
(**C**) Expression levels of genes in the *bicycle* gene paralog group that includes *dgc* from a single individual with a *dgc*$^{\text{Green}}$/*dgc*$^{\text{Green}}$ genotype at all 11 SNPs that inhabited a red gall. *Dgc* is expressed at similar levels as *Horco_16072*, similar to the pattern observed for individuals that make green galls (Fig. 3B). Thus, it is likely that this individual, and possibly the remaining rare fundatrices that induce red galls with *dgc*$^{\text{Green}}$/*dgc*$^{\text{Green}}$ genotypes, do not do so through reduction

in *dgc* expression, but instead carry genetic variation at one or more other genes that induce red galls.

(**D**) Number of fundatrix salivary gland RNA-seq reads containing alternative *dgc* exonic SNPs at two genomic location for aphids collected from green (blue font) and red (red font) galls. Four out of eight green and one out of two red samples deviate from the expectation of equal expression from both alleles in an individual. Thus, there is no evidence for allelic imbalance between fundatrices making green and red galls. In addition, both alternative alleles in red samples are expressed at substantially lower levels than the same alleles in green samples. Thus, a single copy of the SNPs that repress *dgc* expression (Fig. 3B) appear to repress *dgc* from both alleles in heterozygous individuals.

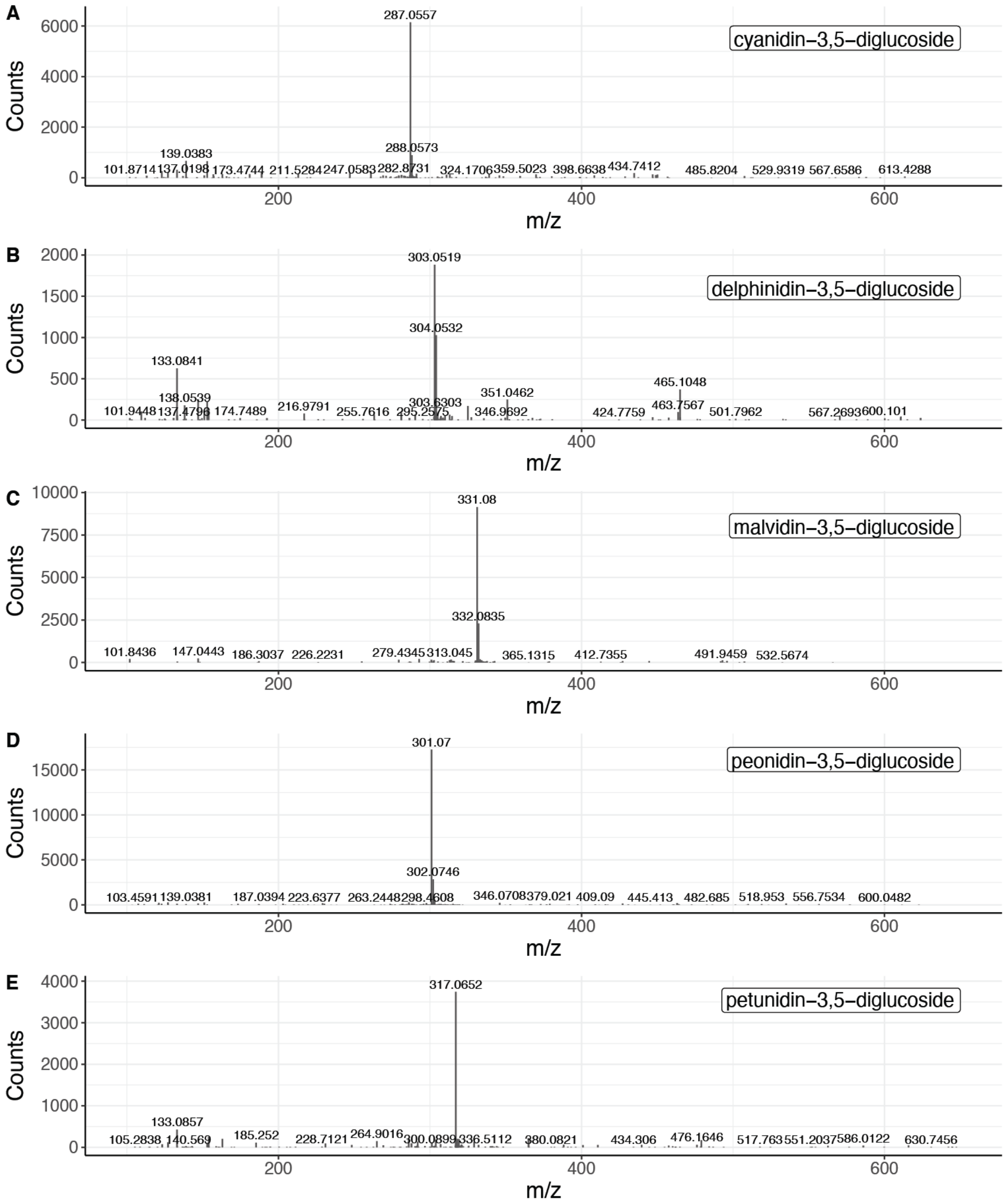

**Fig. S8. MS/MS spectra of anthocyanins detected in the red gall extract, which clearly demonstrate the aglycone product ions previously reported for cyanidin-3,5-diglucoside (A), delphinidin-3,5-diglucoside (B), malvidin-3,5-diglucoside (C), peonidin-3,5-diglucoside (D), and petunidin-3,5-diglucoside (E).**

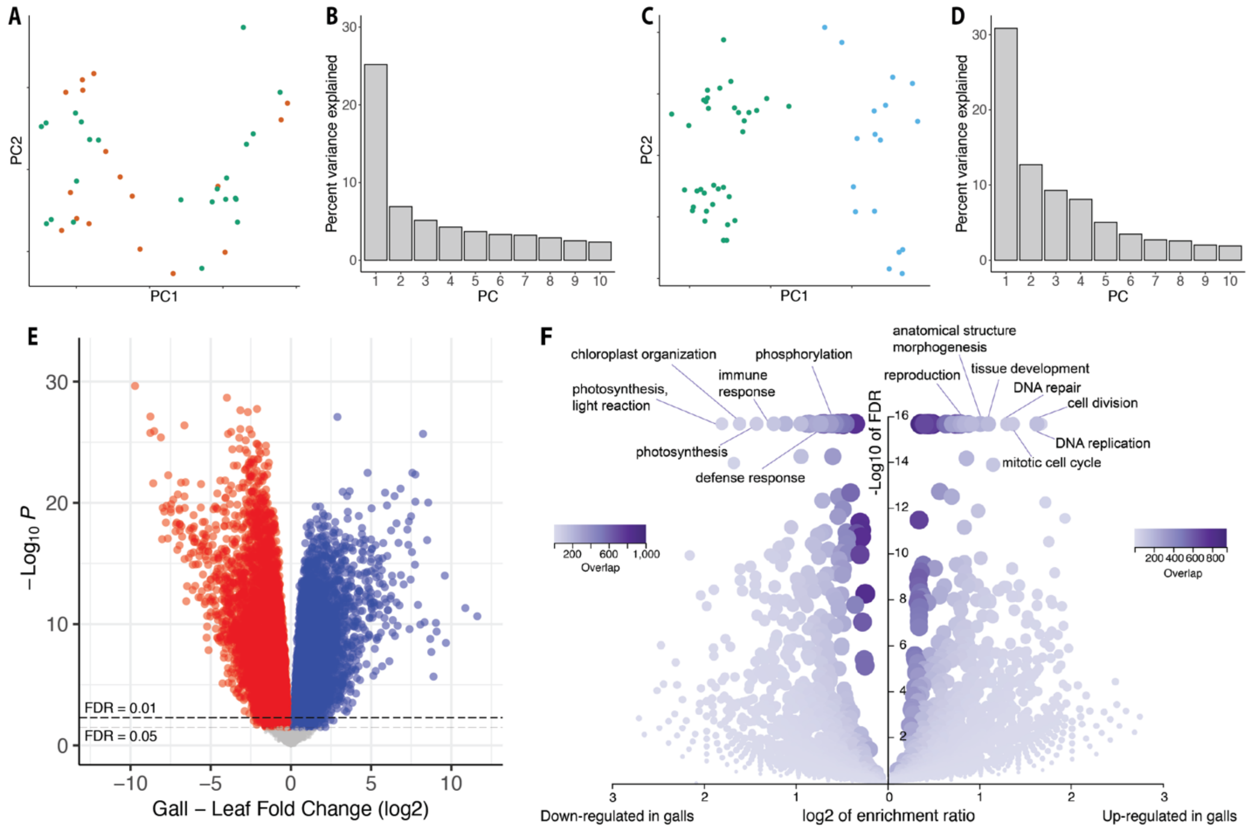

**Fig. S9. Genome-wide differential expression analysis of *H. virginiana* galls versus leaves**
(**A-D)** Principal components analysis (**A, C**) and percent variance explained of first 10 principal components (**B, D**) for the RNA-seq data of red (red dots) and green (green dots) galls (**A**, **B**) and galls (green dots) and leaves (blue dots) (**C, D**).
(**E**) Genome-wide differential expression analysis of *H. virginiana* transcripts isolated from galls (N = 36) versus leaves (N = 17). Approximately 60% of expressed genes are differentially expressed between gall and leaf tissue at FDR < 0.05.
(**F**) Gene ontology analysis of GO terms down (left) and up-regulated (right) in galls, presented as volcano plots. Genes involved in cell division and morphogenesis were strongly upregulated in galls and genes involved in photosynthesis were strongly down-regulated in galls.

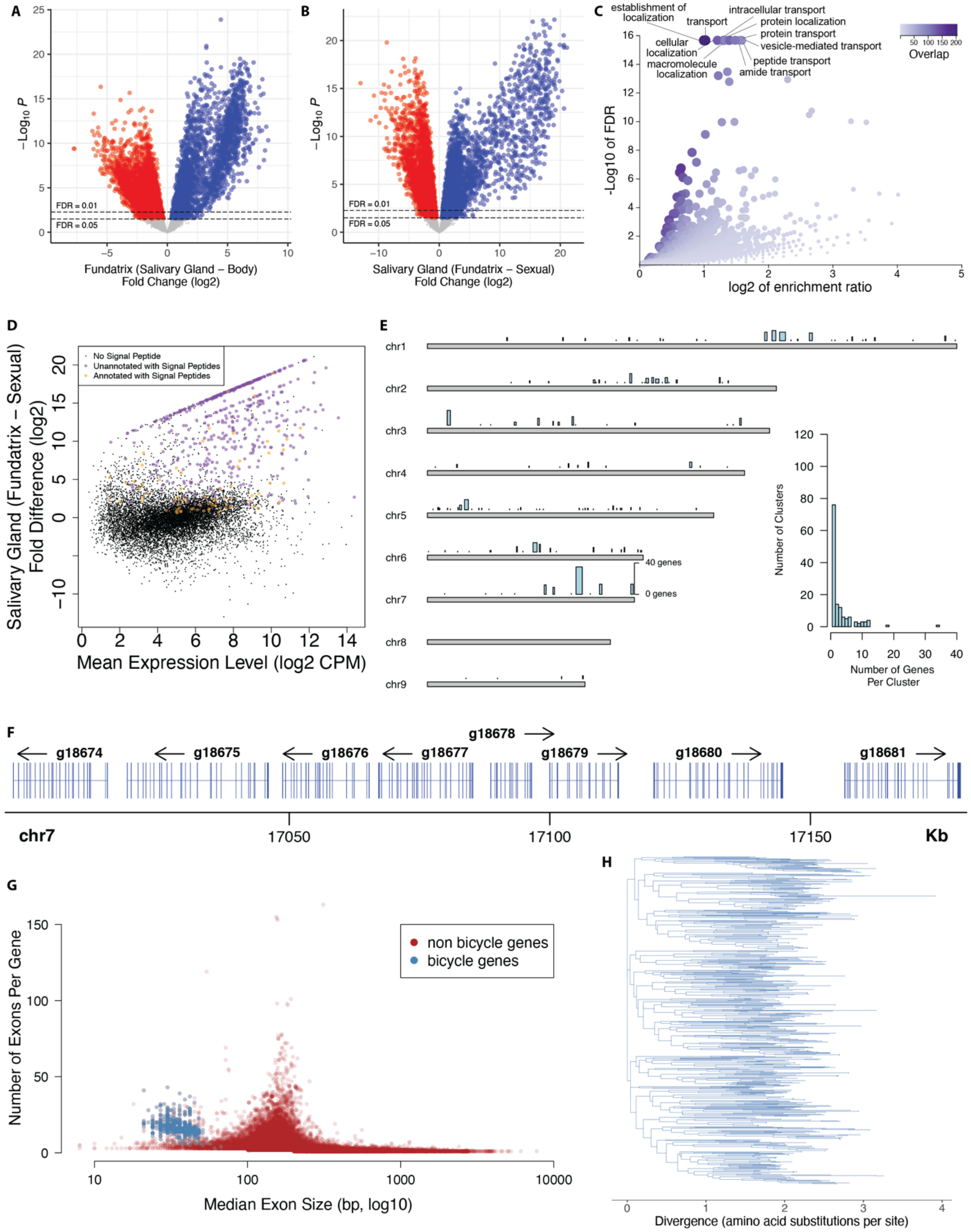

**Fig. S10. Further analysis of *H. cornu* fundatrix salivary gland enriched and *bicycle* genes.**

(**A, B**) Differential expression comparing gene expression in fundatrix salivary glands versus fundatrix whole bodies (**A**) and salivary glands of fundatrices versus sexuals (**B**) shown as volcano plots. Both comparisons reveal that a large number of genes are strongly over-expressed in fundatrix salivary glands. Intersection of the genes over-expressed in these two comparisons yielded the focal collection of genes over-expressed specifically in fundatrix salivary glands (Fig. 4B). Genes differentially over- or under-expressed at FDR < 0.05 are shown in blue or red, respectively.
(**C**) Volcano plot of Gene Ontology over-representation analysis of "Annotated" genes from Fig. 6B showing KEGG pathway terms for biological processes that were over-represented with a raw P value < 0.05. No GO terms were significantly under-represented with a raw P value < 0.05. Analyses performed with WebGestalt (Liao et al. 2019).
(**D**) MD plot of differential expression results shown in panel (B) with "Unannotated" and "Annotated" genes with signal peptides over-expressed in fundatrix salivary glands labeled in purple and yellow, respectively.
(**E**) Distribution of singleton *bicycle* genes and paralog clusters in the *H. cornu* genome. Number of genes per cluster and genomic range is indicating by height and width, respectively, of blue bars above chromosomes. Histogram of number of *bicycle* genes per paralog cluster is shown in inset.
(**F**) Example of part of a paralog cluster of *bicycle* genes from chromosome 7 of the *H. cornu* genome, illustrating abundance of small exons in each gene.
(**G**) Number of exons per gene versus median exon size for *H. cornu bicycle* (blue) and non-*bicycle* (red) genes. *Bicycle* genes possess an unusually large number of unusually small exons.
(**H**) Maximum likelihood phylogenetic tree of *H. cornu bicycle* gene amino acid sequences reveals extensive sequence divergence of *bicycle* genes.

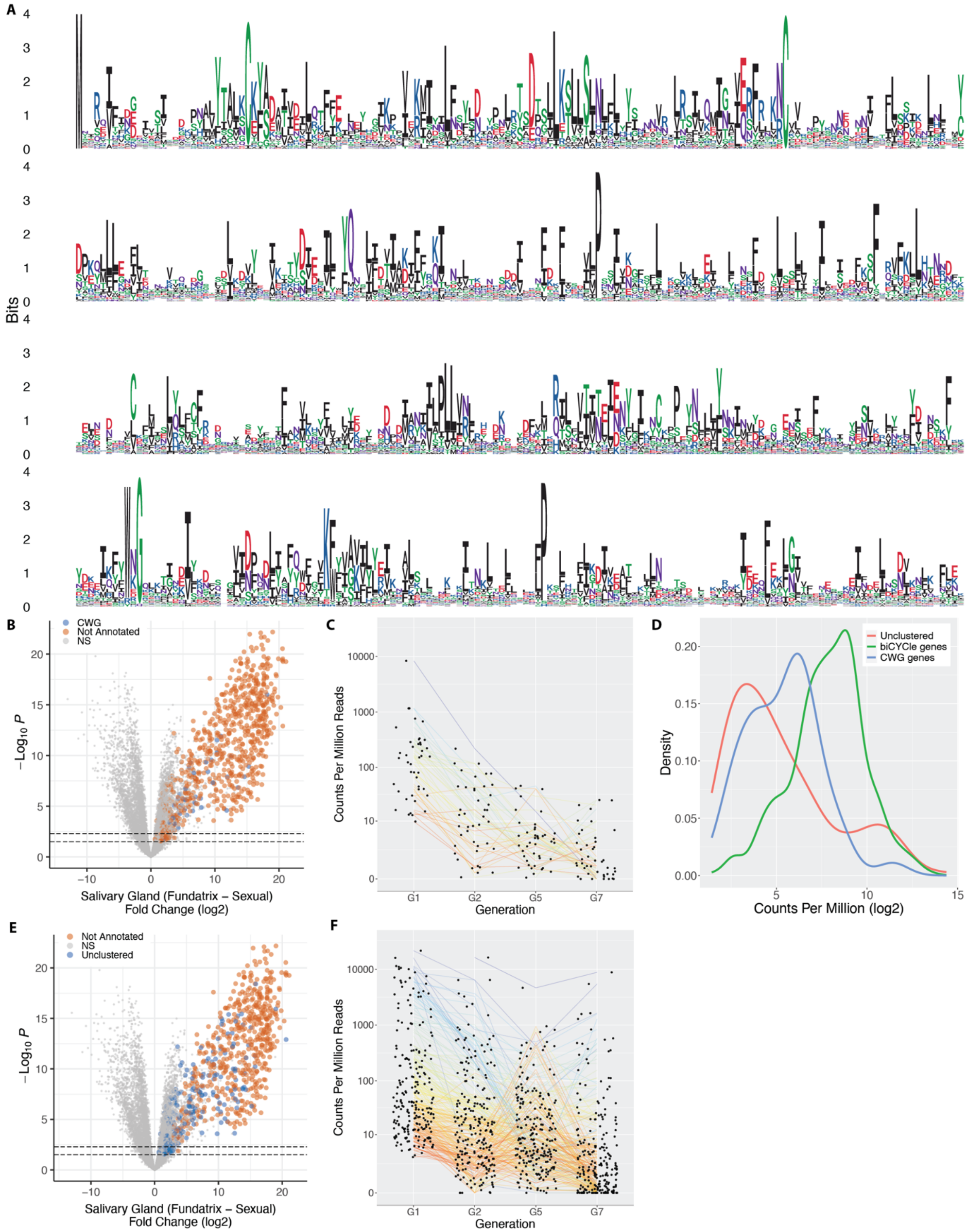

**Fig. S11. Further analysis of unannotated fundatrix salivary gland enriched genes.**
(**A**) Logo plot of the aligned predicted protein sequences from CWG genes. Sequence alignment used for logo was filtered to highlight conserved positions.

(**B, E**) Volcano plots of differential expression of fundatrix versus sexual salivary glands with *CWG* genes (**B**) and Unclustered genes (**E**) labelled in purple for clusters identified through hierarchical clustering (Fig. 6C).
(**C, F**) Expression levels of *CWG* (**C**) and Unclustered (**F**) genes in the salivary glands of individuals across four generations of the *H. cornu* life cycle. Lines connect the same gene across generations and are color coded by relative expression levels in G1, with blue to red representing most to least strongly expressed, respectively.
(**D**) *bicycle* genes are, on average, the most strongly differentially expressed category of unannotated genes.

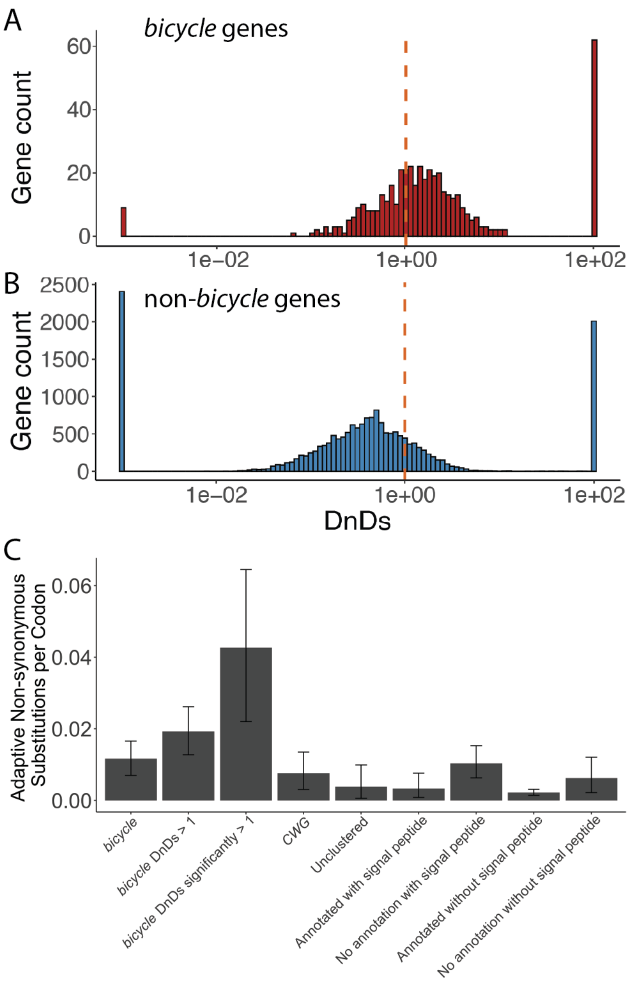

**Figure S12. Comparison of rate of non-synonymous to synonymous substitutions in *bicycle* versus non-*bicycle* genes**

(**A**) The majority of *bicycle* genes display $d_N/d_S$ values great than 1, with few showing strong sequence conservation ($d_N/d_S$ <<1). Dashed vertical red line indicates $d_N/d_S = 1$.

(**B**) Non-*bicycle* genes are more conserved, on average, than *bicycle* genes (Mann-Whitney U test p=2.6e-76. Dashed vertical red line indicates $d_N/d_S = 1$.

(**C**) Mean number of adaptive non-synonymous substitutions scaled by protein length for different categories of genes over-expressed in fundatrix salivary glands. As a proportion of protein length, *bicycle* genes display the fastest rate of adaptive evolution of any category of these genes. Error bars represent 95% confidence intervals. Note that the four categories on the right include all genes shown on the left, but categorized by whether genes were annotated and included a signal peptide. Thus, for example, the category "No annotation with signal peptide" is composed mostly of *bicycle* and *CWG* genes.

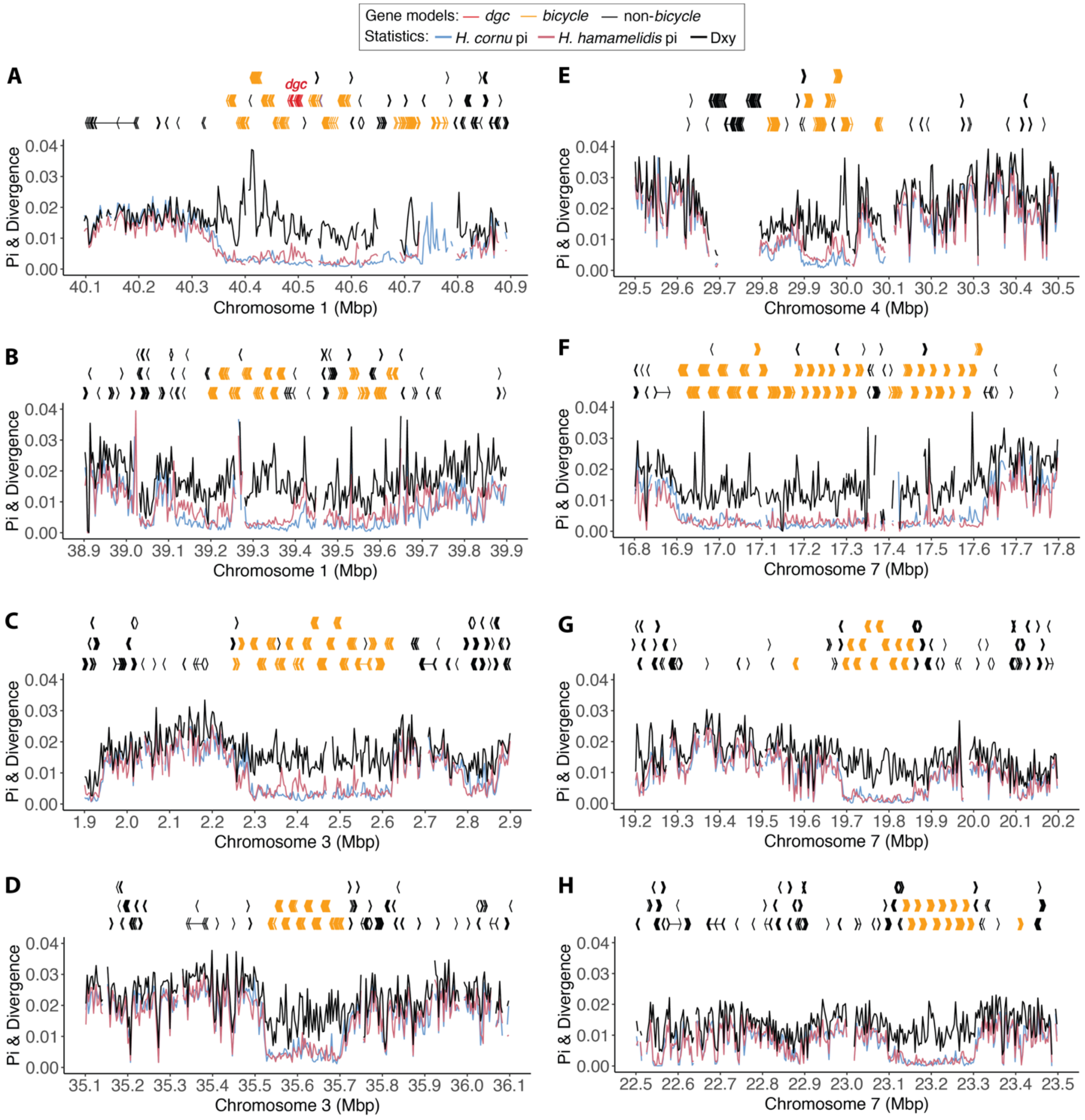

**Fig. S13. Polymorphism is depressed relative to divergence in *bicycle* gene regions.**
(**A**) Gene models in 800 kb region including *bicycle* gene cluster containing *dgc* shown above. Divergence between (black line) and polymorphism within *H. cornu* (blue line) and *H. hamamelidis* (pink line) in 3000bp windows shown below.
(**B-H**) Gene models and population genomic statistics for seven additional genomic regions containing *bicycle* gene clusters.

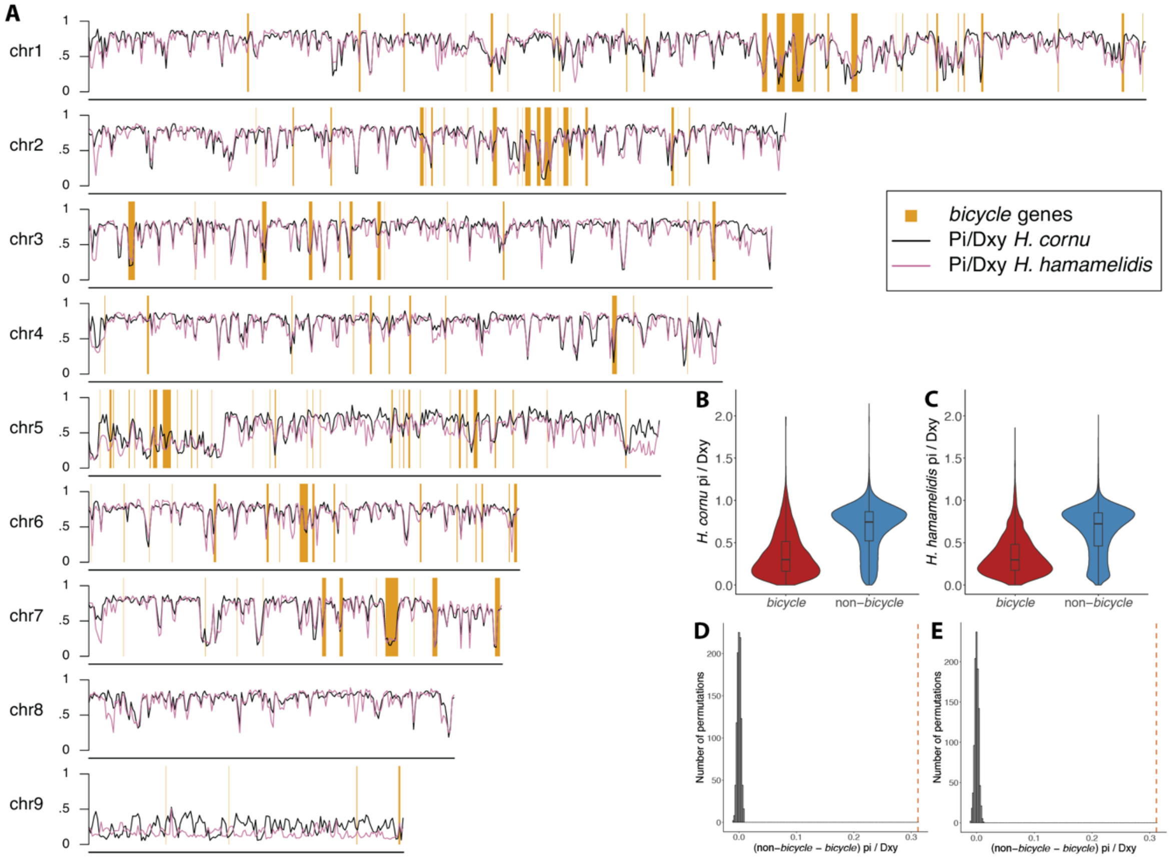

**Fig. S14. Strongly reduced polymorphism to divergence levels are found in *bicycle* gene regions**
(**A**) Genome-wide non-overlapping window moving averages of polymorphism (Pi) divided by divergence (Dxy) between *H. cornu* (black lines) and *H. hamamelidis* (purple lines). Gene models in the 800 kbp region centered on *dgc*. Bicycle genes are located within ranges indicated in orange.
(**B** and **C**) Ratio of Pi to Dxy for *bicycle* and non-*bicycle* gene regions in *H. cornu* (**B**) and *H. hamamelidis* (**C**).
(**D** and **E**) The observed difference in Pi/Dxy between non-*bicycle* and *bicycle* genes (dashed red line) is much larger than the expectation generated by permuting the locations of Pi/Dxy values relative to gene locations for both *H. cornu* (**D**) and *H. hamamelidis* (**E**).

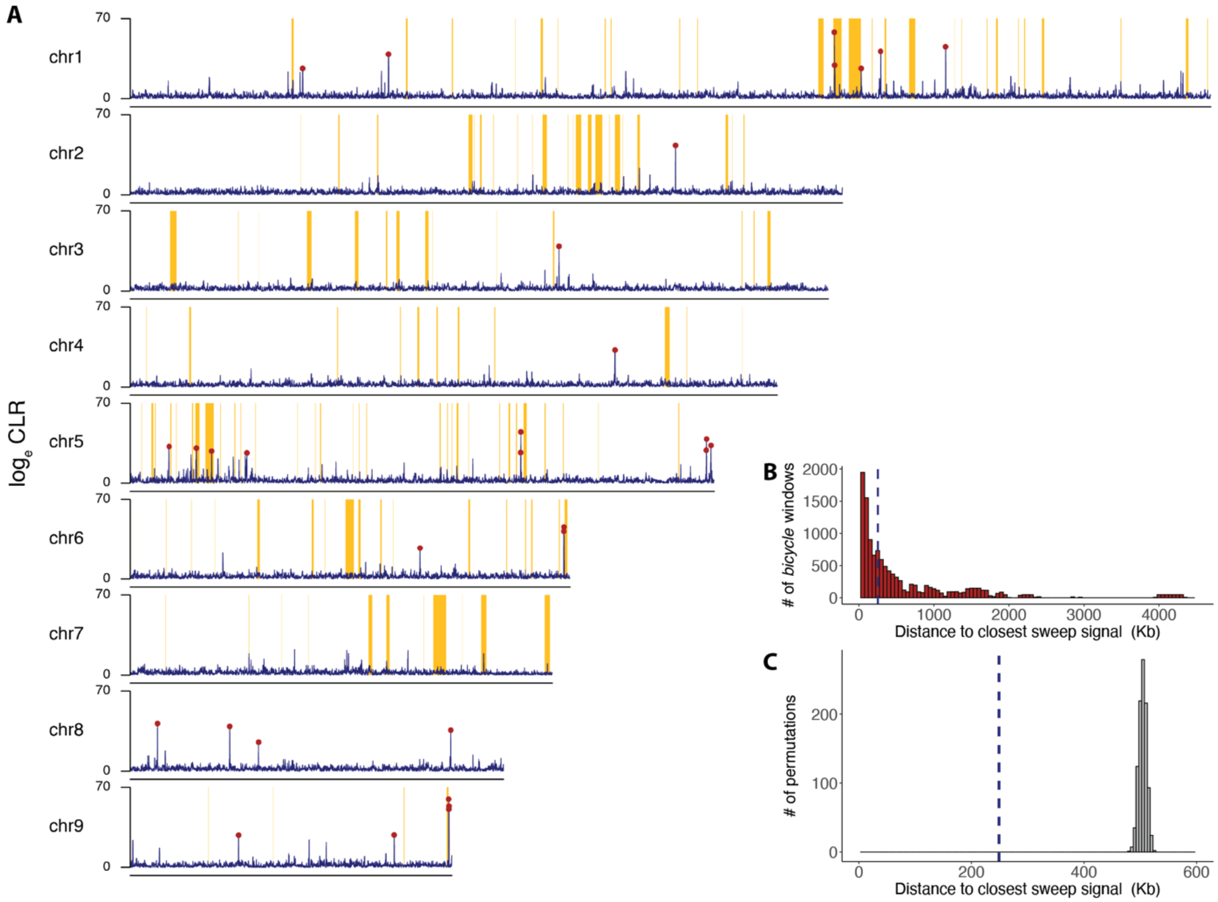

**Fig. S15. Genome-wide distribution of signatures of recent selective sweeps detected by SweeD in *H. cornu*.**
(**A**) Genome-wide distribution of the log$_e$ of the composite likelihood ratio (CLR) for evidence of a recent selective sweep is shown in purple. The location of significant signals ($P < 0.01$) calculated after simulations assuming neutrality (Supplementary text) is indicated with red circles. The locations of *bicycle* genes are indicated with orange rectangles.
(**B**) Distance from each *bicycle* gene to the closest significant selective sweep signal is shown as red histogram and dashed blue line indicates the median of this distribution.
(**C**) The median distance from each *bicycle* gene to the closest significant selective sweep signal (from **B**) is shown with dashed blue line and the values after 1000 permutations of sweep signals relative to gene locations are shown as grey histogram. The observed sweep signals are closer to *bicycle* genes than expected by chance.

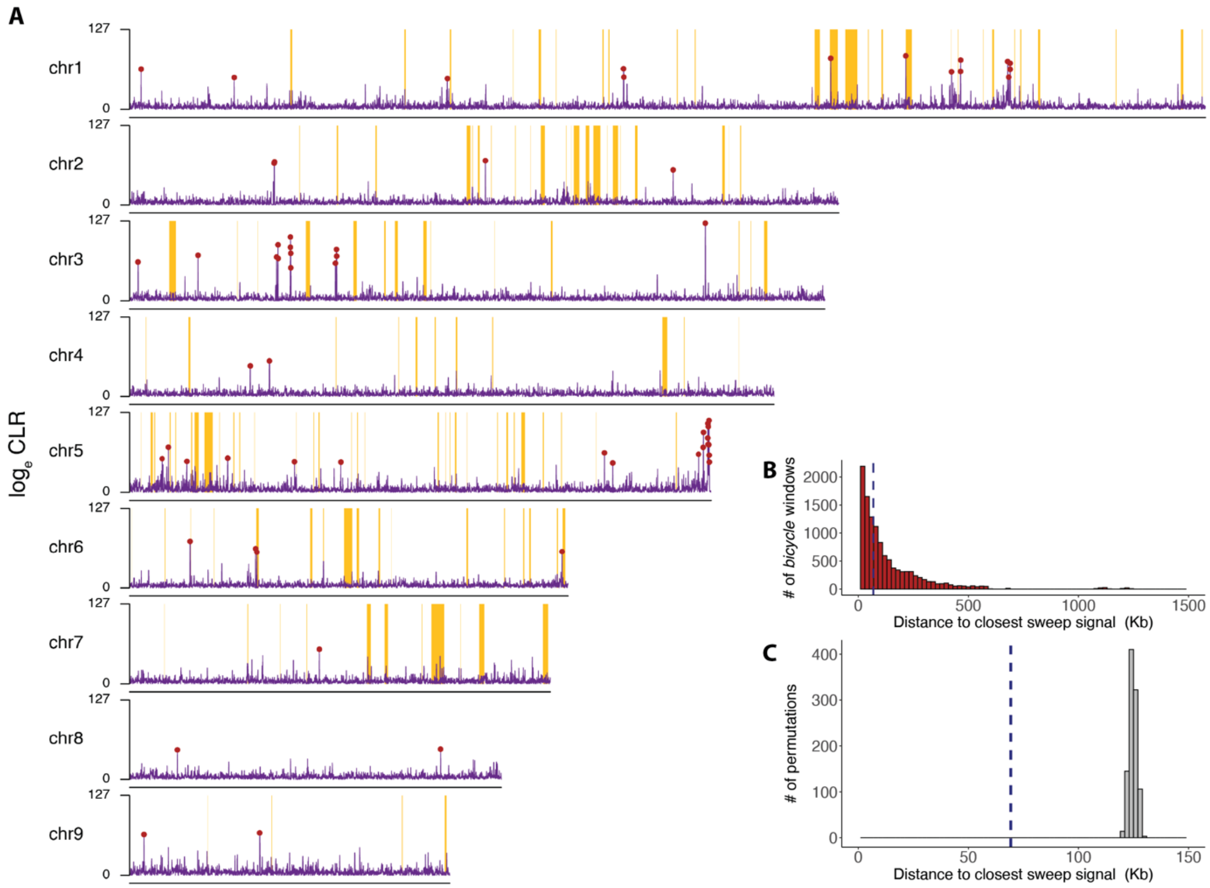

**Fig. S16. Genome-wide distribution of signatures of recent selective sweeps detected by SweeD in *H. hamamelidis*.**

(**A**) Genome-wide distribution of the $\log_e$ of the composite likelihood ratio (CLR) for evidence of a recent selective sweep is shown in purple. The location of significant signals ($P < 0.01$) calculated after simulations assuming neutrality (Supplementary text) is indicated with red circles. The locations of *bicycle* genes are indicated with orange rectangles.

(**B**) Distance from each *bicycle* gene to the closest significant selective sweep signal is shown as red histogram and dashed blue line indicates the median of this distribution.

(**C**) The median distance from each *bicycle* gene to the closest significant selective sweep signal (from **B**) is shown with dashed blue line and the values after 1000 permutations of sweep signals relative to gene locations are shown as grey histogram. The observed sweep signals are closer to *bicycle* genes than expected by chance.

**Table S1. $d_N/d_S$ test for significant positive selection on genes diverged between *H. cornu* and *H. hamamelidis*. Only genes estimated to have experienced significant positive or negative are included.**

| | $d_N/d_S$ | | |
|---|---|---|---|
| | <1 | >1 | Proportion >1 |
| non-*bicycle* | 5716 | 229 | 0.039 |
| *bicycle* | 10 | 58 | 0.853 |
| Proportion *bicycle* | 0.002 | 0.202 | |

**Table S2. McDonald-Kreitman alpha and gene statistics for different classes of genes overexpressed in *H. cornu* fundatrix salivary glands.**

| | CMH alpha | CMH alpha CI | non-synonymous substitutions ± SD | synonymous substitutions ± SD | Mean # CDS |
|---|---|---|---|---|---|
| *bicycle* | 0.33 | 0.242-0.408 | 8.0 ± 6.3 | 1.7 ± 1.5 | 683 |
| *bicycle* dnds > 1 | 0.45 | 0.362-0.533 | 9.8 ± 6.7 | 1.3 ± 1.2 | 689 |
| *bicycle* dnds significantly > 1 | 0.62 | 0.450-0.733 | 15.9 ± 8.8 | 1.1 ± 1.3 | 692 |
| *CWG* | 0.38 | 0.246-0.498 | 15.4 ± 7.8 | 6.3 ± 3.4 | 2337 |
| unclustered | 0.27 | 0.110-0.410 | 3.4 ± 4.0 | 1.4 ± 2.0 | 710 |
| Annotated with signal peptide | 0.42 | 0.243-0.557 | 3.8 ± 4.8 | 2.5 ± 2.2 | 1414 |
| No annotation with signal peptide | 0.34 | 0.262-0.404 | 8.7 ± 6.9 | 2.2 ± 2.3 | 848 |
| Annotated without signal peptide | 0.41 | 0.349-0.461 | 3.1 ± 4.8 | 3.2 ± 4.1 | 1750 |
| No annotation without signal peptide | 0.33 | 0.189-0.439 | 4.5 ± 4.8 | 1.4 ± 1.8 | 710 |
